# Imprints of tumor mutation burden on chromosomes and relation to cancer risk in humans: A pan-cancer analysis

**DOI:** 10.1101/2020.04.20.050989

**Authors:** Xin Li, D. Thirumalai

**Author notes:** Electronic mail. Electronic mail (D. Thirumalai).

## Abstract

Cancers, resulting in uncontrolled cell proliferation, are driven by accumulation of somatic mutations. Genome-wide sequencing has produced a catalogue of millions of somatic mutations, which contain the evolutionary history of the cancers. However, the connection between the mutation accumulation and disease development and risks is poorly understood. Here, we analyzed more than 1,200,000 mutations from 5,000 cancer patients with whole-exome sequencing, and discovered two novel signatures for 16 cancer types in The Cancer Genome Atlas (TCGA) database. A strong correlation between Tumor Mutation Burden (TMB) and the Patient Age at Diagnosis (PAD) is observed for cancers with low TMB (mean value less than 3 mutations per million base pairs) but is absent in cancers with high TMB. We also validate this finding using whole-genome sequencing data from more than 2,000 patients for 24 cancer types. Surprisingly, the differences in cancer risk between the sexes are also mainly driven by the disparity in mutation burden. The TMB variations, imprinted at the chromosome level, also reflect accumulation of mutation clusters within small chromosome segments in high TMB cancers. By analyzing the characteristics of mutations based on multi-region sequencing, we found that a combination of TMB and intratumor heterogeneity could a potential biomarker for predicting the patient survival and response to treatment.

## I. INTRODUCTION

Cancer is a complex disease caused by a gradual accumulation of somatic mutations, which occur through several distinct biological processes. Mutations could arise spontaneously during DNA replication^1^, through exogenous sources, such as ultraviolet radiation, smoking, virus inflammation and carcinogens^2^ or could be inherited^3^. During the past two decades, a vast amount of genomic data have been accumulated from patients with many cancers, thanks to developments in sequencing technology^4^. The distinct mutations contain the fingerprints of the cancer evolutionary history. The wealth of data have lead to the development of mathematical models^5–8^, which provide insights into biomarkers that are reasonable predictors of the efficacy of cancer therapies^9^. Our study is motivated by the following considerations:

### At what stage do most mutations accumulate?

The relation between mutation accumulation during cancer progression and disease risks in different tissue types and populations are not only poorly understood but also have resulted in contradictory conclusions. For instance, it has been suggested that half or more of the genetic mutations in cancer cells are acquired before tumor initiation^10,11^. In contrast, analyses of colorectal cancer tumors^12^ suggest that most of the mutations accumulate during the late stages of clonal expansion. In the cancer phylogenetic tree, representing tumor evolution, most mutations would appear in the trunk in the former case^10^ while the branches would contain most of the mutations if the latter scenario holds^12^. Therefore, simple measures that discriminate between the two scenarios would be invaluable either for effective drug screening (given more targetable trunk mutations) or prediction of resistance (presence of more branched mutations)^13^.

### The determinants of cancer risk

The number of mutations accumulated in cancer cells is likely related to great variations in cancer risks among different human tissue types^14^. According to the bad-luck theory^15^, cancer risk across tissues is strongly correlated with the total number of adult stem cell (ASC) divisions during a lifetime, with enhanced mutations accumulating in those that divide frequently. The implication is that some tissue types are at a greater disease risk than others. However, it is still unclear how mutation burden of cells is related to cancer risk among different tissues and genders.

### Biomarkers for patient survival and response to treatment

With advances in understanding the immune response against cancer, immunotherapy is fast becoming a very promising treatment for metastatic cancer patients^16–18^. However, a large fraction of patients do not respond to immunotherapy positively, and suffer from the inflammatory side effects, and likely miss out on other alternative treatments^19,20^. Therefore, finding biomarkers that could be used to separate patients who show response from those who do not^19,21^ is crucial. The tumor mutation burden (TMB), although is an important biomarker^22,23^ is not accurate, raising the need for other predictors^24^.

Motivated by the considerations described above, we developed computational methods to analyze the dynamics of accumulation of somatic mutations^15,25^ during cancer progression through a systematic pan-cancer study. We used the publicly available whole-exome sequencing (WES) data produced by The Cancer Genome Atlas (TCGA) project^26^, together with the whole-genome sequencing (WGS) data recently provided by the Pan-Cancer Analysis of Whole Genomes (PCAWG) Consortium^27^, which contain a large and well-characterized patient cohorts from multiple cancer types. In recent years, mutations in many cancers^28,29^ have been analyzed at the base pair level, to arrive at a number of signatures, which are combination of mutation types that drives different mutational process. A strong relation between the mutation load of two signatures and the patient age is found in many cancers (even in high TMB cancers)^28,30^. However, either only a small fraction (23%)^28,30^ of the genetic mutations are considered or it is found as data from all cancer types are mixed^27^. Here, we developed a simple coarse-grained measure of the total mutation load, instead of a small fraction, and a theoretical model that explains the distinct role of mutations in different cancers. We arrive at four major conclusions. (1) In the 16 cancers (with WES data), we discovered two distinct signatures for the accumulation of somatic mutations. There is a strong correlation between the tumor mutation burden (TMB) and the patient age as long as the overall mutation load is below a critical value of ~ 3 mutations/Mb. In sharp contrast, the correlation is weak for cancers with TMB that exceeds the critical value. Similar finding is observed for 24 cancers with WGS data. (2) Using a simple model, we show that if TMB is less than ~ 3 mutations/Mb, the majority of the mutations appear before the tumor initiation^10,11^. In the high TMB limit, most mutations accumulate during the tumor expansion stage, especially for some of the most lethal cancer types. (3) Although the mutation burden is one of the predominant factors that determines cancer risk, it only accounts for less than 50% of the differences in cancer risks among different tissues. Factors, such as the number of driver mutations required for each cancer, immunological and sex-hormone differences, also play a vital role in determining cancer risk. After excluding these factors, we show that the cancer risk disparity between the sexes is explained by the mutation load differences. (4) Our findings also show that a combination of TMB and intratumor heterogeneity could as serve as a two-dimensional biomarker for predicting patient survival and response to treatment.

## II. RESULTS

### Low Tumor Mutation Burden (TMB) strongly correlates with Patient Age at Diagnosis (PAD)

We define TMB as the total number of non-synonymous and synonymous single nucleotide variants (SNVs) per megabase (Mb) of the genome. First, we consider 16 cancer types with WES data taken from the TCGA database^26,31^ (see Table I in the Supplementary Information (SI)). Two mutation signatures emerge across these cancers. A strong positive correlation between TMB and PAD is found for cancers with low overall mutation load (TMB < 3 mutations/Mb). However, such a correlation is absent for cancers with high TMB. The distinct mutation signatures are a reflection of different underlying evolutionary dynamics among these distinct cancer types. We note parenthetically that the boundary between low and high TMB is somewhat arbitrary^32^. Interestingly, TMB ≈ 3 mutations/Mb, also provides a separation between cancers even at the chromosome level.

Two key inferences can be drawn from Figs. 1**a**-1**f**: (1) The linear behavior, for the six cancers in Fig. 1, between TMB and PAD reveals strong correlation with a Pearson correlation coefficient (*ρ*) varying between 0.66 to 0.93. The strong correlation (with a P value ≪ 0.001) is also evident directly from the violin plots shown in Fig. 1**g** (see also Fig. S1 in the SI), which present the full mutation distribution for the cancers as a function of PAD. Note that if the overall mean TMB is less than 3 mutations/Mb, as is the case for LAML, THCA, KIRC, and KIRP cancers, the Pearson correlation coefficient is ≥ 0.85. However, the correlation is weaker as the TMB approaches and exceeds 3 mutations/Mb, as is the case for GBM and KICH (see Figs. 1e and 1f). The TMB in these cancers becomes higher as the PAD increases, which is similar to the behavior in normal tissues (see especially Fig. 1 here and Fig. 2b in^25^). (2) The mutation rates calculated from the slope of the solid lines in Figs. 1**a**-1**f** vary from 13 (LAML) to 81 (KIRC) mutations per year for the whole genome. The range is in line with the mutation rate observed in normal tissues (40 to 65 mutations per year for normal colon, small intestine, liver and oesophageal epithelia cells)^12,25,33^. The calculated mutation rate, 13 mutations per year from Fig. 1**a**, for acute myeloid leukemia (LAML) coincides with the measure value (13±2) in previous studies^34^. Therefore, the cell mutation rate is approximately stable, and does not depend strongly on the tumor evolutionary process among the group of low TMB cancers (Fig. 1).

**FIG. 1:**
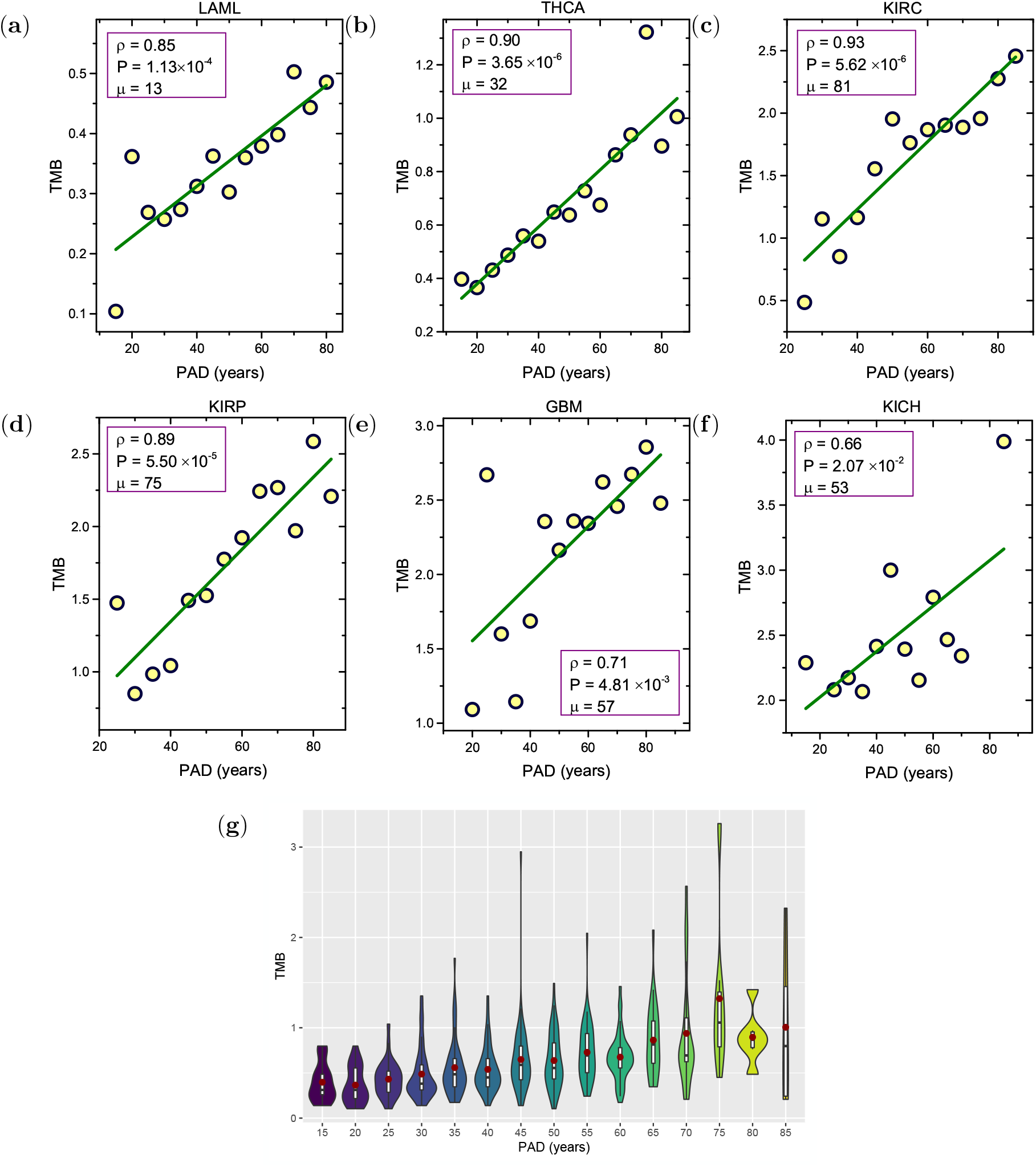
(a)-(e) The tumor mutation burden, TMB, (total number of non-synonymous and synonymous mutations per megabase) as a function of patient age at diagnosis (PAD). The mean value of TMB is used over a 5-year period. The green line shows the regression line. The Pearson correlation coefficient *ρ*, the significant P value from an F test and also the mutation rate *μ* are listed in the figures. The value of *μ* is derived from the slope of the green curve (0.0042 mutations/(Mb×year) for LAML in (a) for example, which leads to *μ* = 0.0042×3,000 ≈ 13 mutations/year for the whole genome). (g) Violin plots for the distribution of TMB as a function of PAD for THCA (see Fig. S1 for other cancers). Good correlation holds only if TMB is low. Data are taken from TCGA^26^.

**FIG. 2:**
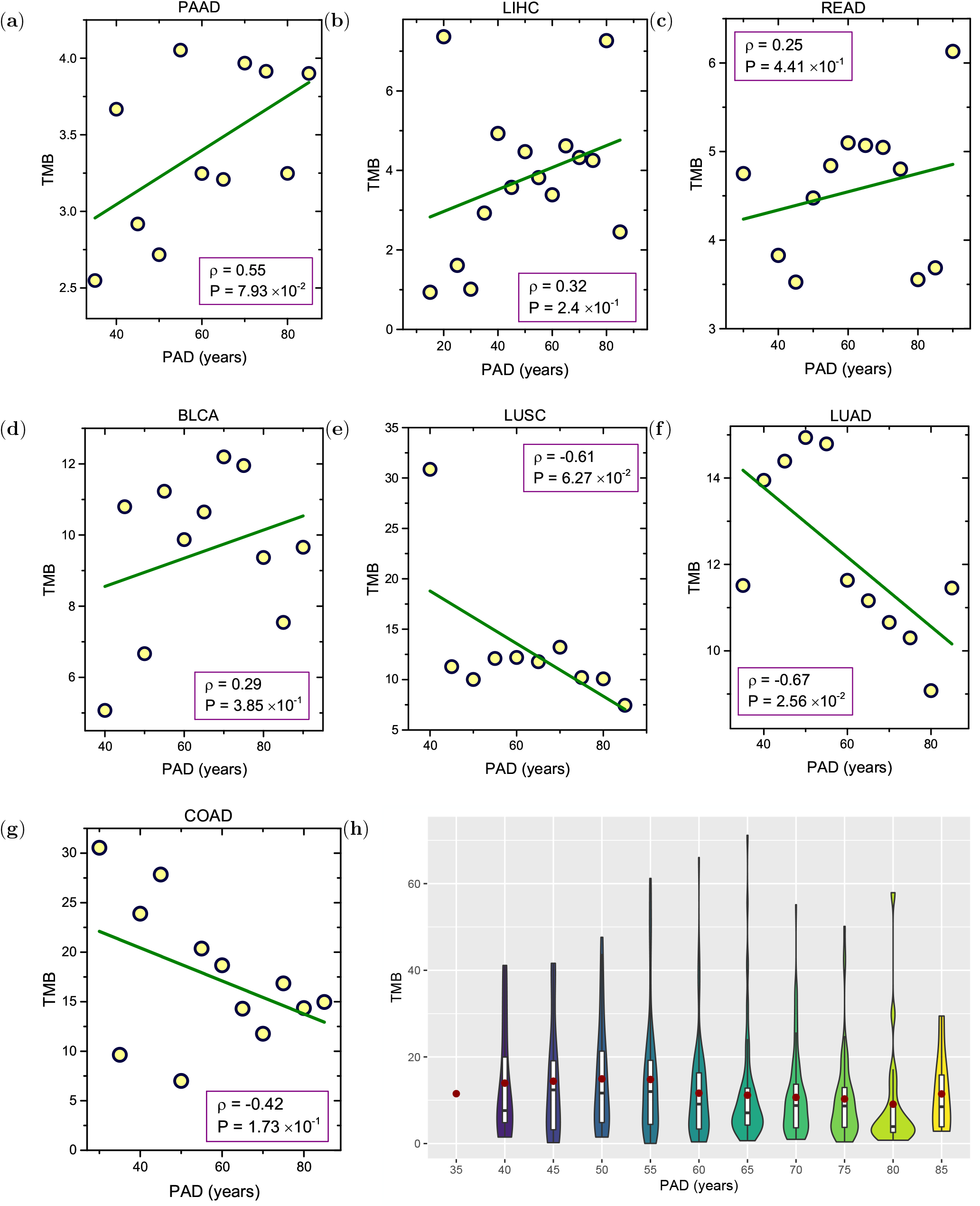
(a)-(g) Same as Fig.1 except the data are for cancers with high TMB. The green regression lines show a lack correlation between TMB and patient age. (h) Violin plots for the distribution of TMB as a function of PAD for LUAD (Fig. S2 in the SI provides data for other cancers).

### Large fraction of mutations accumulate before tumor initiation in low TMB cancers

The accumulation of mutations in cells of a tissue may be pictorially represented as a fish with the head, body and tail corresponding to the tissue development, self-renewal and tumor formation, respectively (see Fig. 1 in^10^). A prediction of the theory based on the “fish-like” picture is that the self-renewal stage (with a stable cell number) of a tissue is proportional to the patient age at tumor initiation. Therefore, a linear relation between the number of somatic mutations and the PAD must exist, assuming that a fixed time elapses from tumor initiation to diagnosis^10^. This important prediction is supported by our analyses for low TMB cancers, as shown in Fig. 1. Indeed, we can calculate the fraction of mutations accumulated in cells before tumor initiation once we know the average time it takes for tumor detection from the initiation time. Consider the thyroid carcinoma (THCA) as an example (Fig. 1b). The estimated latency period (*τ_L_*, from tumor initiation to diagnosis) is 5 to 20 years^35^. The number of mutations that accumulate in this time frame would be 150 to 600 (assuming an average accumulation rate of 30 mutations per year, see Fig. 1**b**). The median age of THCA patient at diagnosis is 46, and the number of accumulated mutations at that age would be around 2100, which is the product of TMB and the human genome length. By noting that the TMB is around 0.7/Mb at 46 years (Fig. 1**b**), and the human genome length 3,000Mb we arrive at 2100 mutations. Thus, for *τ_L_*=5 years only 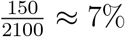 of the mutations occur during the period of tumor initiation till it is detected. This, implies that 93% and 71% of the mutations must have appeared before the initiation of the thyroid carcinoma for *τ_L_*=5 and 20 years, respectively. In this case, the time from tumor initiation to detection is usually much shorter than the age of patients at diagnosis. From a similar analysis (see Table II in the SI), we conclude that the majority of mutations appear before the tumor initiation for cancers shown in Fig. 1. This is consistent with the results of other studies^10,11,34^. Take LAML as an example. By sequencing the M3-LAML genome with a known initiating event versus the normal karyotype M1-LAML genome and the exomes of hematopoietic stem/progenitor cells (HSPCs) from healthy people^34^, it was found that most of the mutations in LAML genomes are indeed random events that occurred in HSPCs before they acquired the initiating mutation. We also observe in Fig. 1 that the correlation between the TMB and patient age becomes weaker as the overall TMB increases. As noted above, the *ρ* value decreases from 0.9 to 0.7 as the overall mean value (averaged over all patients for different ages) of TMB increases from 0.4 mutations/Mb to 3.0 mutations/Mb.

### High TMB and PAD are weakly correlated

As the overall TMB exceeds 3 mutations/Mb, there is no meaningful correlation between TMB and PAD (Fig. 2 and Fig. S2). Interestingly, the results in Fig. 2 show that the TMB even decreases in certain cancer types as the PAD increases (negative correlation if one exists at all), see Figs. 2**f**-2**h** as examples. This finding contradicts the conclusions reached in previous studies^10,36^. Clearly, the findings for cancers with high TMB cannot be explained by the fish-like model because of the absence of a linear relation between the TMB and PAD, which holds only for cancers with low TMB (Fig. 1).

### A simple model explains the TMB data

To understand the drastically different behavior between cancers with low and high TMBs in Figs. 1 and 2, we propose a simple theoretical model. It has been noted that the mutation rate can increase by a few orders of magnitude in tumor cells compared with normal cells caused by factors such as DNA mismatch repair deficiency or microsatellite instability^37,38^. As a consequence, a very high TMB would be detected in the tumor cells at the time of diagnosis. Therefore, the number of accumulated mutations (AM) in cells during the tumor formation stage could be comparable or even larger than the number during the tissue self-renewal stage. Let *μ*_1_ be the mutation rate during normal DNA replication process, and Δ*μ*_1_, the enhanced mutation rate (relative to *μ*_1_) in tumor cells. The value of Δ*μ*_1_ would be zero if the mutation rate does not change during tumor progression. In this case, the number *N_t_* of AM in tumor cells is given by *N_t_* = *μ*_1_*T* where *T* is the age of the patient in years. The linear relation holds if the TMB is low (Fig. 1). However, the mutation rate increases in high TMB cancers (Fig. 2). Then, the number of mutations *N_t_* is given by,

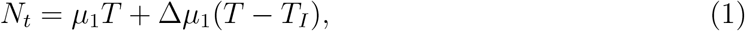

where the first term is the AM in the normal replication process, and the second term arises due to the accelerated mutations generated during the tumor formation stage. Note that *T_I_* corresponds to the time at tumor initiation. Because the latency period *T* − *T_I_* for tumors from the initiation till diagnosis is likely to be similar for patients with the same type of cancer^10^, the second term in Eq. (1) is likely to be a constant subject to only minor variations. If Δ*μ*_1_ ≫ *μ*_1_, then the first term in Eq. (1) is negligible, which leads to the weak dependence of TMB on the patient age as found in Fig. 2. Another potential mechanism for the finding in Fig. 2 could be that catastrophic events (such as chromoplexy, and chromothripsis) can lead to a burst of mutations that accumulate in tumors during a short period of time as observed in some cancers^39,40^.

It is instructive to calculate the fraction of accumulated mutations before tumor initiation for a cancer with high TMB. For the hepatocellular carcinoma (LIHC) the median age of patients at diagnosis is 61, so the number *N_I_* = *μ*_1_*T_I_* of AM before tumor initiation is less than *μ*_1_*T* ≈ 3000 (with rate *μ*_1_ ≈ 50 mutations/year^12,25^). From Fig. 2**b**, which shows that the TMB is ≈4 mutations/Mb at the same age and taking the genome length to be 3,000Mb, the total number of mutations accumulated at age 61 is about 12,000 (4 × 3000). Thus, the fraction of AM before tumor initiation should be less than 25%. We surmise that in cancers with high TMB (Fig. 2) most of the somatic mutations occur during the late stages of clonal expansion (see Table III in the SI). This conclusion agrees with the observations in colorectal cancers^12^, which allows us to conclude must be a cancer type with high TMB (> 3 mutations/Mb).

### Extrinsic Factors dominate mutation load in certain cancers

So far we have focused on somatic mutations arising from the intrinsic process due to DNA replication. The tissues of several cancers, such as head and neck, stomach, and skin, are constantly exposed to ultraviolet (UV) radiation and chemical carcinogens^41^. These factors directly increase the mutation rate (the “extrinsic origin”) of normal cells in the absence of DNA replication during the cell lifetime^42^. To account for extrinsic factors, we modify Eq. (1) by adding an additional term *N_ext_* = *μ_ext_T*. As long as *μ_ext_* ≫ Δ*μ*_1_, a linear relation between TMB and patient age holds (see Fig. 3(a)-(d) and Fig. S3 in the SI). In these cases, just as in Fig. 1, more mutations appear at the trunk of tumor phylogenetic trees types (see Table II in the SI and Fig. 3(d)). The strong linear correlation, with high Pearson correlation coefficient in Fig. 3(a)-(c), shows that in these cancers *μ_ext_* far exceeds *μ*_1_ and Δ*μ*_1_. It might be relatively easy for the immune system or drugs to clear cancer cells from patients with such cancers because a large number of mutations are shared among all the cancer cells^13^.

**FIG. 3:**
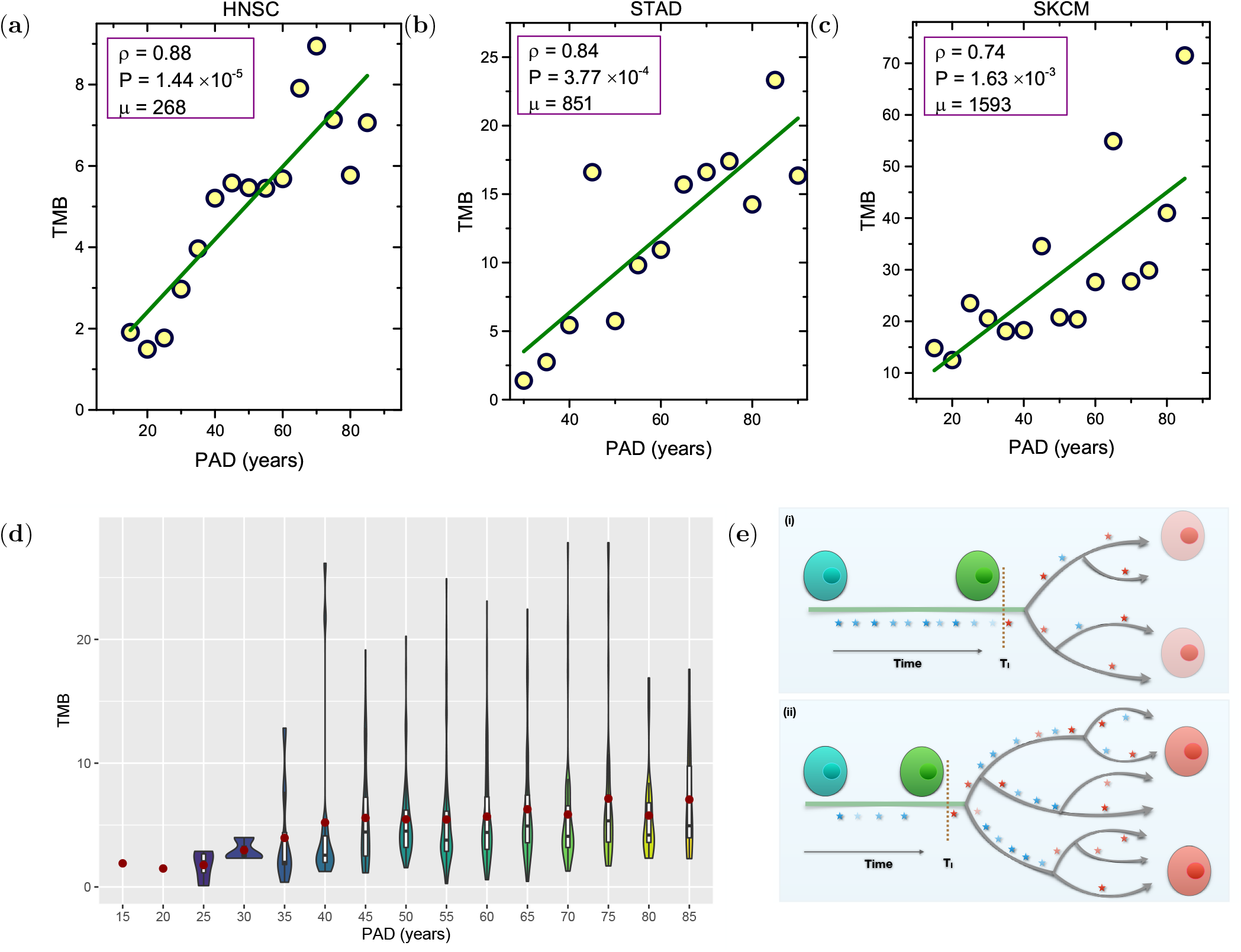
(a)-(c) TMB as a function of age for cancers strongly influenced by environments. The regression lines are in green. The Pearson correlation coefficient *ρ*, the significant P value, and the mutation rate *μ* (number of mutations per year for the whole genome) are shown in the figures. (d) Violin plots for the distribution of TMB as a function of PAD for HNSC (see Fig. S3 for other cancers). (e) Schematic of the mutation accumulation in cells during cancer progression. (i) corresponds to the case in Fig. 1 and Fig. 3(a)-(c), and (ii) shows the case in Fig. 2. Mutations are indicated by stars, and *T_I_* is the tumor initiation time. Normal cells are in blue/green colors without/with neutral mutations (blue stars) while cancer cells are in red color with driver mutations (red stars).

In contrast, the majority of the mutations are expected to be present in low frequencies and appear at the branches and leaves of the tumor phylogenetic tree (see Fig. 3(d)) for the cancers shown in Fig. 2. This results in immune surveillance escape and drug resistance^12^. In support of this finding, we found that the most deadly cancers (ones with the lowest 5-year survival rate), such as pancreatic cancer (8.5%), liver hepatocellular carcinoma (17.7%), and lung cancer (18.6%)^43^ (see Fig. S4 in the SI), appear in Fig. 2.

### Relation of TMB and PAD from whole-genome sequencing

Because a strong linear relation is observed (see Fig. S5 in the SI) between the number of SNVs for the whole genome and that for the whole exome of cancer patients, we expect similar correlation to be borne out by analyzing the WGS data. Using the recently available whole-genome sequencing data provided by the ICGC/TCGA Pan-Cancer Analysis of Whole Genomes (PCAWG) Consortium^27^, we performed a similar analysis for 24 cancer types (see Fig. 4). Due to the limited number of samples for each cancer type (see Table VII in the SI), we plotted all the patient data without binning into age groups. We found a strong correlation between TMB and PAD for cancer types with low mutation burden (see Fig. 4a-k and Table VII in the SI). However, such a correlation is absent for cancer types with high mutation burden (see Fig. 4l-x), which agrees well with our findings from the analysis of the WES data. For those cancers (Fig. 4r, s, x), under a high mutation burden but strongly influenced by extrinsic factors, a weak correlation is observed which might be due to the limited number of samples. A relatively strong correlation emerges for HSCC (head and neck cancer) compared to STAD (Stomach Adenocarcinoma) given a larger sample size for the former (see Fig. 4r and s). For MELA (Skin Melanoma), a very larger sample might be needed to observe the correlation due to substantial variations in the mutation burden. Therefore, the findings extracted from the WES data is robust, and is independent of the type of database we used.

**FIG. 4:**
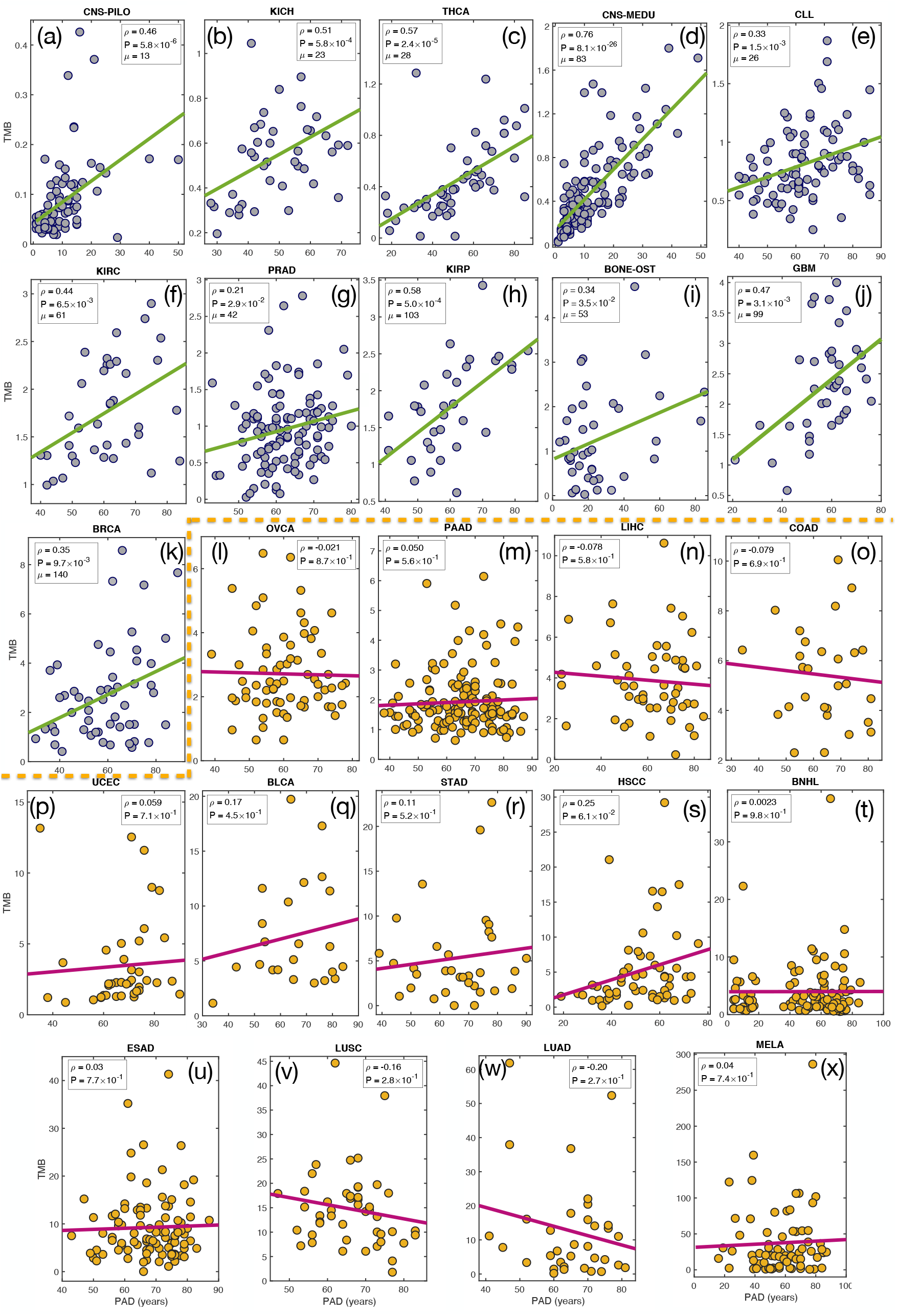
Tumor mutation burden, TMB, (total number of non-synonymous and synonymous mutations per megabase) as a function of patient age at diagnosis (PAD) for 24 types of cancers from the whole-genome sequencing^27^. Each filled circle represents one patient data. The green/magenta line shows the regression line. The Pearson correlation coefficient *ρ*, the significant P value from an F test and also the mutation rate *μ* (number of mutations per year for the whole genome in Figs. 4a-k) are listed in the figures. Good correlation holds only if TMB is low which confirms the findings in Figs. 1 and 2 obtained by analyzing the whole-exome sequencing data.

### Two universal patterns in cancers with low and high TMB

Our analyses above show two distinct signatures for TMB across many cancers. In order to elucidate if there are underlying universal behaviors in the cancers demarcated by the TMB, we first rescaled the TMB based on linear regression over the PAD for cancers listed in Figs. 1 and 3. Most interestingly, the data from the nine cancers fall on a linear curve, 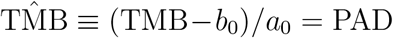 (see Fig. 5**a**), which clearly demonstrates a clock-like mutational process (with a constant mutation rate) in cancer evolution^30^. For the remaining cancers in Fig. 2, we rescaled the TMB only by the mean value *μ*_0_ because of the absence of a linear relationship. Surprisingly, the data from these cancers also fall into a straight line but with a nearly *zero* slope (see the Pearson correlation coefficient *ρ*, P-value and the green line in Fig. 5**b**), which supports the absence of *any* correlation between TMB and PAD for these cancer types.

**FIG. 5:**
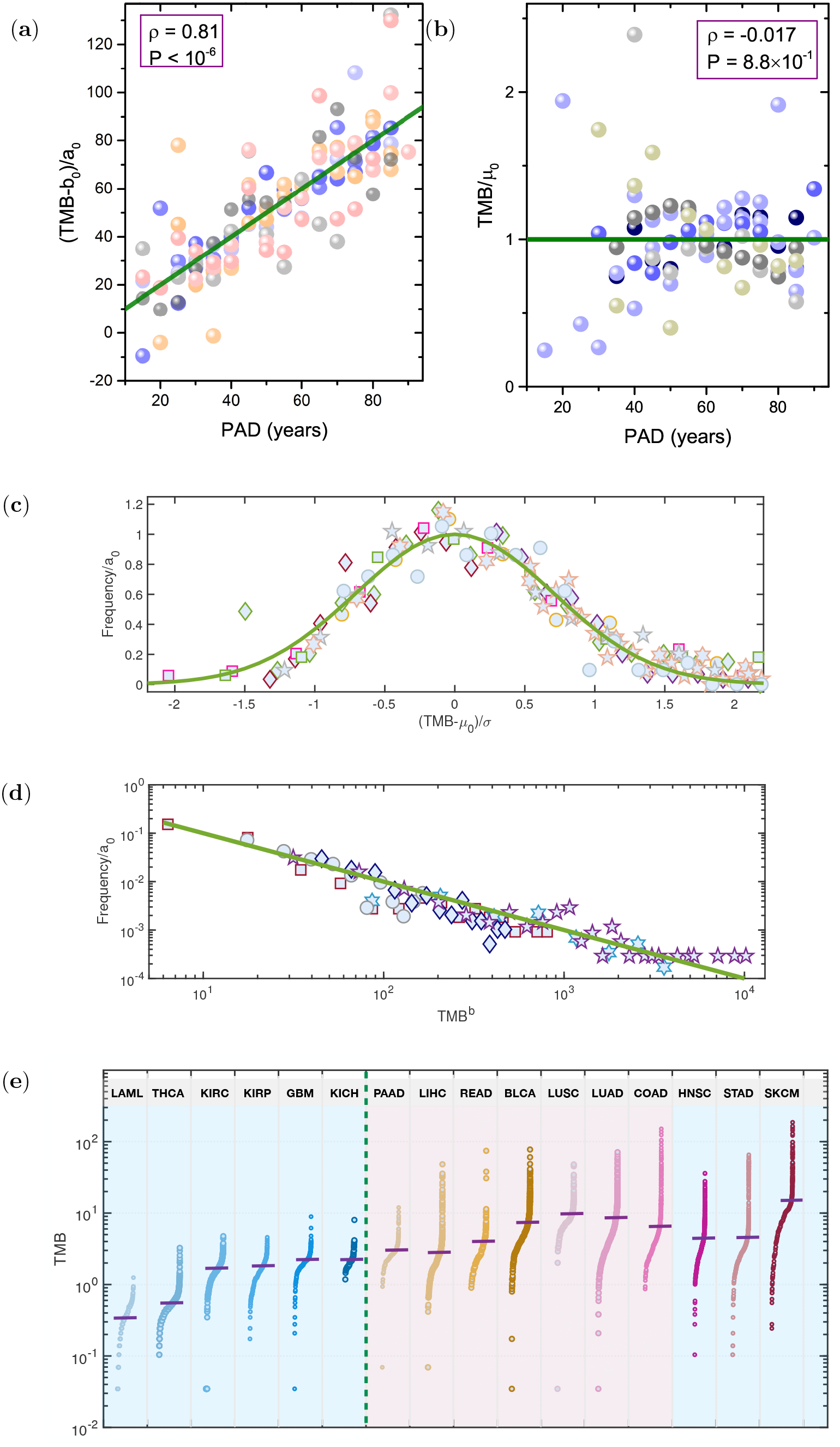
The rescaled TMB versus PAD for the 16 cancer types. (a) All nine types of cancer in Figs. 1 and 3 are displayed together in (a) after the TMB is rescaled by the slope (*a*_0_) and intercept (*b*_0_) of the corresponding linear regression lines. (b) The other seven types of cancers (Fig. 2) with TMB rescaled by the mean TMB value (*μ*_0_) of each cancer. The green line is described by *y* = *x* in (a) and *y* = 1 in (b). The Pearson correlation coefficient *ρ*, and the significant P value are given in the figure. (c) Rescaling the TMB distribution for each cancer type gives a Gaussian distribution (with the mean value *μ*_0_, variance *σ*, and coefficient *a*_0_) fits the data for cancers in Figs. 1 and 3. The rescaled TMB distributions for these cancers collapse into the standard normal distribution (see the green line). (d) Same procedure as in (c) leads to a fat tail power-law distribution (*P* (TMB) = *a*_0_ × TMB^*b*^) for data analyzed in Fig. 2. The green linear line has a slope value of −1. Each symbol represents one type of cancer. (e) The TMB across the 16 types of cancer. Each dot represents the TMB of a single cancer patient. The purple solid lines mark the median TMB value for each cancer. The green dashed line shows the boundary between cancers with low and high TMB defined in Figs. 1 and 2. The three cancer types discussed in Fig. 3 are listed at the end.

Next, we investigated the TMB distribution for each cancer type. Surprisingly, we found two universal TMB distribution patterns. A Gaussian distribution is found for all cancers analyzed in Figs. 1 and 3 (see Fig. S6 in the SI). Remarkably, after rescaling, the TMB distributions from the nine cancer types collapse onto a single master curve described by the normal distribution (see Fig. 5**c**). The universal TMB distribution found in Fig. 5**c** for the nine cancers is related to the Gaussian distribution of PAD (see Figs. S17 and S18 in the SI), and the linear relation between TMB and PAD (see Figs. 1 and 3). By simulating a population of patients with such properties, we obtain a Gaussian distribution for TMB (see Fig. S8 and the related discussion in the SI). We find it remarkable that for tumor evolution, which is a complex process, essentially a *single easily calculable parameter (TMB) from database* explains a vast amount of data for low TMB cancers.

In contrast, the distribution of cancers in Fig. 2 with large TMB is governed by a power-law (see Fig. S7 in the SI and Fig. 5**d**), except for PAAD (with TMB close to the critical value), and LUSC which have a Gaussian distribution. In these cancers, majority of the mutations accumulate after tumor initiation and the dynamics is likely to be highly complicated with many interrelated mutational processes contributing to clonal expansion. Similar signatures are found for cancers with low and high TMB, respectively, after we separate the patient data into two groups according to sex (see Figs. S9-S11 in the SI). The entirely different distributions of TMB found for the two groups of cancer show that cancers with high and low TMB follow distinct evolutionary processes. In low (high) TMB cancers, most of the mutations occur before (after) disease initiation.

Taken together, we discovered two distinct universal signatures for TMB across many cancers, which could be understood in terms of the mean/median TMB (~ 3 mutations/Mb, see the green dashed line in Fig. 5**e** as the boundary and Figs. S12 and S13 in the SI for further support). The linear relation between TMB and PAD, and the universal Gaussian distribution of TMB in the first group of cancers imply a clock-like mutation dynamics holds^30^. In contrast, the power-law distribution of TMB and lack of linear relation between TMB and PAD suggest a more complex mutation dynamics in the other group of cancers.

### Imprints of TMB in chromosomes

To illustrate the different mutation signatures between cancers with low and high TMB further, we investigate the mutation profile of each chromosome^44^ using LAML from Fig. 1 and LICH from Fig. 2 as two examples. The mutation frequency is much higher along each chromosome (at 10 Mb resolution) in the high TMB LIHC compared to the low TMB LAML (see Figs. 6(a) and (b)) except in the region around the centromere where a very low mutation frequency is observed in many chromosomes. From this coarse-grained description, a non-uniform distribution is observed clearly for the mutation profile in LIHC while it is rather uniform with only small deviations in LAML.

**FIG. 6:**
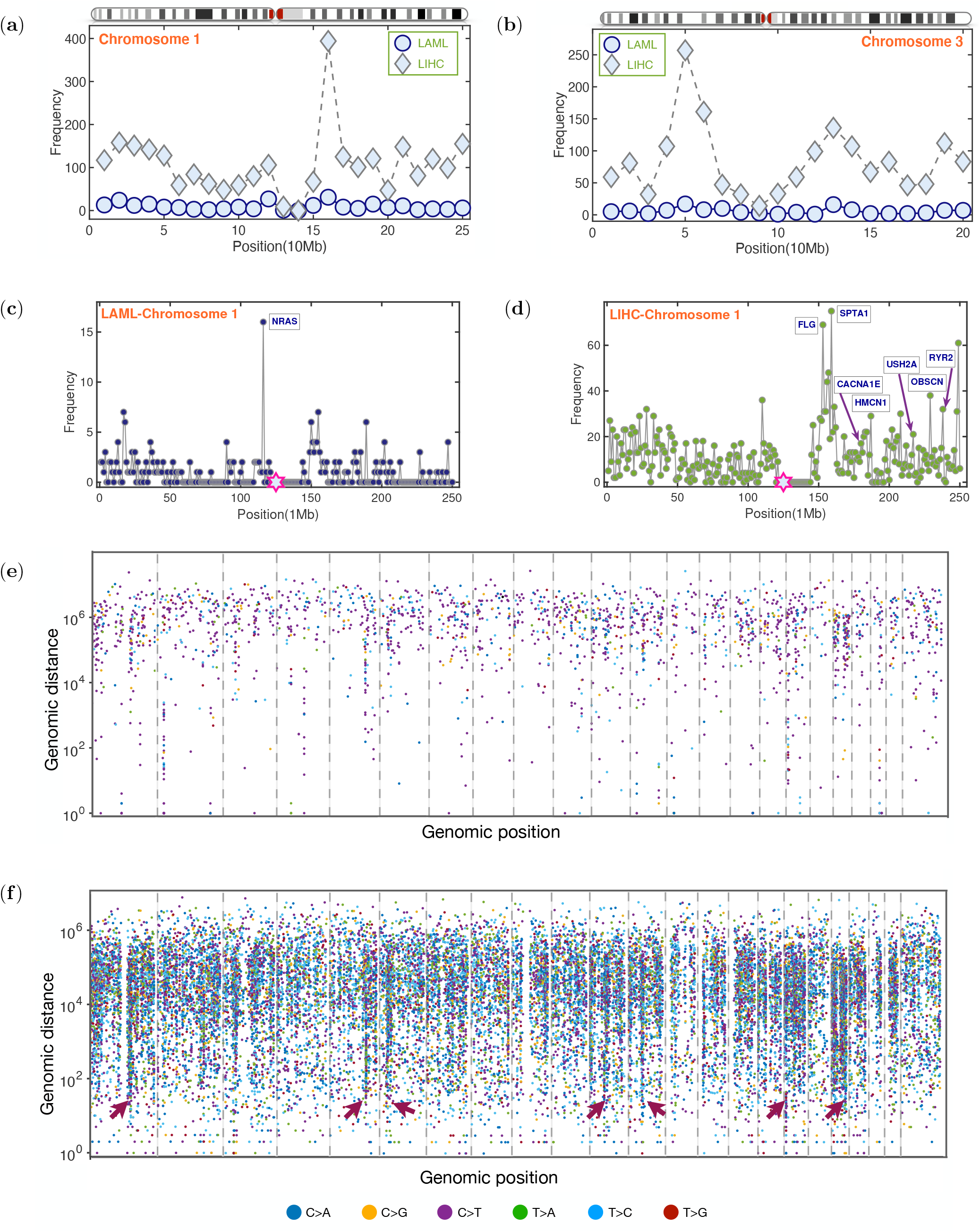
Tumor mutation profiles along chromosomes in LAML (low TMB) and LIHC (high TMB). (a) and (b): Number of mutations per 10 Mb across chromosomes 1 and 3 for LAML (197 patients) and LIHC (198 patients). The chromosome ideograms are also listed on top of each figure. (c) and (d): Number of mutations per 1 Mb along chromosomes 1 for LAML and LIHC. The genes (labelled in blue) mutated at a high frequency are also listed in the figures. The hexagram in magenta indicates the centromere location on each chromosome. (e) and (f): Rainfall plots for LAML and LIHC. The somatic mutations represented by small dots are ordered on the x axis in line with their positions in the human genome. The vertical value for each mutation is given by the genomic distance from the previous mutation. The colors of the dots represent different classes of base substitution, as demonstrated at the bottom of the figure. The red arrowheads indicate hypermutation regions. The 22 autosome and X chromosome are separated by the dashed lines.

Next, we consider a higher resolution (1Mb) mutation profiles (see Figs. 6**c**-6**d** and Figs. S13 and S14 in the SI) to capture more nuanced differences between these two cancers. Although the mutation frequency profiles are uniform in chromosomes 1 and 3 in LAML, occasionally, deviation from the mean value is detected (see the peak labeled with the name of a mutated gene in Fig. 6**c** and Fig. S14 in the SI) as a consequence of strong selection of driver mutations^45^. For the high TMB cancer, LIHC, a much more complex landscape is observed in the mutation profile. In addition to great variations of mutation frequency from one region to another, more highly mutated genes are detected and are often found to appear in a very close region along the genome (see Fig. 6**d** and Fig. S15 in the SI). This has been found in other high TMB cancers such as PAAD and LUAD^28^.

We then created the rainfall plot, which is frequently used to capture mutation clusters in chromosome^28^, to describe the different mutation characteristics for low and high TMB cancers (see Figs. 6**e** and 6**f**). The genomic distance between two nearby mutations is around 10^5^ - 10^6^*bp* for LAML. There is no clear hypermutation region for the 23 chromosomes (see Fig. 6**e** and Fig. S16) in this cancer type, which is consistent with the mutation profile illustrated in Fig 6**a**-6**c** and the simple Gaussian distribution found in Fig. 5**c** for low TMB cancers. In contrast, many hypermutation regions (with intermutation distance < 10^4^*bp*) are present in LIHC (see the red arrowheads in Fig. 6**f** and Fig. S17 for the detailed signatures in chromosome 1). Such a non-trivial mutation profile for high TMB cancers provides a hint for the power-law TMB distribution found in Fig. 5**d**. Similar profiles are also found for the kataegis patterns based on the WGS data (see Fig. S18 in the SI). These findings suggest that, on all scales, the evolutionary dynamics is much more complicated in high TMB cancers compared to low TMB cancers.

### Cancer risk and TMB

It is thought that the continuous accumulation of mutations drives the transformation of normal cells to cancerous cells. However, there is a controversy on the origin of the great variations in cancer risk among different tissues. A strong correlation is found between cancer risk in different tissues and the total number of adult stem cell divisions^15^. Therefore, the variations of cancer risk might be mainly determined by the mutation burden in the tissues, which seems to be consistent with the extreme variations of TMB observed in different cancer types (see Fig. 5**e**). We now assess the correlation between the cancer risk and the mutation burden in cancer patients directly by taking data for the 16 cancers considered above from the TCGA and SEER database (see Tables IV, V and the materials in the SI). The cancer risk for males and females vary greatly (see Table V in the SI), which we discuss further in the following sections. Here, we include the data for both the sexes. The median age for all cancer patients regardless of sex differences at diagnosis is around 66 years^14^. In order to rationalize the age disparity in different cancers for both the sexes (see Figs. S19-S20, and Table VI in the SI), we adjusted the value of TMB for both the sexes. We assumed that both the sexes accumulate mutations in a similar way (see Figs. S21-S23 and Section VIII in the SI for validation). We investigated the relation between cancer risk and TMB (see Fig. S24 (a) and (b)). The Pearson correlation coefficient *ρ* is 0.6 with a P value of 2.67 × 10^−4^ (*ρ* = 0.7 with *P* = 2.70 × 10^−5^) between cancer risk and TMB for 16 (15) cancers. Additional results are shown in Fig. S24 (c) and (d) without age-adjustment. A similar Pearson correlation coefficient (*ρ* = 0.66) is also found between cancer risk and mutation frequency across 41 cancers (using double-logarithmic coordinates with the data from a uniform sequencing pipeline) in a recent study^46^. Therefore, the cancer risk and TMB are correlated.

Because TMB accounts for less than 50% (≈ 0.7^2^) of differences in cancer risks among different tissues other factors must be taken into account to explain quantitatively cancer risks among all tissues. These include the driver mutation number^47^, immunological^48^ and sex-hormone differences^49^. Nevertheless, the mutation burden is one of the predominant factors for predicting cancer risks across different tissues. If we only compare cancer risks of the same type for both the sexes, excluding gender-specific cancers (with 16 types of cancers considered here as shown in Fig. 1-3), we might be able to exclude the influences of these factors on cancer incidence, which would allow us to focus only on TMB.

### TMB explains the role of sex in cancer risk

The risk ratio for the male to female changes from 0.39 (thyroid) to 28.73 (Kaposi sarcoma)^50^. For most types of cancer, men have a higher risk compared to women. Although breast and prostate cancers are gender-specific, driven by sex steroid hormones^49^, the enhanced risk in men is still a puzzle for many other non-gender-specific cancers. There are many potential factors leading to the disparity. These include but are not limited to genetic and expression variation^51^, sex hormones^52^, environmental exposures^53^.

We investigated the relation between cancer risk and the mutation burden for both the sexes for each cancer separately (see Figs. 7(a)-(b)). Surprisingly, the cancer risk can be explained solely by the TMB score in 13 out of the 16 cancers. Nine of them, which show high cancer risk, also have high TMB score (HNSC, KIRC, KIRP, LIHC, THCA, BLCA, LUAD, LUSC, and SKCM), and four of them show almost the same risk and have similar TMB (GBM, LAML, PAAD, and READ). The negative correlation between cancer risk and TMB in KICH in Fig. 7 might be because the sample size (66 in total) is small. In this case there is a large deviation in the median age of patients between men and women populations (see Fig. S20g). The lower risk, but higher TMB in women in STAD and COAD, could be caused by other factors such as the immune response and sex-hormone. Inflammation is frequently observed in these two cancer types, which results in the activation of the immune system^54^. Because the efficiency of the immune system declines faster in men^48^, and estrogen suppresses tumor growth in female patients with COAD^55^, they might decrease the cancer risk in women with STAD and COAD.

**FIG. 7:**
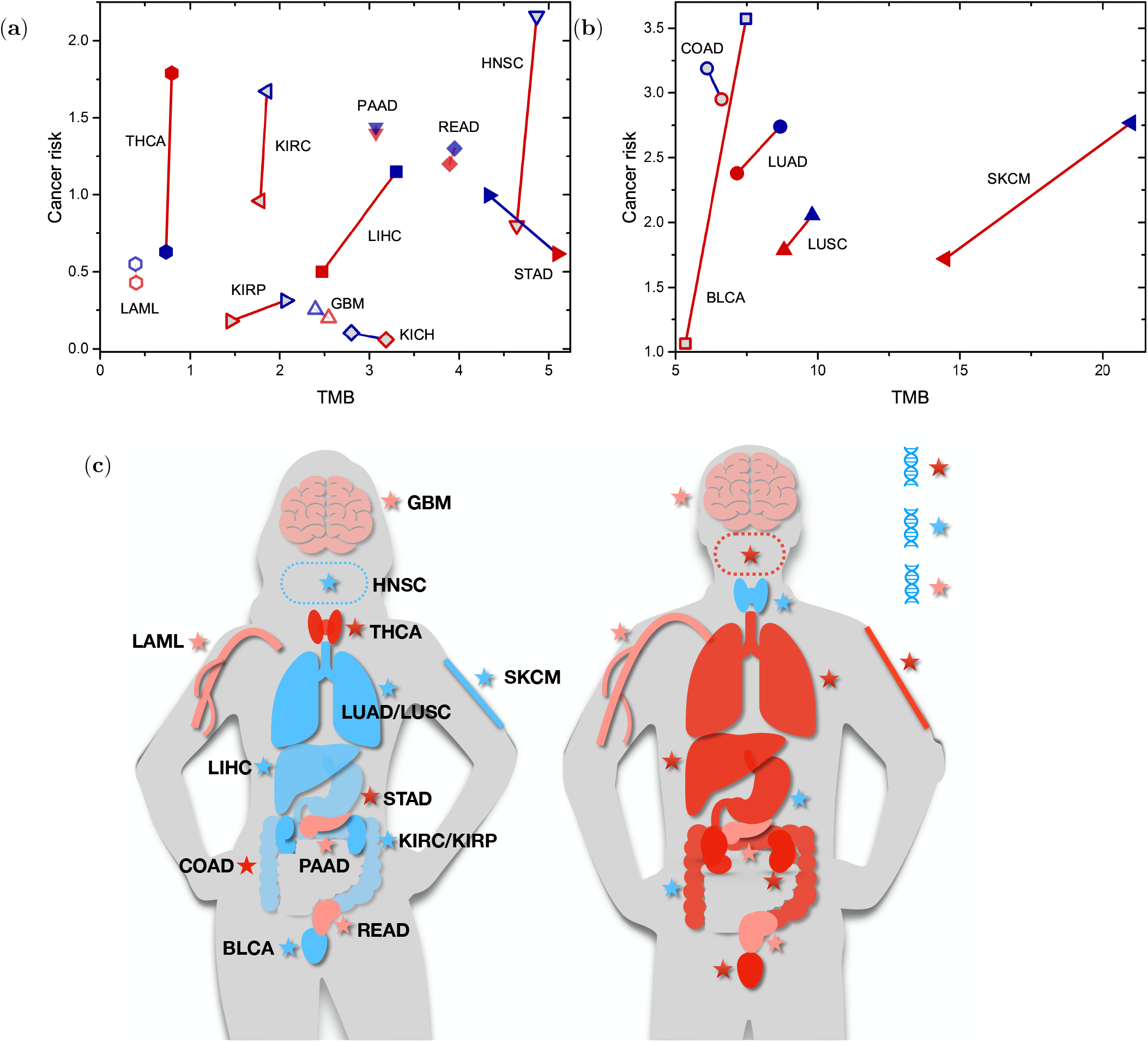
The relationship between lifetime cancer risk (%) and TMB for both the sexes. (a)-(b) The data are the same as in Fig. S22 (a) and (b) in the SI, while the solid lines (red for a positive, navy for a negative correlation) connect the data for both the sexes for the same cancer type. The data are separated into two groups for better visualization. Red (navy) symbols show the data for females (males). (c) Schematic of the relative cancer risks and the TMB for females and males. The organs in red, blue, and pink colors indicate a higher, lower and similar cancer risk (relative to the other gender), respectively. Similarly, the red, blue and pink stars show a higher, lower and similar TMB in the tissue accordingly. For clarity, we just use three colors to illustrate the relative values of these two quantities between the two genders. The exact values are shown in (a)-(b).

Taken together our analyses show that the risk disparity between men and women for the same type of cancer can be explained mainly through the mutation burden in the cancer cells (see the schematic illustration in Fig. 7(c)). However, other factors on cancers have to be considered to explain the risk among different tissues as shown in Fig. S24 and in previous studies^15,46^. The number (*N_d_*) of driver mutations required for cancer emergence is an important factor^47^. Because it takes a long time for a normal cell to accumulate driver mutations^56^, making the simplest assumption of independent driver mutation events, the cancer risk for tissues with a higher threshold of *N_d_* would be reduced. Consider THCA for instance. It is thought that a single driver mutation is required for the transition from a normal to a cancerous cell^47^. This might explain the relatively higher risk (see Fig. 7**a**) of THCA with low TMB compared to other cancers with lower risk but higher mutation burden, such as GBM which might require 6 driver mutations^47^. Although the mutation burden for SKCM is much higher than other cancers such as LUAD (HNSC), the large number of drivers (11) required for SKCM emergence could be the critical factor resulting in a similar risk as LUAD (HNSC), which might only require 4 (5) driver mutations (see Fig. 7**b**)^47^.

### Influence of TMB and intratumor heterogeneity (ITH) on patient survival

We have shown that high TMB is frequently related to high cancer risk for both the sexes for the same type of cancer. Although not desirable, high TMB could also elicit enhanced expression of a large number of neoantigens in cancer cells, which could be recognized by the T-cells for apoptosis and affect patient survival^57^. However, cancer cells have evolved different escape pathways, and could even disarm immune surveillance^58^. As we gain more knowledge about the mechanisms associated with these critical processes, immunotherapy may be modified to tame many metastatic cancers^16^ regardless of the TMB. At present, TMB is used as a potential biomarker for immunotherapy. Indeed, pembrolizumab has just been approved by FDA for the treatment of adult and pediatric patients with unresectable or metastatic tumor with high TMB based on cohorts with ten tumor types from the phase 2 KEYNOTE-158 study^59^. However, TMB alone is not a perfect predictor for patient response and survival^24^.

As an example, we investigated the influence of TMB, and patient age on patient response to immunotherapy (see Fig. S26 and the materials in the SI). Surprisingly, we found a much higher fraction of older patients, compared to younger ones, showed favorable response given a similar mutation burden level, which cannot be explained if only TMB is used as a predictor of the efficacy of treatment. We explained the data using a theoretical model based on the dynamics of mutation accumulation. We propose that older patients would have accumulated more clonal mutations under the same TMB level, which could be the underlying mechanism for the improved response.

To learn how the clonal characteristics of mutations influence patient survival further, we utilized the multi-region sequencing data which distinguishes the clonal from subclonal mutations better than single-region sequencing data^60^. We found that patients (lung adenocarcinoma), with a high clonal tumor mutation burden (cTMB, number of clonal mutations), show a better survival probability compared to those with a low cTMB (see Fig. 8a), thus supporting our conjecture above. However, the difference is not substantial which might due to the limited patient samples and short clinical trial duration. In addition, we also found that the patients with a low intratumor heterogeneity (ITH, defined based on the number of subclones^60^) have a better survival probability compared to those with a high ITH (see Fig. 8b).

**FIG. 8:**
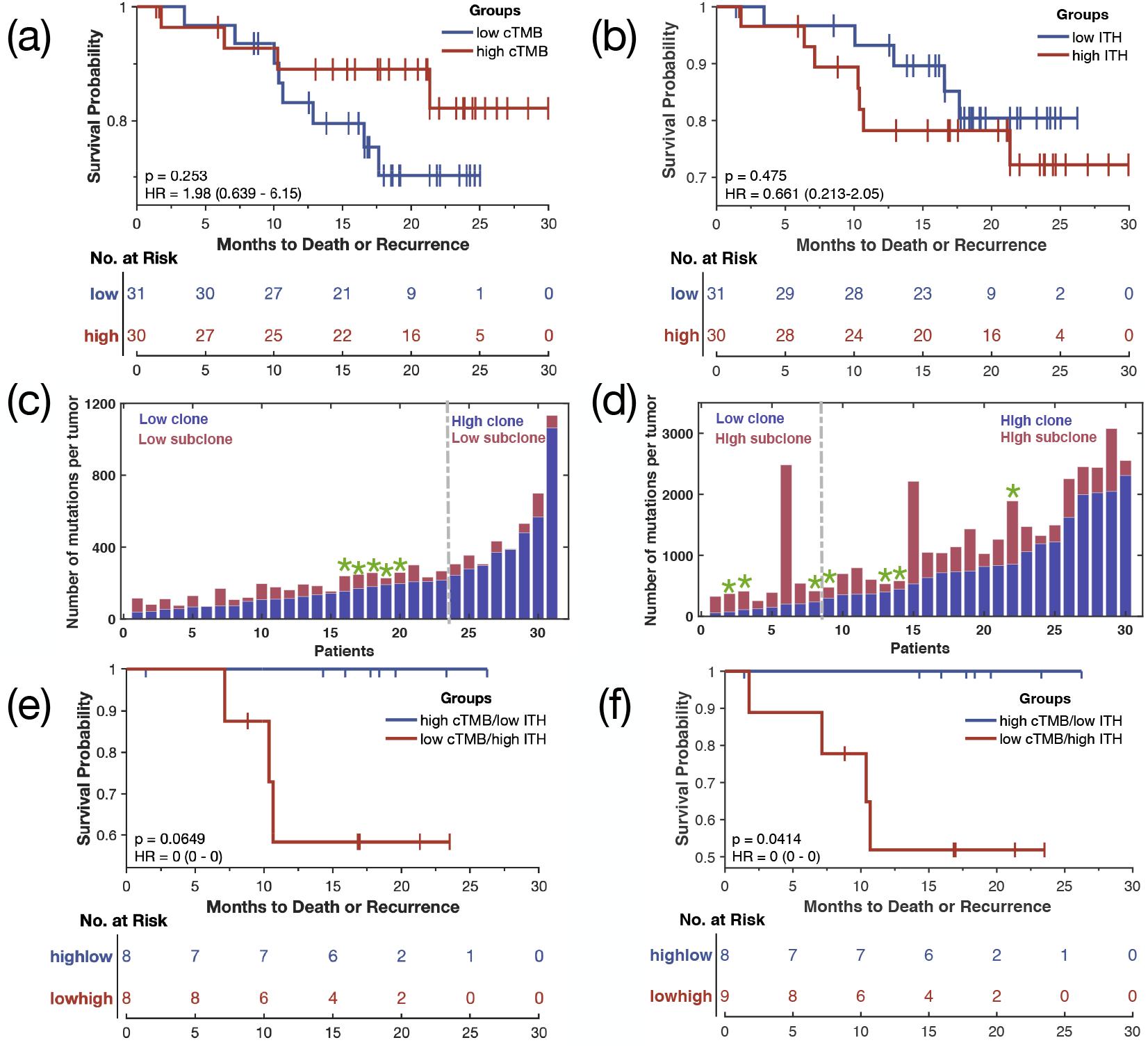
The influence of tumor mutation burden and intratumor heterogeneity on cancer (lung adenocarcinoma^60^) patient’s survival. (a) The relapse-free survival probability over a 30-month period for patients under high (above the median)/low clonal (below the median) tumor mutation burden (cTMB). (b) The relapse-free survival probability for patients under high (above the median)/low (below the median) intratumor heterogeneity (ITH, number of subclonal mutations). (c) The number of clonal (violet) and subclonal (red) mutations for patients with low ITH. (d) The number of clonal and subclonal mutations for patients with higher ITH. The dashed dotted lines in (c) and (d) separate the population into two categories based on cTMB. Green stars in (c) and (d) indicate recurrence or death. (e) The relapse-free survival probability over a 30-month period for patients under high cTMB and low ITH (low cTMB and high ITH) shown by the blue (red) line. (f) Same as Fig. 8(e) except for putting patient 9 (with similar number of clonal/subclonal mutations as patient 8) in Fig. 8(d) into the patient group under low cTMB and high ITH. The p-value and hazard ratio are shown in (a), (b), (e), and (f) using the log-rank test.

Based on these observations, instead of a single variable, we combined the cTMB and ITH and separated the patients into four groups (see Figs. 8c-d). We found that only the patient group with high cTMB but low ITH does not have recurrence or death events (see Fig. 8c). A significant higher survival probability is found in this group relative to the one with low cTMB and high ITH (see Figs. 8e-f). The reconstructed phylogenetic trees (see Figs. S27-28) show entirely different structures for the two categories of cancer patients. Therefore, instead of using one variable (TMB or ITH), a combination of these two factors could better predict the patient survival. These findings for lung adenocarcinoma show that the cancer biomarker landscape must at least be two dimensional, one representing ITH and the other TMB. Interestingly, a recent study^61^ has arrived at similar conclusions. By combining experiments on UVB-induced melanoma in mice and probing the immune response and analysis of TCGA melanoma patients, they discovered that the worst survival probability is found when TMB is low but ITH is high (Fig. 1E in^61^). Remarkably, this is also reflected in Fig. 8e. It does appear that both TMB and ITH, in principle measurable, could potentially predict survival probability more accurately, although high ITH is almost always detrimental.

## III. DISCUSSION

We investigated the evolutionary signatures focusing on the mutation burden in cancer patients, and their consequences on disease risks, variations between the sexes, and age related responses to treatment. By analyzing the data for 16 cancers from the WES and 24 cancers from the WGS, we found that two distinct scenarios for the accumulation of somatic mutations during cancer progression is determined by a single parameter. If the overall mutation burden is low (< 3 mutations/Mb), a strong positive correlation exists between the number of mutations and the patient age at diagnosis, which is absent for high TMB cancers. If the TMB is suitably rescaled then the distribution is a standard Gaussian for all cancers with low mutation load. In contrast, a power-law distribution of TMB is found for high TMB cancers. The mutation profiles along each chromosome are different for the two groups with high TMB cancers exhibiting a large number of mutations within small segments of the chromosomes. The variations of the mutation frequency along cancer genome could be caused by chromatin structure, replication timing and other factors^62^. We also found a few exceptions with high TMB where the positive linear relation between these two quantities is recovered for cancers such as HNSC, STAD, and SKCM. Interestingly, all these types of cancer are strongly influenced by environment^41^, which we explained by including the effect of extrinsic factors on mutations within the framework of a simple model.

For cancers that show a linear relation between the TMB and PAD, half or more mutations appear before tumor initiation^10^, which might be due to the relatively short time from tumor initiation to diagnosis relative to the duration of the tissue self-renewal. In sharp contrast, the majority of the mutations appear during the tumor expansion stage for other types of cancer, which exhibit weak correlation between the TMB and PAD, such as colorectal cancer^12^. In other words, the majority of the somatic mutations appear in the trunk of the cancer phylogenetic tree for cancers with low TMB while most of them would arise in the branches for high TMB cancers, as illustrated in Fig. 3**e**. This is consistent with the observations in other studies on acute myeloid leukemia, chronic lymphocytic leukemia, melanoma, and colorectal cancers.^10–12,34^. Our conclusion can be tested for other different cancer types with similar studies in the future.

The evolution of mutations in cancer cells, quantified using TMB, can serve as a valuable prognostic biomarker, allowing for a more informed treatment. Analysis of data from patients who underwent immunotherapy shows that better response in melanoma and non-small cell lung cancer is obtained in patients with high TMB^22^. In addition, recently FDA has approved pembrolizumab for several other cancer types with high TMB^59^. Nevertheless, TMB alone is not a perfect predictor for patient response^24^. We found that the fraction of older patients who show response to immunotherapy is much higher than younger ones under the same level of TMB. A further study leads us to conclude that two variables, TMB and ITH, could serve as a potential biomarker.

Apart from the genetic factors, the complex microenvironment of tumor cells is found to play a critical role during tumor initiation, progression and metastasis^63,64^. Some features of the tumor microenvironment (TME) is related to patient response to immune checkpoint blockade (ICB) therapy^65^. Therefore, a deep analysis of TME factors, such as the expression of PD-L1, the level and location of tumor-infiltrating CD8+ T cells and other features is likely to reveal additional biomarkers^65,66^. Finally, we can search for an optimized combination of biomarkers from somatic mutations and TME factors by data mining from TCGA and other resources^26^ in order to improve the prediction of patient survival and response.

The cancer risk for different tissues has been reported to be strongly correlated with the number of the stem cell divisions^15^ while a recent research claims that the cancer risk is strongly correlated with the mutation load^46^. One of our key finding is that the mutation burden is indeed correlated with the cancer risk. Interestingly, mutation burden alone explains the disparity of cancer risk found between the sexes after we exclude sex-hormone-related cancers. However, the present and previous studies^46^ with data from a uniform sequencing pipeline, show that the mutation load alone cannot account for all the differences in cancer risk among different tissues. Therefore, factors such as the number of driver mutations, immune responses, and sex hormones must be important in understanding the variations in the cancer risks among different tissues.

The signatures linking TMB and cancer risk is also imprinted in mutations at the base pair levels in the chromosomes. For instance, driver genes, such as DNMT3A and TET2 (see Fig. S14 in the SI), are significantly mutated in LAML cancer patients. Meanwhile, recent findings show that such mutations are also found in apparently healthy persons, and are associated with enhanced cancer risk^67^. Thus, there is a link between chromosomal abnormalities and TMB. Because TMB, a coarse-grained measure alone may not be sufficient as a biomarker, it should be combined with mutations at the chromosome level to assess cancer risk.

A deeper understanding of the evolutionary dynamics is the key to devise effective therapies for cancer. Although, we have investigated this process for more than 20 cancers covering a large population of patients, it would be valuable to assess if our findings are also applicable to other cancer types. Additional data will be needed to verify the correlation between cancer risk and the mutation load for KICH and other cancers between the genders. We have assumed that the mutation load found in cancer patients reflects the mutations accumulated in ancestral cells^46^. Therefore, we need more mutation profiles from normal and precancerous lesions to further validate the conclusions reached here^12,25,33^. In addition, the influences of other types of mutations such as copy number variations (no clear correlation is observed between the CNV burden and PAD from our preliminary analysis), and aneuploidy should also be considered in the future. Due to the complexity of cancer diseases and presence of many different risk factors simultaneously, it is valuable to evaluate one factor at a time (even if additivity principle does not hold), if at all possible, to better understand their roles in cancer progression.

## Methods

See the Supplementary Information for detailed methods, additional figures and data.

## Acknowledgments

We are grateful to Dr. S. Gail Eckhardt for her interest and useful comments, and Philipp M. Altrock, Shaon Chakrabarti, Anatoly B. Kolomeisky, and Sumit Sinha for pertinent discussions. This work is supported by the National Science Foundation (PHY 17-08128), and the Collie-Welch Chair through the Welch Foundation (F-0019).

## Author contributions

X.L. and D.T. conceived and designed the project, and co-wrote the paper. X.L. performed the research.

## Competing interests

We declare we have no competing interests.

## Notes

### Competing Interest Statement

The authors have declared no competing interest.

## REFERENCES

1 Thomas A Kunkel and Katarzyna Bebenek. DNA replication fidelity. Annual review of biochemistry, 69(1):497–529, 2000.

2 Errol C Friedberg, Lisa D McDaniel, and Roger A Schultz. The role of endogenous and exogenous DNA damage and mutagenesis. Current opinion in genetics & development, 14(1):5–10, 2004.

3 Eric R Fearon. Human cancer syndromes: clues to the origin and nature of cancer. Science, 278(5340):1043–1050, 1997.

4 Levi A Garraway and Eric S Lander. Lessons from the cancer genome. Cell, 153(1):17–37, 2013.

5 Philipp M Altrock, Lin L Liu, and Franziska Michor. The mathematics of cancer: integrating quantitative models. Nature Reviews Cancer, 15(12):730, 2015.

6 Abdul N Malmi-Kakkada, Xin Li, Himadri S Samanta, Sumit Sinha, and D. Thirumalai. Cell growth rate dictates the onset of glass to fluidlike transition and long time superdiffusion in an evolving cell colony. Physical Review X, 8(2):021025, 2018.

7 Martin A Nowak, Franziska Michor, and Yoh Iwasa. The linear process of somatic evolution. Proceedings of the national academy of sciences, 100(25):14966–14969, 2003.

8 Xin Li and D. Thirumalai. Share, but unequally: a plausible mechanism for emergence and maintenance of intratumour heterogeneity. Journal of the Royal Society Interface, 16(150):20180820, 2019.

9 Marta Luksza, Nadeem Riaz, Vladimir Makarov, Vinod P Balachandran, Matthew D Hellmann, Alexander Solovyov, Naiyer A Rizvi, Taha Merghoub, Arnold J Levine, Timothy A Chan, et al. A neoantigen fitness model predicts tumour response to checkpoint blockade immunotherapy. Nature, 551(7681):517, 2017.

10 Cristian Tomasetti, Bert Vogelstein, and Giovanni Parmigiani. Half or more of the somatic mutations in cancers of self-renewing tissues originate prior to tumor initiation. Proceedings of the National Academy of Sciences, 110(6):1999–2004, 2013.

11 Nicholas McGranahan and Charles Swanton. Clonal heterogeneity and tumor evolution: past, present, and the future. Cell, 168(4):613–628, 2017.

12 Sophie F Roerink, Nobuo Sasaki, Henry Lee-Six, Matthew D Young, Ludmil B Alexandrov, Sam Behjati, Thomas J Mitchell, Sebastian Grossmann, Howard Lightfoot, David A Egan, et al. Intra-tumour diversification in colorectal cancer at the single-cell level. Nature, 556(7702):457, 2018.

13 Nabil Amirouchene-Angelozzi, Charles Swanton, and Alberto Bardelli. Tumor evolution as a therapeutic target. Cancer discovery, 7(8):805–817, 2017.

14 N Howlader, AM Noone, M Krapcho, N Neyman, R Aminou, SF Altekruse, CL Kosary, J Ruhl, Z Tatalovich, H Cho, et al. SEER Cancer Statistics Review, 1975-2009 (vintage 2009 populations), National Cancer Institute. Bethesda, MD. *MD, USA*, 2012.

15 Cristian Tomasetti and Bert Vogelstein. Variation in cancer risk among tissues can be explained by the number of stem cell divisions. Science, 347(6217):78–81, 2015.

16 Ira Mellman, George Coukos, and Glenn Dranoff. Cancer immunotherapy comes of age. Nature, 480(7378):480, 2011.

17 Suzanne L Topalian, Charles G Drake, and Drew M Pardoll. Immune checkpoint blockade: a common denominator approach to cancer therapy. Cancer cell, 27(4):450–461, 2015.

18 Michael A Postow, Robert Sidlow, and Matthew D Hellmann. Immune-related adverse events associated with immune checkpoint blockade. New England Journal of Medicine, 378(2):158–168, 2018.

19 Suzanne L Topalian, Janis M Taube, Robert A Anders, and Drew M Pardoll. Mechanism-driven biomarkers to guide immune checkpoint blockade in cancer therapy. Nature Reviews Cancer, 16(5):275, 2016.

20 Padmanee Sharma, Siwen Hu-Lieskovan, Jennifer A Wargo, and Antoni Ribas. Primary, adaptive, and acquired resistance to cancer immunotherapy. Cell, 168(4):707–723, 2017.

21 Padmanee Sharma and James P Allison. The future of immune checkpoint therapy. Science, 348(6230):56–61, 2015.

22 Aaron M Goodman, Shumei Kato, Lyudmila Bazhenova, Sandip P Patel, Garrett M Frampton, Vincent Miller, Philip J Stephens, Gregory A Daniels, and Razelle Kurzrock. Tumor mutational burden as an independent predictor of response to immunotherapy in diverse cancers. Molecular cancer therapeutics, 16(11):2598–2608, 2017.

23 Matthew D Hellmann, Tavi Nathanson, Hira Rizvi, Benjamin C Creelan, Francisco Sanchez-Vega, Arun Ahuja, Ai Ni, Jacki B Novik, Levi MB Mangarin, Mohsen Abu-Akeel, et al. Genomic features of response to combination immunotherapy in patients with advanced non-small-cell lung cancer. Cancer cell, 33(5):843–852, 2018.

24 Denis L Jardim, Aaron Goodman, Debora de Melo Gagliato, and Razelle Kurzrock. The challenges of tumor mutational burden as an immunotherapy biomarker. Cancer Cell, 39, 2021.

25 Francis Blokzijl, Joep De Ligt, Myrthe Jager, Valentina Sasselli, Sophie Roerink, Nobuo Sasaki, Meritxell Huch, Sander Boymans, Ewart Kuijk, Pjotr Prins, et al. Tissue-specific mutation accumulation in human adult stem cells during life. Nature, 538(7624):260, 2016.

26 John N Weinstein, Eric A Collisson, Gordon B Mills, Kenna R Mills Shaw, Brad A Ozenberger, Kyle Ellrott, Ilya Shmulevich, Chris Sander, Joshua M Stuart, Cancer Genome Atlas Research Network, et al. The cancer genome atlas pan-cancer analysis project. Nature genetics, 45(10):1113, 2013.

27 The ICGC/TCGA Pan-Cancer Analysis of Whole Genomes Consortium. Pan-cancer analysis of whole genomes. Nature, 578(7793):82, 2020.

28 Ludmil B Alexandrov, Serena Nik-Zainal, David C Wedge, Samuel AJR Aparicio, Sam Behjati, Andrew V Biankin, Graham R Bignell, Niccolo Bolli, Ake Borg, Anne-Lise Børresen-Dale, et al. Signatures of mutational processes in human cancer. Nature, 500(7463):415–421, 2013.

29 Ludmil B Alexandrov, Serena Nik-Zainal, David C Wedge, Peter J Campbell, and Michael R Stratton. Deciphering signatures of mutational processes operative in human cancer. Cell reports, 3(1):246–259, 2013.

30 Ludmil B Alexandrov, Philip H Jones, David C Wedge, Julian E Sale, Peter J Campbell, Serena Nik-Zainal, and Michael R Stratton. Clock-like mutational processes in human somatic cells. Nature genetics, 47(12):1402, 2015.

31 Mehmet Kemal Samur. RTCGAToolbox: a new tool for exporting TCGA Firehose data. PloS one, 9(9):e106397, 2014.

32 Timothy A Chan, Mark Yarchoan, Elizabeth Jaffee, Charles Swanton, Sergio A Quezada, Albrecht Stenzinger, and Solange Peters. Development of tumor mutation burden as an immunotherapy biomarker: utility for the oncology clinic. Annals of Oncology, 30(1):44–56, 2019.

33 Akira Yokoyama, Nobuyuki Kakiuchi, Tetsuichi Yoshizato, Yasuhito Nannya, Hiromichi Suzuki, Yasuhide Takeuchi, Yusuke Shiozawa, Yusuke Sato, Kosuke Aoki, Soo Ki Kim, et al. Age-related remodelling of oesophageal epithelia by mutated cancer drivers. Nature, 565(7739):312, 2019.

34 John S Welch, Timothy J Ley, Daniel C Link, Christopher A Miller, David E Larson, Daniel C Koboldt, Lukas D Wartman, Tamara L Lamprecht, Fulu Liu, Jun Xia, et al. The origin and evolution of mutations in acute myeloid leukemia. Cell, 150(2):264–278, 2012.

35 AF Tsyb, EM Parshkov, VV Shakhtarin, VF Stepanenko, VF Skvortsov, and IV Chebotareva. Thyroid cancer in children and adolescents of Bryansk and Kaluga Regions. Age, 4(10-14):15–19, 1996.

36 Dmitriy I Podolskiy, Alexei V Lobanov, Gregory V Kryukov, and Vadim N Gladyshev. Analysis of cancer genomes reveals basic features of human aging and its role in cancer development. Nature communications, 7:12157, 2016.

37 Christoph Lengauer, Kenneth W Kinzler, and Bert Vogelstein. Genetic instabilities in human cancers. Nature, 396(6712):643, 1998.

38 Lawrence A Loeb. A mutator phenotype in cancer. Cancer research, 61(8):3230–3239, 2001.

39 Sylvan C Baca, Davide Prandi, Michael S Lawrence, Juan Miguel Mosquera, Alessandro Romanel, Yotam Drier, Kyung Park, Naoki Kitabayashi, Theresa Y MacDonald, Mahmoud Ghandi, et al. Punctuated evolution of prostate cancer genomes. Cell, 153(3):666–677, 2013.

40 Andrea Sottoriva, Haeyoun Kang, Zhicheng Ma, Trevor A Graham, Matthew P Salomon, Junsong Zhao, Paul Marjoram, Kimberly Siegmund, Michael F Press, Darryl Shibata, et al. A big bang model of human colorectal tumor growth. Nature genetics, 47(3):209, 2015.

41 Cristian Tomasetti, Lu Li, and Bert Vogelstein. Stem cell divisions, somatic mutations, cancer etiology, and cancer prevention. Science, 355(6331):1330–1334, 2017.

42 Song Wu, Scott Powers, Wei Zhu, and Yusuf A Hannun. Substantial contribution of extrinsic risk factors to cancer development. Nature, 529(7584):43, 2016.

43 AM Noone, N Howlader, M Krapcho, D Miller, A Brest, M Yu, J Ruhl, Z Tatalovich, A Mariotto, DR Lewis, et al. SEER Cancer Statistics Review, 1975-2015, National Cancer Institute. Bethesda, MD, 2018.

44 Graham R Bignell, Chris D Greenman, Helen Davies, Adam P Butler, Sarah Edkins, Jenny M Andrews, Gemma Buck, Lina Chen, David Beare, Calli Latimer, et al. Signatures of mutation and selection in the cancer genome. Nature, 463(7283):893–898, 2010.

45 Klaus H Metzeler, Tobias Herold, Maja Rothenberg-Thurley, Susanne Amler, Maria C Sauerland, Dennis Görlich, Stephanie Schneider, Nikola P Konstandin, Annika Dufour, Kathrin Bräundl, et al. Spectrum and prognostic relevance of driver gene mutations in acute myeloid leukemia. Blood, 128(5):686–698, 2016.

46 Dapeng Hao, Li Wang, and Li-jun Di. Distinct mutation accumulation rates among tissues determine the variation in cancer risk. Scientific Reports, 6:19458, 2016.

47 Iñigo Martincorena, Keiran M Raine, Moritz Gerstung, Kevin J Dawson, Kerstin Haase, Peter Van Loo, Helen Davies, Michael R Stratton, and Peter J Campbell. Universal patterns of selection in cancer and somatic tissues. Cell, 171(5):1029–1041, 2017.

48 Sam Palmer, Luca Albergante, Clare C Blackburn, and TJ Newman. Thymic involution and rising disease incidence with age. Proceedings of the National Academy of Sciences, 115(8):1883–1888, 2018.

49 Gail P Risbridger, Ian D Davis, Stephen N Birrell, and Wayne D Tilley. Breast and prostate cancer: more similar than different. Nature Reviews Cancer, 10(3):nrc2795, 2010.

50 Michael B Cook, Sanford M Dawsey, Neal D Freedman, Peter D Inskip, Sara M Wichner, Sabah M Quraishi, Susan S Devesa, and Katherine A McGlynn. Sex disparities in cancer incidence by period and age. Cancer Epidemiology and Prevention Biomarkers, 18(4):1174–1182, 2009.

51 Yuan Yuan, Lingxiang Liu, Hu Chen, Yumeng Wang, Yanxun Xu, Huzhang Mao, Jun Li, Gordon B Mills, Yongqian Shu, Liang Li, et al. Comprehensive characterization of molecular differences in cancer between male and female patients. Cancer cell, 29(5):711–722, 2016.

52 Shelia Hoar Zahm and Joseph F Fraumeni Jr. Racial, ethnic, and gender variations in cancer risk: considerations for future epidemiologic research. Environmental health perspectives, 103(Suppl 8):283, 1995.

53 Frederica P Perera. Environment and cancer: who are susceptible? Science, 278(5340):1068–1073, 1997.

54 Lisa M Coussens and Zena Werb. Inflammation and cancer. Nature, 420(6917):860, 2002.

55 M Tevfik Dorak and Ebru Karpuzoglu. Gender differences in cancer susceptibility: an inadequately addressed issue. Frontiers in genetics, 3:268, 2012.

56 Ivana Bozic, Tibor Antal, Hisashi Ohtsuki, Hannah Carter, Dewey Kim, Sining Chen, Rachel Karchin, Kenneth W Kinzler, Bert Vogelstein, and Martin A Nowak. Accumulation of driver and passenger mutations during tumor progression. Proceedings of the National Academy of Sciences, 107(43):18545–18550, 2010.

57 Ton N Schumacher and Robert D Schreiber. Neoantigens in cancer immunotherapy. Science, 348(6230):69–74, 2015.

58 Ryungsa Kim, Manabu Emi, and Kazuaki Tanabe. Cancer immunoediting from immune surveillance to immune escape. Immunology, 121(1):1–14, 2007.

59 Aurélien Marabelle, Marwan Fakih, Juanita Lopez, Manisha Shah, Ronnie Shapira-Frommer, Kazuhiko Nakagawa, Hyun Cheol Chung, Hedy L Kindler, Jose A Lopez-Martin, Wilson H Miller Jr, et al. Association of tumour mutational burden with outcomes in patients with advanced solid tumours treated with pembrolizumab: prospective biomarker analysis of the multicohort, open-label, phase 2 keynote-158 study. The Lancet Oncology, 21(10):1353–1365, 2020.

60 Mariam Jamal-Hanjani, Gareth A Wilson, Nicholas McGranahan, Nicolai J Birkbak, Thomas BK Watkins, Selvaraju Veeriah, Seema Shafi, Diana H Johnson, Richard Mitter, Rachel Rosenthal, et al. Tracking the evolution of non–small-cell lung cancer. New England Journal of Medicine, 376(22):2109–2121, 2017.

61 Yochai Wolf, Osnat Bartok, Sushant Patkar, Gitit Bar Eli, Sapir Cohen, Kevin Litchfield, Ronen Levy, Alejandro Jiménez-Sánchez, Sophie Trabish, Joo Sang Lee, et al. UVB-induced tumor heterogeneity diminishes immune response in melanoma. Cell, 179(1):219–235, 2019.

62 Paz Polak, Rosa Karlić, Amnon Koren, Robert Thurman, Richard Sandstrom, Michael S Lawrence, Alex Reynolds, Eric Rynes, Kristian Vlahoviček, John A Stamatoyannopoulos, et al. Cell-of-origin chromatin organization shapes the mutational landscape of cancer. Nature, 518(7539):360–364, 2015.

63 TL Whiteside. The tumor microenvironment and its role in promoting tumor growth. Oncogene, 27(45):5904–5912, 2008.

64 Daniela F Quail and Johanna A Joyce. Microenvironmental regulation of tumor progression and metastasis. Nature medicine, 19(11):1423–1437, 2013.

65 Nadeem Riaz, Jonathan J Havel, Vladimir Makarov, Alexis Desrichard, Walter J Urba, Jennifer S Sims, F Stephen Hodi, Salvador Martín-Algarra, Rajarsi Mandal, William H Sharfman, et al. Tumor and microenvironment evolution during immunotherapy with nivolumab. Cell, 171(4):934–949, 2017.

66 Mikhail Binnewies, Edward W Roberts, Kelly Kersten, Vincent Chan, Douglas F Fearon, Miriam Merad, Lisa M Coussens, Dmitry I Gabrilovich, Suzanne Ostrand-Rosenberg, Catherine C Hedrick, et al. Understanding the tumor immune microenvironment (TIME) for effective therapy. Nature medicine, 24(5):541–550, 2018.

67 Giulio Genovese, Anna K Kähler, Robert E Handsaker, Johan Lindberg, Samuel A Rose, Samuel F Bakhoum, Kimberly Chambert, Eran Mick, Benjamin M Neale, Menachem Fromer, et al. Clonal hematopoiesis and blood-cancer risk inferred from blood DNA sequence. New England Journal of Medicine, 371(26):2477–2487, 2014.

