## Supplementary Information for "Imprints of tumor mutation burden on chromosomes and relation to cancer risk in humans: A pan-cancer analysis"

(Dated: 26 February 2021)

In the Supplementary Information (SI) we provide in depth analyses of substantial additional data, accompanied by figures that are not only relevant to the results in the main text but also fortify the essential conclusions reached in our study.

The Cancer Genome Atlas (TCGA) research network has collected and analyzed the genomic, protein and epigenetic profiles of more than 11,000 specimens from more than 30 cancer types[1,2]. We focus on the link between tumor mutation burden (TMB) and cancer risk as well as discernible differences between the sexes based on TMB. Hence, we choose 16 major cancers, and excluded data from cancers with small samples and those that are gender specific. We take the mutation data for each cancer from the Firehose pipeline developed by the Broad Institute (http://firebrowse.org). The MuTect algorithm[3–5] is used by the platform to detect the mutation profile of cancer patients. The patient age and clinical information are obtained through the RTCGAToolbox[6]. The data with patient age and mutation information that are missing in the database are excluded from our analysis. The number of patients for each cancer type is listed in Table I.

Some details of the bioinformatics pipeline for the whole-exome sequencing (WES) data: the standard parameters in the MuTect algorithm are used to detect the single nucleotide variants (SNVs) which can be found in the original reference[5]. Bayesian statistical analysis of bases and their qualities in the tumor and normal Binary Alignment Maps (BAMs) is used by MuTect to identify candidate SNVs at a given genomic locus. A minimum of 14 reads in the tumor and 8 in the normal tissue is required for mutation calling. The minimal allelic fraction cutoff is 0.1. A panel of normal samples as a filter to reduce false positives and miscalled germ-line events. The public dbSNP database is used to distinguish two cases: (i) sites which are known to be variant in the population and (ii) all other sites. More details about the method can be found in the original article[5].

The whole-genome sequencing (WGS) data are taken from the ICGC/TCGA Whole Genomes Consortium which collects the genome data from 2834 patients across 38 tumor types[7]. Four different algorithms: MuTect, RADIA, Strelka, and SomaticSniper are used to detect the SNVs (at least called by 3 out of the 4 algorithms or called by SomaticSniper and one other algorithm). The minimal allelic fraction cutoff is 0.1. The SNVs are further filtered by the database dbSNP. More details about the SNV calls can be found in the SI of the reference[7].

### I. DISTRIBUTION OF TUMOR MUTATION BURDEN (TMB) FOR THE SIXTEEN TYPES OF CANCER FOR DIFFERENT AGE GROUPS

The distribution of TMB for distinct age groups (five years interval for each group) for the 16 cancers are shown in Figs. S1-S3 using the violin plot representation. The mean value of TMB for each group of patients is indicated by the red dot in the figure. A box plot is also shown within each violin plot. For better visualization of the TMB distribution over the whole age regime a few points with a relatively high mutation burden are not shown in some of the figures. Here, we took the latest data from the TCGA database with a much larger number of cases in contrast to previous studies[8] which used an old version of the database with a limited number of cases. As an example, the total number of COAD and READ patients is 224[8], while we have 486 patients for the two cancer types here. We also separate the cancer types based on their origins, instead of combining them together as in previous studies[7,8]. The different overall mutation rates (see Figures 1-4 in the main text) reflect the different underlying evolutionary processes of these cancer types.

Binning age groups does not change our results same as observed in previous studies[9]. In addition, we found similar results by using all patient data from WGS (see Figure 4 in the main text and the Pearson/Spearman correlations shown Table VII). Besides, we found that a rather accurate mutation rate could be reached by binning the WES patient data by ages as the value we obtained is almost the same (13/year for LAML as an example, see Figure 1(a) in the main text) as measured in other studies (13±2 for LAML in the reference[10]). Therefore, our results are robust and do not depend on which method we used for patient data.

### II. THE FIVE-YEAR RELATIVE SURVIVAL RATE OF CANCERS

The five-year relative survival rates[11] for various cancers are shown in Fig. S4. Survival rate is defined as the ratio of the percentage of patients who are alive for 5 years after the diagnosis of cancer to the percentage of people in the general population alive over the same time period with respect to age, sex and race[12]. Three out of the four of the most lethal cancers shown in Fig. S4 (pancreatic, liver and lung cancers, denoted by the yellow box) have relatively high TMB (see Fig. 2 in the main text). We do not have sufficient mutation data for Mesothelioma, the other lethal cancer, which requires wide and deep sequencing of the mesothelioma patients[13,14].

### III. CORRELATION OF SNVS FROM THE WHOLE GENOME AND EXOME

Among the 2834 donors of the ICGC/TCGA Whole Genomes Consortium, we used 2583 in this study which are found to have the optimal quality[7]. From the number of SNVs from coding and noncoding DNA of each patient, we plot the number of SNVs for the whole-genome (sum of the SNVs from coding and noncoding DNA) as a function of SNVs number from the whole-exome (see Fig. S5). Each circle represents the data from one patient. A strong linear correlation is found between these two variables (with the Pearson correlation coefficient $\rho = 0.98$ and a P-value smaller than 0.001). The exome fraction is about 1% of the total genome in human[15], and the total number of SNVs in exome is also about 1% of the value from the whole genome (see the magenta solid line in Fig. S5). Therefore, our definition of tumor mutation burden (TMB, the number of SNVs per Megabase) does not depend on which dataset (WES, or WGS) is used. This is further confirmed by analyzing the WES and WGS data which show that the conclusions are robust.

### IV. THERE ARE TWO TYPES OF TMB DISTRIBUTION FOR THE SIXTEEN CANCER TYPES

Instead of considering the TMB distribution over different age groups, we also investigated the overall TMB distribution for every cancer discussed in this article. From the patient mutation data obtained from TCGA[1], we calculated the TMB distributions for the sixteen cancers (see Fig. S6 and Fig. S7). Remarkably, for the nine cancers listed in Fig. 1 and Fig. 3 in the main text, we found that their TMB distribution can be well described by a universal Gaussian function (see the green lines in Fig. S6 and the corresponding functional forms are presented in the upper right box of the figures). In contrast, a power-law ($P(TMB) \sim TMB^b$) relation is observed (except for PAAD and LUSC) for those cancers with high TMB listed in Fig. 2 in the main text (see the green lines and their functional forms in Fig. S7).

For all cancers, the distribution of patient age at diagnosis (PAD) can be approximated by a Gaussian function (see the discussion below for the PAD distribution for both sexes and Fig. S19 and Fig. S20). Given a Gaussian distribution for PAD (see Fig. S8**a**) and a linear relation between TMB and PAD (see Fig. S8**b**) as noted in Fig. 1 and Fig. 3 in the main text, we would obtain a Gaussian distribution for TMB (Fig. S8**c**), thus rationalizing the findings in Fig. 5c in the main text and Fig. S6.

### V. TMB DISTRIBUTION FOR BOTH THE SEXES

We investigated the TMB distribution for both the sexes after separating the patient data into two groups according to their gender. Interestingly, similar patterns (see Fig. S9 and Fig. S10) are found (see Fig. S6 and Fig. S7). After rescaling the TMB and the frequency, the TMB distributions for the both sexes across the nine cancers (listed in Fig. 1 and Fig. 3 in the main text) again collapse onto a single curve represented by the standard normal distribution (see Fig. S11**a**). In contrast, for cancers (except PAAD and LUSC) shown in Fig. 2 in the main text, the rescaled TMB distributions collapse onto a single straight line with slope value of -1 (see Fig. S11**b**), which is the same as noted in Fig. 5d in the main text.

The rescaling approach used in Fig. 5 in the main text and Fig. S11 is often used in other areas in order to reveal whether there is a universal underlying mechanism hidden in the data set from different measurements. Our analysis clearly shows that a clock-like mutation process indeed exists in cancers shown in Fig. 5a, although the rate can be different for each cancer type. The value of $\mu_0$ for Fig. 5b is the mean TMB value for each cancer type. Because the curves for different cancers are universal one can, in principle, use an analytic formula to relate different cancer types to each other.

### VI. THE JOINT PROBABILITY DISTRIBUTION FOR TMB AND PAD

We also calculated the joint probability distribution P(X,Y) for the rescaled TMB (TMB/$T_{med}$ ≡X) and PAD (PAD/$P_{med}$ ≡Y) where $T_{med}$ and $P_{med}$ are the median value of TMB and PAD, respectively (see Fig. S12 and Fig. S13). Due to the limited data for each cancer type, we could not obtain smooth distributions for P(X,Y). Nevertheless, we could still distinguish between the cancers with relatively low and high TMB from P(X,Y). The probability distribution P(X,Y) for cancers listed in Fig. 1 and Fig. 3 in the main text is symmetric along both axes (see Fig. S12). However, the P(X,Y) for most cancers shown in Fig. 2 in the main text is irregular and biased, extending to very large values along the horizontal axis (see Fig. S13). This is consistent with the power law fat-tail distribution observed in Fig. S7.

### VII. FRACTION ($F_{ini}$) OF ACCUMULATED MUTATIONS BEFORE THE TUMOR INITIATION

In the main text, we estimated $F_{ini}$ for cancers shown in Figs. 1,3 and Fig 2. For cancers displaying a strong positive correlation between TMB and patient age (shown in Figs. 1 and 3 in the main text), we calculated the number ($N_{af}$) of accumulated mutations after tumor initiation from the mutation rate (the slope of the green curve in the figures), and the latency period of the cancer. Then, we can obtain $F_{ini}$ from $N_{af}$ and the average TMB ($N_t$) for patients at the mean age of diagnosis which leads to

$$F_{ini} = \frac{N_t - N_{af}}{N_t}. \tag{1}$$

For cancers with high TMB (shown in Fig. 2 in the main text), we first calculated the upper bound of the total number ($N_I$) of mutations a cell can accumulate before tumor initiation by taking the product of mutation rate (about 50 mutations/per year) for normal cells[16–18], and the mean age of patients at diagnosis of the cancer. The upper bound for $F_{ini}$ can be estimated from $N_I$ and the average TMB ($N_t$) of patients at the mean age of diagnosis as,

$$F_{ini} = \frac{N_I}{N_t}. \tag{2}$$

For a few cancers (Fig. 2 of the main text), the mutation rates for normal cells are not available. Hence, the value of $F_{ini}$ for these might show some derivations from our estimation. Using these two approaches, we estimated the values of $F_{ini}$ for the 16 cancers listed in Table II (for cancers shown in Figs. 1 and 3), and Table III (for cancers shown in Fig. 2). As discussed in the main text, the majority of mutations accumulate before the tumor initiation for cancers listed in Table II, while is precisely the opposite of what is found for cancers listed in Table III.

### VIII. AGE DISTRIBUTION OF CANCER PATIENTS AT DIAGNOSIS FOR BOTH THE SEXES

There is age disparity for patients at diagnosis among different types of cancer between the two sexes (see Table VI). In principle, we should not compare the cancer risks between the two populations with different ages at diagnosis directly because it is possible that one population could accumulate the same mutation burden at a younger age and at higher rate compared with the other population. The age distributions for both the sexes at diagnosis for 9 cancers (see Figs. S19) show that the median age (see the dashed line) for women is higher than for males. However, it is the opposite for 7 other cancers shown in Fig. S20. The solid lines in these two figures come from Gaussian fits. The mean value for the Gaussian distribution gives the same trend as the median age (indicated by the dashed lines) for the age disparity observed between the two sexes. The values of the median ages are used for analysis in the main text.

### IX. THE EVOLUTION OF THE NUMBER OF MUTATIONS FOR BOTH SEXES

In our study, we assumed that both the sexes accumulate mutations in a similar fashion. Because the accumulation of mutations is a linear function of time in many types of cancers (see Figs. 1 and 3 in the main text), we expect a similar trend should hold for the accumulation of somatic mutations in both female and male populations. We plot the TMB score as a function of the patient age for both the sexes in Figs. S21-S23. For cancers with low overall TMB (Fig. S21) or high TMB, but being strongly influenced by environment, (Fig. S23), a similar positive correlation between TMB and patient age is observed. Thus, both the sexes indeed accumulate somatic mutations in a similar way and we simply adjust their TMB with a factor of 66/$P_{med}$, with 66 of the median age for all cancer patients and $P_{med}$ taken from Table VI. For cancers shown in Fig. S22, the TMBs for both the sexes almost do not change with the patient age. The correlation between cancer risk and TMB observed in Figs. S24**a** and S24**b** almost does not change after the age adjustment (see Figs. S24**c** and S24**d**), while the pattern shown in Fig. 6 is mainly changed only if the TMB score is low (THCA, KIRC, and GBM) compared with those shown in Fig. S25. The TMB for HNSC is relatively high but the same positive correlation is observed for this cancer due to the strong environmental influence. Therefore, the assumption that mutations accumulate in both the sexes in a roughly similar manner, as assumed in the main text, holds.

### X. CANCER RISK AND TMB WITHOUT AGE-ADJUSTMENT

We also examined the correlation between cancer risk and TMB by considering all the mutation data obtained from TCGA database (see Table IV). As cancer risk for the two sexes varies greatly, we included the data for both of them (see Table V) without considering age disparity (see Figs. S19 and S20) between male and female patients among different cancers. The Pearson correlation coefficient $\rho = 0.65$ is obtained for the cancer risk and TMB (see the best linear fit in Fig. S24**c**) similar to that ($p = 0.6$) found in Fig. S24**a** with age-adjusted TMB. A higher value ($p = 0.7$) is obtained after removing SKCM cancer as shown in Fig. S24**d**. This value again is almost the same ($p = 0.69$) as calculated using the data in Fig. S24**b**. Although cancer risk is related to the tumor mutation burden, the TMB alone cannot explain more than 50% ($\sim 0.7^2$) of the differences in cancer risks among different tissue types.

### XI. CANCER RISK FOR BOTH THE SEXES CORRELATES WITH TMB AFTER AGE-ADJUSTMENT

The cancer risk as a function of TMB (without age-adjustment) for both sexes across different cancers are shown in Fig. S25. The data in this figure is the same as used in Fig. S24**c** but with lines connect the data for male and female populations for each cancer type. From Fig. S25**a**, it appears that there is no strong correlation between the cancer risk and TMB between males and females for cancers with low TMB. The red and blue lines show positive and negative correlations, respectively and the orange lines indicate the same TMB. On the other hand, the cancer risk is frequently associated with TMB as the latter reaches high values (see Fig. S25**b**). However, we neglected the known age disparity between female and male patients at diagnosis to obtain the results in Figs. S19 and S20. After we adjust the TMB by patient ages, as discussed in the main text, we find that the mutation burden is the critical factor in determining the different risks between female and male patients across many types of cancer.

### XII. TMB, PATIENT AGE AND IMMUNOTHERAPY FOR METASTATIC MELANOMA PATIENTS

We investigated the influence of TMB, and patient age on response to immunotherapy. As an example, we first examined the age distribution for all the patients and the ones who show favorable response to immune checkpoint blockade from a clinical study for metastatic melanoma[19] (see Fig. S26**a**). The age varies considerably in both the cases. The median age for patients who show response is 65 years while it is 61.5 years for the whole population. Meanwhile, the number of non-synonymous mutations for patients for the same cancer also varies greatly, as illustrated in Fig. S26**b**. The median value (309) of mutations for patients showing response is much higher than the value (197) of the whole population and a significant difference exists in the TMB between patients with and without clinical benefit to immunotherapy[19]. We observed that the TMB is correlated with the patient age for melanoma patients (see Fig. 3c in the main text). Thus, the patient age could also influence the responses to immunotherapy. However, we find that the patients whose age is $\geq 65$ show a similar response (27%) as younger patients (22%).

In order to remove the influence of TMB and focus solely on age, we compared the patient age with different responses with similar TMB. For a similar mutation burden level, (<5% difference), we find a much higher fraction of old patients compared to young patients showing favorable response (see Fig. S26**c**, 63% versus 37%). A similar conclusion was reached by analyzing the data from other immunotherapy studies[20,21] with 75% and 100% fraction of older patients showing response but the sample sizes are smaller (12 and 6 cases, respectively).

We explain the data in Fig. S26**c** using the dynamics of accumulation of mutations discussed in the main text. A constant mutation rate is observed for melanoma patients (see Fig. 3c in the main text). A similar time period ($\tau_d$) for all the patients would be expected from the initiation to the detection of melanoma[8]. Consider two patients A and B with ages $T_A$ and $T_B$ with the same TMB, denoted by $N_T$. The fraction ($F_i^{A/B}$) of mutations accumulated before the tumor initiation for both the patients is,

$$F_i^{A/B} = \frac{N_T - \alpha_{A/B}\tau_d}{N_T}, \tag{3}$$

where $\alpha_{A/B}$ is the rate for the mutation accumulation in patient A or B. If $T_A > T_B$, then the mutation rate obeys $\alpha_A < \alpha_B$ because of $N_T$ for both the patients is the same. Therefore, the older patient would accumulate more mutations ($F_i^A > F_i^B$) in the trunk of the tumor phylogenetic tree. A better response to immunotherapy would be expected if all tumor cells contain the neoantigens recognized by T-cells compared with the case that these neoantigens only appear in a tiny fraction of cell populations. This model explains the higher fraction of older patients showing response to immunotherapy with the same TMB, noted in the clinical studies cited above. This is consistent with the result of Figure 8(a) in the main text where a better survival probability is observed for the patient group with a higher clonal TMB.

### XIII. THE PHYLOGENETIC TREES FOR LUNG ADENOCARCINOMA PATIENTS FROM MULTI-REGION WHOLE EXOME SEQUENCING

The SNVs obtained from the multi-region sequencing of each tumor are first clustered using the PyClone Dirichlet process clustering[22]. The mutation clusters are filtered based on the pigeonhole principle and crossing rule to ensure the constructed phylogenetic tree is accurate[23]. Based on the mutation clusters and also the values of the mean cancer cell prevalence, the phylogenetic trees (see Fig. S27 and S28) are constructed using the tool CITUP[24] for the lung adenocarcinoma patients discussed in Fig. 8(e) in the main text. A detailed description of the phylogenetic tree construction can be found in the SI of the reference[23].

### XIV. CANCER RISKS FOR BOTH SEXES

#### Bladder Urothelial Carcinoma

The lifetime risk of urinary bladder cancer is 1.12% and 3.76% for women and men, respectively (https://seer.cancer.gov/csr/1975_2014/)[25] and 95% of the cancers are bladder urothelial Carcinomas[26]. Therefore, the lifetime risk of bladder urothelial carcinoma for female and male is 1.12%×0.95=1.06% and 3.76% × 0.95=3.57%.

**Colon Adenocarcinoma**

The lifetime risk of colorectal cancer is 4.15% and 4.49% for women and men, respectively[25]. About 71% of colorectal cancers arise in the colon[27]. Thus, the lifetime risk of colon adenocarcinoma for female and male is $4.15\% \times 0.71 = 2.95\%$ and $4.49\% \times 0.71 = 3.19\%$.

**Glioblastoma multiforme**

The lifetime risk of brain cancer is 0.54% and 0.7% for women and men, respectively[25]. About 81% of brain cancers are gliomas and glioblastomas represent 45% of gliomas[28]. Thus, the lifetime risk of glioblastoma multiforme for female and male is $0.54\% \times 0.81 \times 0.45 = 0.20\%$ and $0.7\% \times 0.81 \times 0.45 = 0.255\%$.

**Head and Neck Squamous Cell Carcinoma**

The lifetime risk of oral or pharynx cancer is 0.68% for women and is 0.12% for laryngeal cancer[25]. For men, the lifetime risk of oral or pharynx cancer is 1.61% and it is 0.55% for Laryngeal cancer. Therefore, the lifetime risk of head and neck squamous cell carcinoma (including oral, pharynx and laryngeal cancers) for women and men is 0.80% and 2.16%, respectively.

**Kidney Chromophobe**

The lifetime risk of kidney and renal pelvis is 1.2% and 2.09% for women and men, respectively[25]. Among these cancers, 80% of them are from clear cell, 15% are from papillary cell and 5% are from chromophobe[29]. From these, we estimate the lifetime risk for kidney chromophobe carcinoma is $1.2\% \times 0.05 = 0.06\%$ and $2.09\% \times 0.05 = 0.105\%$ for women and men, respectively.

**Kidney Renal Clear Cell**

Similarly, the lifetime risk of kidney renal clear cell carcinoma is $1.2\% \times 0.8 = 0.96\%$ and $2.09\% \times 0.8 = 1.67\%$ for women and men, respectively.

**Kidney Renal Papillary Cell Carcinoma**

Again, the lifetime risk of kidney renal papillary cell carcinoma is $1.2\% \times 0.15 = 0.18\%$ and $2.09\% \times 0.15 = 0.314\%$ for women and men, respectively.

**Acute Myeloid Leukemia**

The lifetime risk of acute myeloid leukemia is 0.43% and 0.55% for women and men, respectively[25].

**Liver Hepatocellular Carcinoma**

The lifetime risk of liver and intrahepatic bile duct cancer is 0.6% and 1.39% for women and men, respectively[25]. Hepatocellular carcinoma represents 90% of these cancers[30]. Hepatitis C infection is among 10% of all hepatocellular carcinoma[31] and 1% of the US population is infected with Hepatitis[32]. According to the calculation in previous research[33], the lifetime risk of liver hepatocellular carcinoma is 0.50% and 1.15% for women and men not infected by Hepatitis C, respectively.

**Lung Adenocarcinoma**

The lifetime risk of lung and bronchus cancer is 5.95% and 6.85% for women and men, respectively[25]. About 40% of these cancers are adenocarcinoma and 30% of them are squamous cell carcinoma[34]. Therefore, the lifetime risk is $5.95\% \times 0.4 = 2.38\%$ and $6.85\% \times 0.4 = 2.74\%$ for females and males, respectively.

**Lung Squamous Cell Carcinoma**

Similarly to the analysis for lung adenocarcinoma, the lifetime risk is $5.95\% \times 0.3 = 1.79\%$ and $6.85\% \times 0.3 = 2.06\%$ for females and males, accrodingly.

**Pancreatic Adenocarcinoma**

The lifetime risk of pancreatic cancer is 1.54% and 1.58% for females and males, respectively[25]. About 96% of these cancers are exocrine cancers with adenocarcinoma being the dominant one of 95%[35]. The lifetime risk is $1.54\% \times 0.96 \times 0.95 = 1.40\%$ and $1.58\% \times 0.96 \times 0.95 = 1.44\%$ for females and males, respectively.

**Rectum Adenocarcinoma**

The lifetime risk of colorectal cancer is 4.15% and 4.49% for females and males as mentioned above[25]. About 29% of these cancers arise in the rectum[27]. Therefore, the lifetime risk of rectum adenocarcinoma is $4.15\% \times 0.29 = 1.20\%$ and $4.49\% \times 0.29 = 1.30\%$ for female and male, respectively.

**Cutaneous Skin Melanoma**

The lifetime risk of cutaneous skin melanoma for females and males is 1.72% and 2.77%, respectively[25].

**Stomach Adenocarcinoma**

The lifetime risk of stomach cancer is 0.65% and 1.05% for females and males, respectively[25]. Adenocarcinoma represents about 95% of these cancers[35]. Thus, the lifetime risk is $0.65\% \times 0.95 = 0.618\%$ and $1.05\% \times 0.95 = 0.998\%$ for both sexes, accrodingly.

**Thyroid Carcinoma**

The lifetime risk of thyroid carcinoma for females and males is 1.79% and 0.63%, respectively[25].

### XV. TABLES

Table I shows the number of patients for each type of cancer used in our analyses. Table II and III give the fraction ($F_{ini}$) of accumulated mutations before the initiation of tumors for two categories of cancer shown in Figs. 1,3 and Fig. 2 in the main text, respectively. Table VI lists the median age for male and female patients at diagnosis. Table IV gives the median value of (both synonymous and non-synonymous) mutations per megabase for both sexes across 16 types of cancer. Table V shows the cancer risk for both female and male populations in 16 different types of cancer.

### REFERENCES

[1]Weinstein, J. N., Collisson, E. A., Mills, G. B., Shaw, K. R. M., Ozenberger, B. A., Ellrott, K., ... & Cancer Genome Atlas Research Network. The cancer genome atlas pan-cancer analysis project. *Nature genetics*, 45(10), 1113, 2013

[2]Hutter, C., & Zenklusen, J. C. The cancer genome atlas: creating lasting value beyond its data. Cell, 173(2), 283-285, 2018.

[3]Cancer Genome Atlas Research Network. Integrated genomic analyses of ovarian carcinoma. *Nature*, 474(7353), 609, 2011

[4]Berger MF, Lawrence MS, Demichelis F, Drier Y, Cibulskis K, Sivachenko AY, Sboner A, Esgueva R, Pflueger D, Sougnez C, Onofrio R. The genomic complexity of primary human prostate cancer. *Nature*, 470(7333), 214, 2011.

[5]Cibulskis K, Lawrence MS, Carter SL, Sivachenko A, Jaffe D, Sougnez C, Gabriel S, Meyerson M, Lander ES, Getz G. Sensitive detection of somatic point mutations in impure and heterogeneous cancer samples. *Nature biotechnology*, 31(3):213, 2013.

[6]Samur, Mehmet Kemal. RTCGAToolbox: a new tool for exporting TCGA Firehose data. *PloS one*, 9, e106397, 2014.

[7]The ICGC/TCGA Pan-Cancer Analysis of Whole Genomes Consortium. Pan-cancer analysis of whole genomes. *Nature*, 578(7793), 82-93, 2020.

[8]Tomasetti C, Vogelstein B, Parmigiani G. Half or more of the somatic mutations in cancers of self-renewing tissues originate prior to tumor initiation. *Proceedings of the National Academy of Sciences*, 110(6):1999-2004, 2013.

[9]Alexandrov LB, Jones PH, Wedge DC, Sale JE, Campbell PJ, Nik-Zainal S, & Stratton MR. Clock-like mutational processes in human somatic cells. *Nature genetics*, 47(12):1402, 2015.

[10]Welch JS, Ley TJ, Link DC, Miller CA, Larson DE, Koboldt DC, Wartman LD, Lamprecht TL, Liu F, Xia J, & Kandoth C. The origin and evolution of mutations in acute myeloid leukemia. *Cell*, 150(2):264-78, 2012.

[11]Noone AM, Howlader N, Krapcho M, Miller D, Brest A, Yu M, Ruhl J, Tatalovich Z, Mariotto A, Lewis DR, Chen HS, Feuer EJ, Cronin KA (eds). *SEER Cancer Statistics Review*, 1975-2015, National Cancer Institute. Bethesda, MD, 2018.

[12]Baranovsky, A., & Myers, M. H. Cancer incidence and survival in patients 65 years of age and older. *CA: a cancer journal for clinicians*, 36(1), 26-41, 1986.

[13]Guo, G., Chmielecki, J., Goparaju, C., Heguy, A., Dolgalev, I., Carbone, M., ... & Pass, H. I. Whole-exome sequencing reveals frequent genetic alterations in BAP1, NF2, CDKN2A, and CUL1 in malignant pleural mesothelioma. *Cancer research*, 75(2), 264-269, 2015

[14]Bueno, R., Stawiski, E. W., Goldstein, L. D., Durinck, S., De Rienzo, A., Modrusan, Z., ... & Sciaranghella, D. Comprehensive genomic analysis of malignant pleural mesothelioma identifies recurrent mutations, gene fusions and splicing alterations. *Nature genetics*, 47(3), 407, 2015.

[15]Venter JC, Adams MD, Myers EW, Li PW, Mural RJ, ... & Gocayne JD. The sequence of the human genome. *Science*, 291(5507):1304-51, 2001.

[16]Blokzijl, F., De Ligt, J., Jager, M., Sasselli, V., Roerink, S., Sasaki, N., ... & Nijman, I. J. Tissue-specific mutation accumulation in human adult stem cells during life. *Nature*, 538(7624), 260, 2016.

[17]Roerink, S. F., Sasaki, N., Lee-Six, H., Young, M. D., Alexandrov, L. B., Behjati, S., ... & Pronk, A. Intra-tumour diversification in colorectal cancer at the single-cell level. *Nature*, 556(7702), 457, 2018.

[18]Yokoyama, A., Kakiuchi, N., Yoshizato, T., Nannya, Y., Suzuki, H., Takeuchi, Y., ... & Fujii, Y. Age-related remodelling of oesophageal epithelia by mutated cancer drivers. *Nature*, 565(7739), 312, 2019.

[19]Eliezer M Van Allen, Diana Miao, Bastian Schilling, Sachet A Shukla, Christian Blank, ... & Levi A. Garraway, Genomic correlates of response to CTLA4 blockade in metastatic melanoma. *Science*, 350:207–211, 2015.

[20]Hellmann MD, Nathanson T, Rizvi H, Creelan BC, Sanchez-Vega F,... & Wolchok JD. Genomic features of response to combination immunotherapy in patients with advanced non-small-cell lung cancer. *Cancer cell*, 33(5):843-52, 2018.

[21]Rizvi NA, Hellmann MD, Snyder A, Kvistborg P, Makarov V,... &Chan TA. Mutational landscape determines sensitivity to PD-1 blockade in non?small cell lung cancer. *Science*, 348(6230):124-8, 2015.

[22]Roth A, Khattra J, Yap D, Wan A, Laks E, Biele J, Ha G, Aparicio S, Bouchard-Cote A, & Shah SP. PyClone: statistical inference of clonal population structure in cancer. *Nature methods*, 11(4):396-8, 2014.

[23]Jamal-Hanjani M, Wilson GA, McGranahan N, Birkbak NJ, Watkins TB,... & Swanton C. Tracking the evolution of non-small-cell lung cancer. *New England Journal of Medicine*, 1;376(22):2109-21, 2017.

[24]Malikic S, McPherson AW, Donmez N, & Sahinalp CS. Clonality inference in multiple tumor samples using phylogeny. *Bioinformatics*, 1;31(9):1349-56, 2015.

[25]Howlader N, Noone AM, Krapcho M, Miller D, Bishop K, Kosary CL, Yu M, Ruhl J, Tatalovich Z, Mariotto A, Lewis DR, Chen HS, Feuer EJ, Cronin KA (eds). *SEER Cancer Statistics Review*, 1975-2014, National Cancer Institute. Bethesda, MD, 2017.

[26]Kim, M., Jeong, C. W., Kwak, C., Kim, H. H., and Ku, J. H. Are urothelial carcinomas of the upper urinary tract a distinct entity from urothelial carcinomas of the urinary bladder? Behavior of urothelial carcinoma after radical surgery with respect to anatomical location: a case control study. *BMC cancer*, 15(1), 149, 2015

[27]American Cancer Society, Colorectal Cancer Facts & Figures, *American Cancer Society*, Atlanta, GA (2011)

[28]Ostrom, Q. T., Bauchet, L., Davis, F. G., Deltour, I., Fisher, J. L., Langer, C. E., ... and Wrensch, M. R. The epidemiology of glioma in adults: a "state of the science" review. *Neuro-oncology*, 16(7), 896-913, 2014.

[29]Ebele J. N., Sauter G., Epstein J. I., Sesterhenn I. A. Genetics of Tumors of the Urinary system and Male Genital Organs. In: *World Health Organization Classification of Tumors.* Lyon: IARC; p. 2-3, 2004.

[30]Altekruse, S. F., McGlynn, K. A., & Reichman, M. E. Hepatocellular carcinoma incidence, mortality, and survival trends in the United States from 1975 to 2005. *Journal of clinical oncology*, 27(9), 1485, 2009.

[31]Davila, J. A., Morgan, R. O., Shaib, Y., McGlynn, K. A., & El-Serag, H. B. Hepatitis C infection and the increasing incidence of hepatocellular carcinoma: a population-based study. *Gastroenterology*, 127(5), 1372-1380, 2004.

[32]Armstrong, G. L., Wasley, A., Simard, E. P., McQuillan, G. M., Kuhnert, W. L., & Alter, M. J. The prevalence of hepatitis C virus infection in the United States, 1999 through 2002. *Annals of internal medicine*, 144(10), 705-714, 2006.

[33]Tomasetti, C., & Vogelstein, B. Variation in cancer risk among tissues can be explained by the number of stem cell divisions. *Science*, 347(6217), 78-81, 2015.

[34]Hong, W. K. *American Association for Cancer Research*, Holland Frei Cancer Medicine 8 (People's Medical Pub. House, Shelton, Conn., ed. 8th, 2010).

[35]American Cancer Society, www.cancer.org.

[36]Larson, R. A., LeBeau, M. M., Vardiman, J. W., & Rowley, J. D. Myeloid leukemia after hematotoxins. *Environmental health perspectives*, 104(suppl 6), 1303-1307, 1996.

[37]Nadler, D. L., & Zurbenko, I. G. Estimating cancer latency times using a Weibull model. *Advances in Epidemiology*, 2014.

[38]Tsyb, A. F., Parshkov, E. M., Shakhtarin, V. V., Stepanenko, V. F., Skvortsov, V. F., & Chebotareva, I. V. Thyroid cancer in children and adolescents of Bryansk and Kaluga regions. *Age*, 4(10-14):15-19, 1996.

[39]Lee, J. H., Kim, I., Seok, H., Park, I., Hwang, J., Park, J. O., ... & Roh, J. Case report of renal cell carcinoma in automobile manufacturing factory worker due to trichloroethylene exposure in Korea. *Annals of occupational and environmental medicine*, 27(1), 19, 2015.

[40]Mitchell, T. J., Turajlic, S., Rowan, A., Nicol, D., Farmery, J. H., O?Brien, T., ... & Butler, A. P. Timing the landmark events in the evolution of clear cell renal cell cancer: TRACERx renal. *Cell*, 173(3), 611-623, 2018.

[41]Salvati, M., Frati, A., Russo, N., Caroli, E., Polli, F. M., Minniti, G., & Delfini, R. Radiation-induced gliomas: report of 10 cases and review of the literature. *Surgical neurology*, 60(1), 60-67, 2003.

[42]Makimoto, Y., Yamamoto, S., Takano, H., Motoori, K., Ueda, T., Kazama, T., ... & Hanazawa, T. Imaging findings of radiation-induced sarcoma of the head and neck. *The*

*British journal of radiology*, 80(958), 790-797, 2007.

[43]Pang, Z. C., Zhang, Z., Wang, Y., & Zhang, H. Mortality from a Chinese asbestos plant: overall cancer mortality. *American journal of industrial medicine*, 32(5), 442-444, 1997.

[44]Haluza, D., Simic, S., & Moshammer, H. Temporal and spatial melanoma trends in Austria: An ecological study. *International journal of environmental research and public health*, 11(1), 734-748, 2014.

[45]Garland, C. F., Garland, F. C., & Gorham, E. D. Rising trends in melanoma an hypothesis concerning sunscreen effectiveness. *Annals of epidemiology*, 3(1), 103-110, 1993.

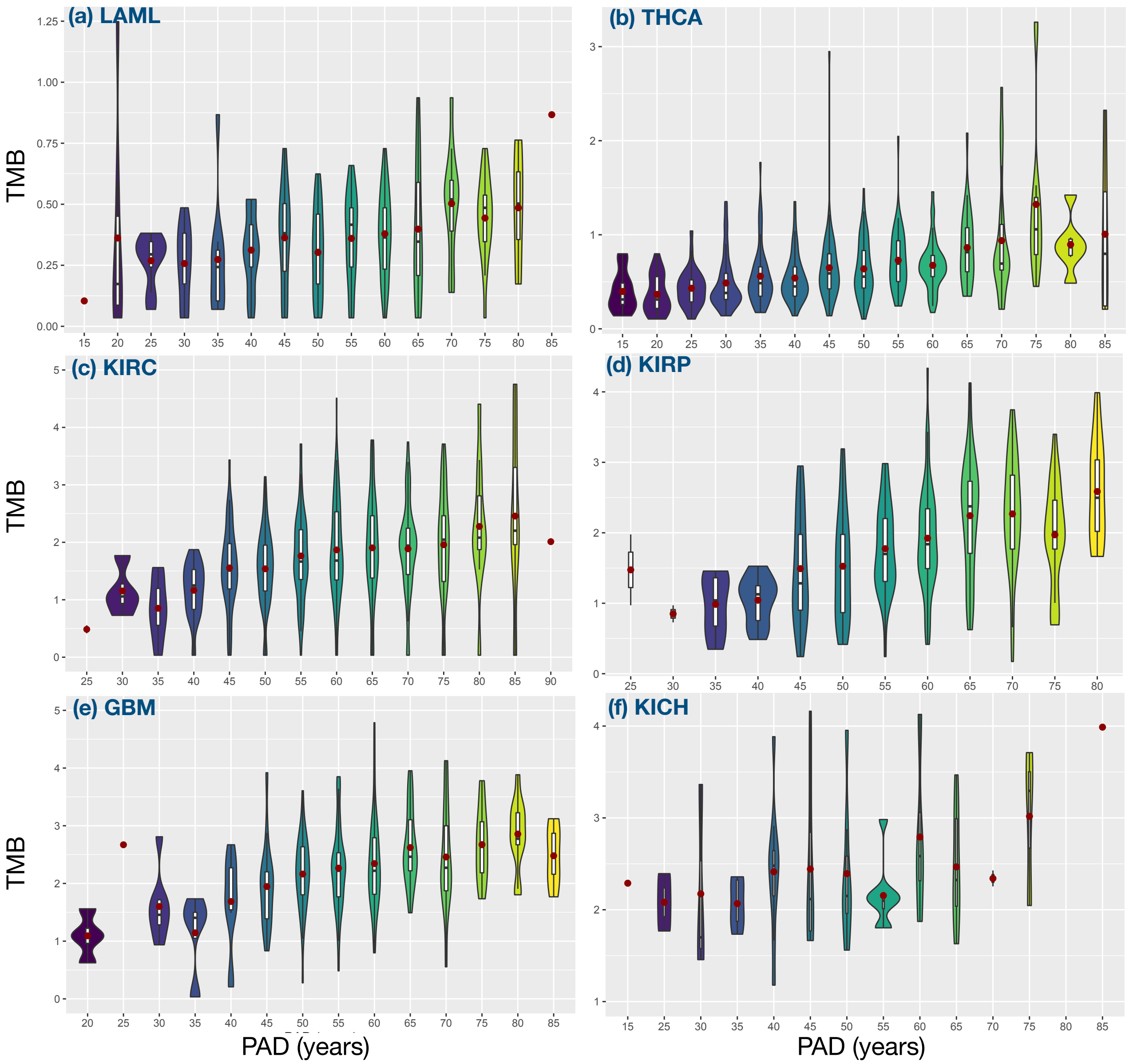

FIG. S1. Violin plots for the distribution of tumor mutation burden (TMB) as a function of patient age for low TMB cancers shown in Fig. 1 in the main text. The red dot is the mean value of the TMB. The inset with each violin plot shows a box plot. The unit for TMB is the number of mutations per Mb. The type of cancer is labeled in dark blue color. In Figs. (S2-S3), we show the same plot but for different cancer types. A few patient data with very high TMB are removed in Figs. (S2-S3) for better illustration.

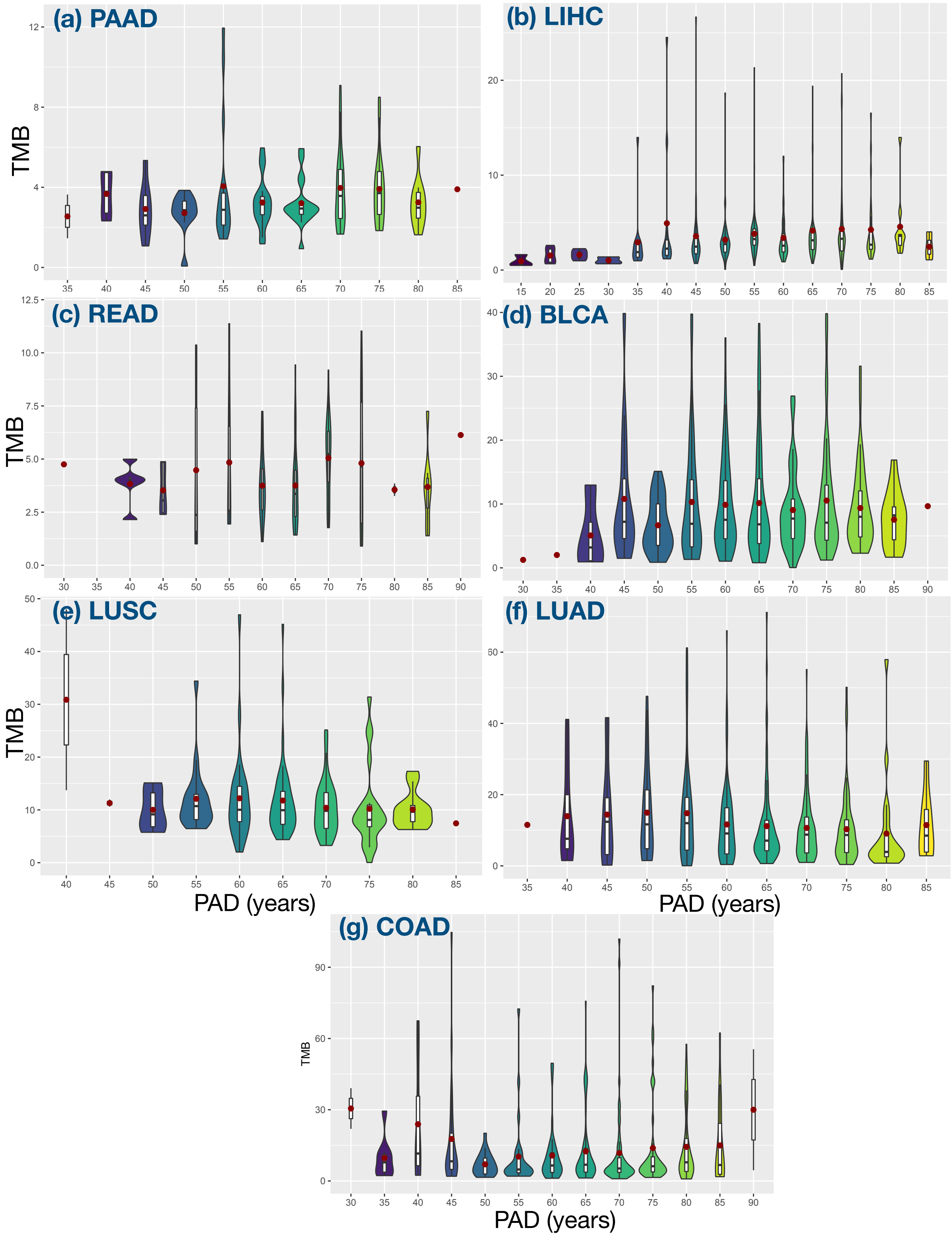

FIG. S2. Violin plots for high TMB cancers shown in Fig. 2 in the main text.

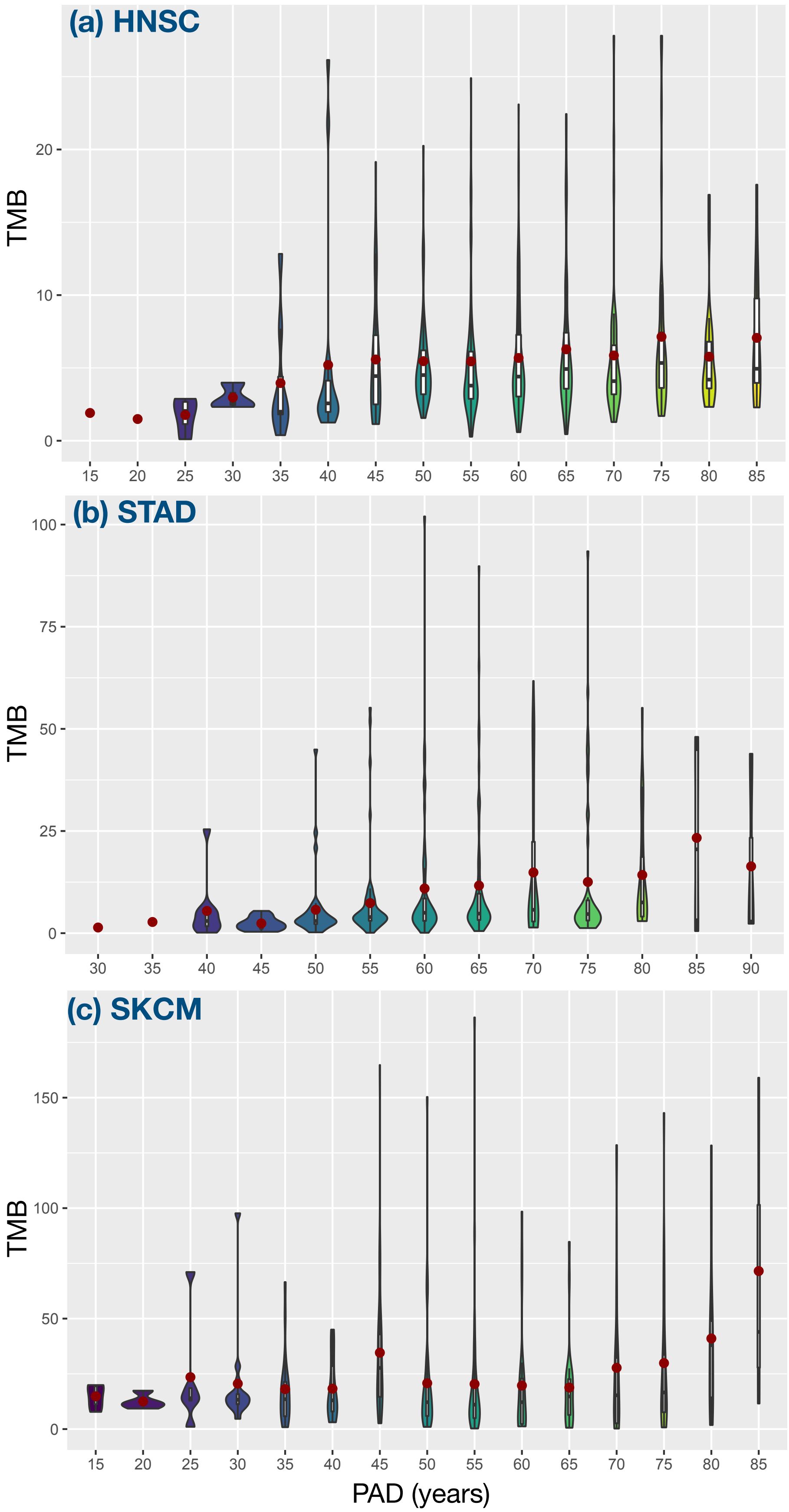

FIG. S3. Violin plots for HNSC, STAD, and SKCM shown in Fig. 3 in the main text.

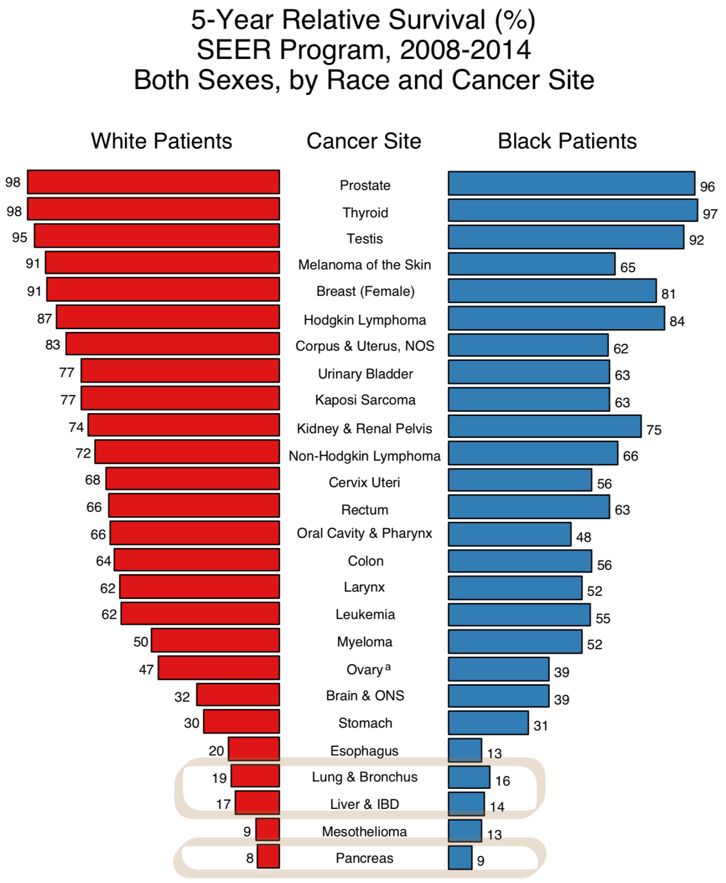

FIG. S4. Five-year relative survival rate for different cancers[11]. Three of the most lethal cancers (lung, liver and pancreatic) in the box belong to the category shown in Fig. 2 in the main text.

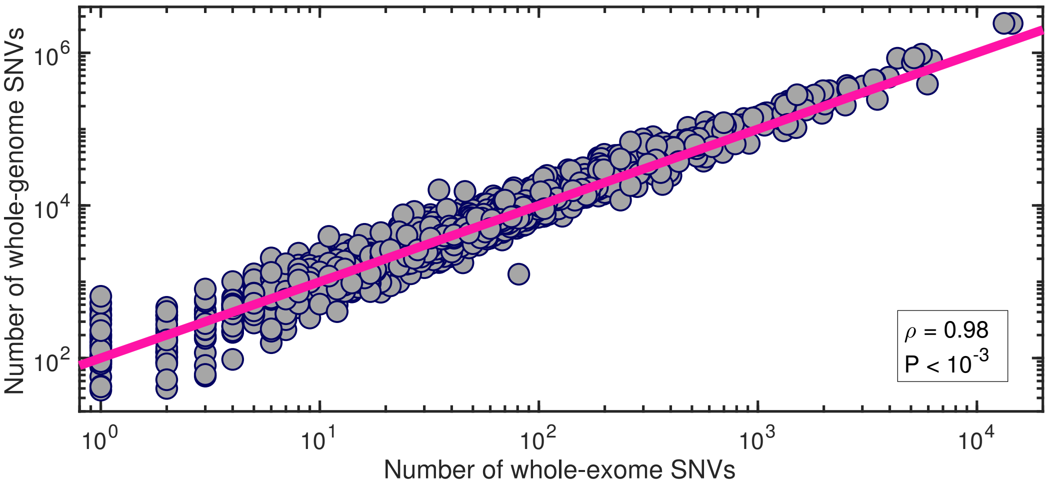

FIG. S5. The correlation between the number of SNVs for the whole-genome and that for the whole-exome. The data (filled circles) are taken from the whole-genome sequencing of 2583 cancer patients across 38 tumor types[7]. A clear linear relation (with the Pearson correlation coefficient $\rho = 0.98$ and a P-value smaller than 0.001) is observed between these two variables. The line is described by the function: $\mathrm{Y} = 100\mathrm{X}$.

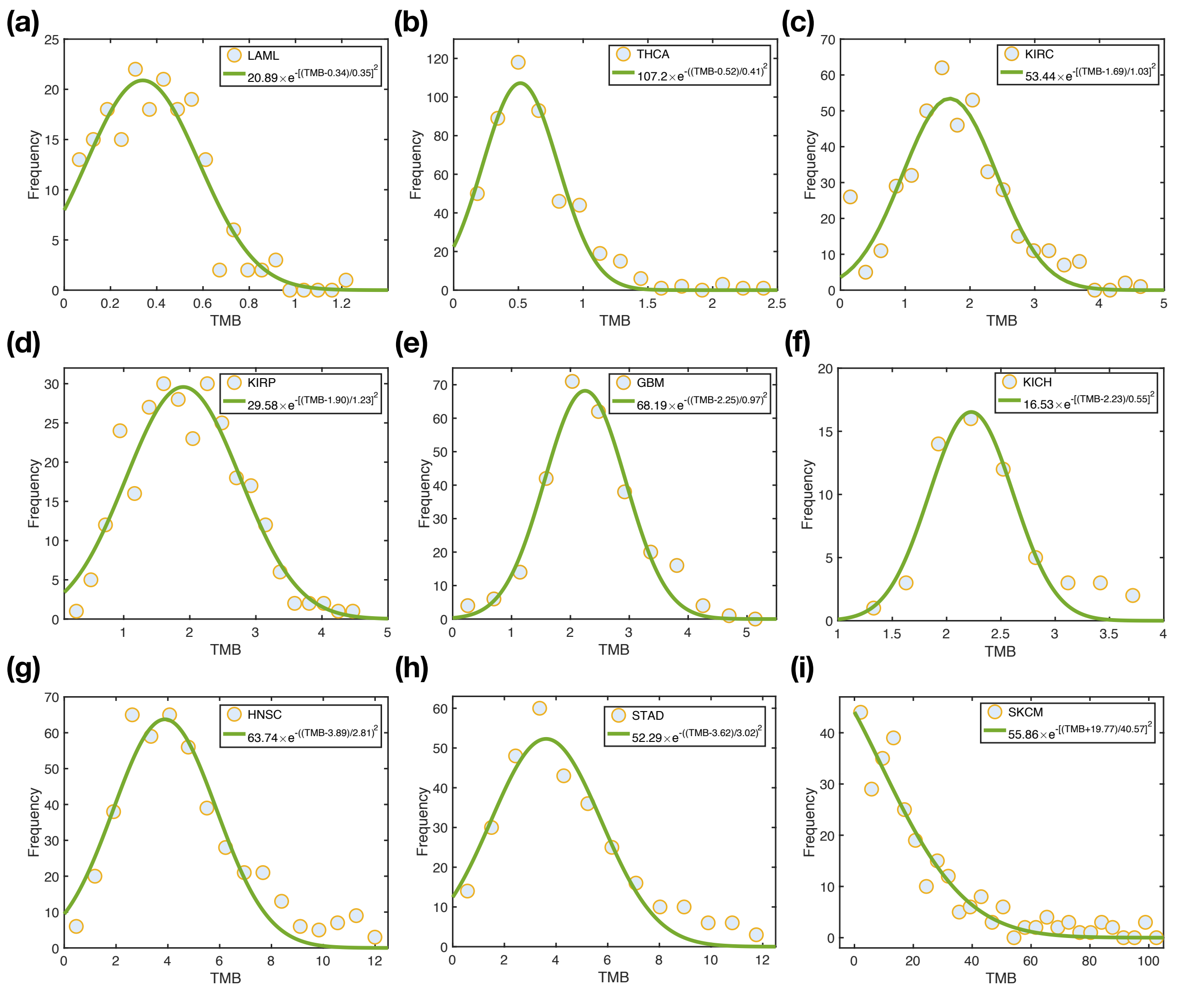

FIG. S6. The TMB distribution over the nine types of cancer showed in Fig. 1 and Fig. 3 in the main text. The data (filled circles) are taken from the TCGA database as in the main text. The solid green curves (described by the functions in the upper right box of the figure) show the best fit to the data.

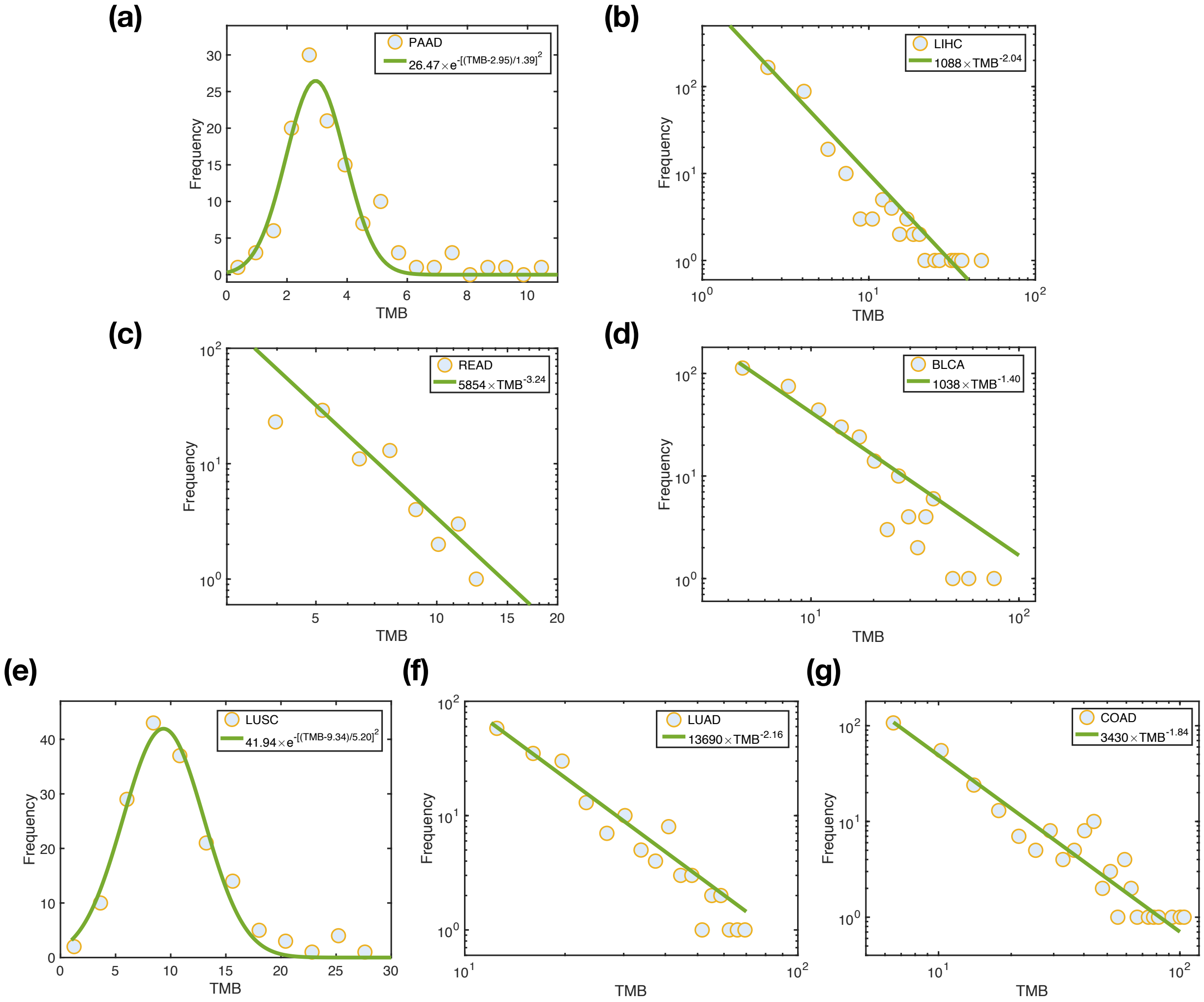

FIG. S7. The TMB distribution for the cancers analyzed in Fig. 2. The meaning of the symbols and curves are the same as in Fig. S6.

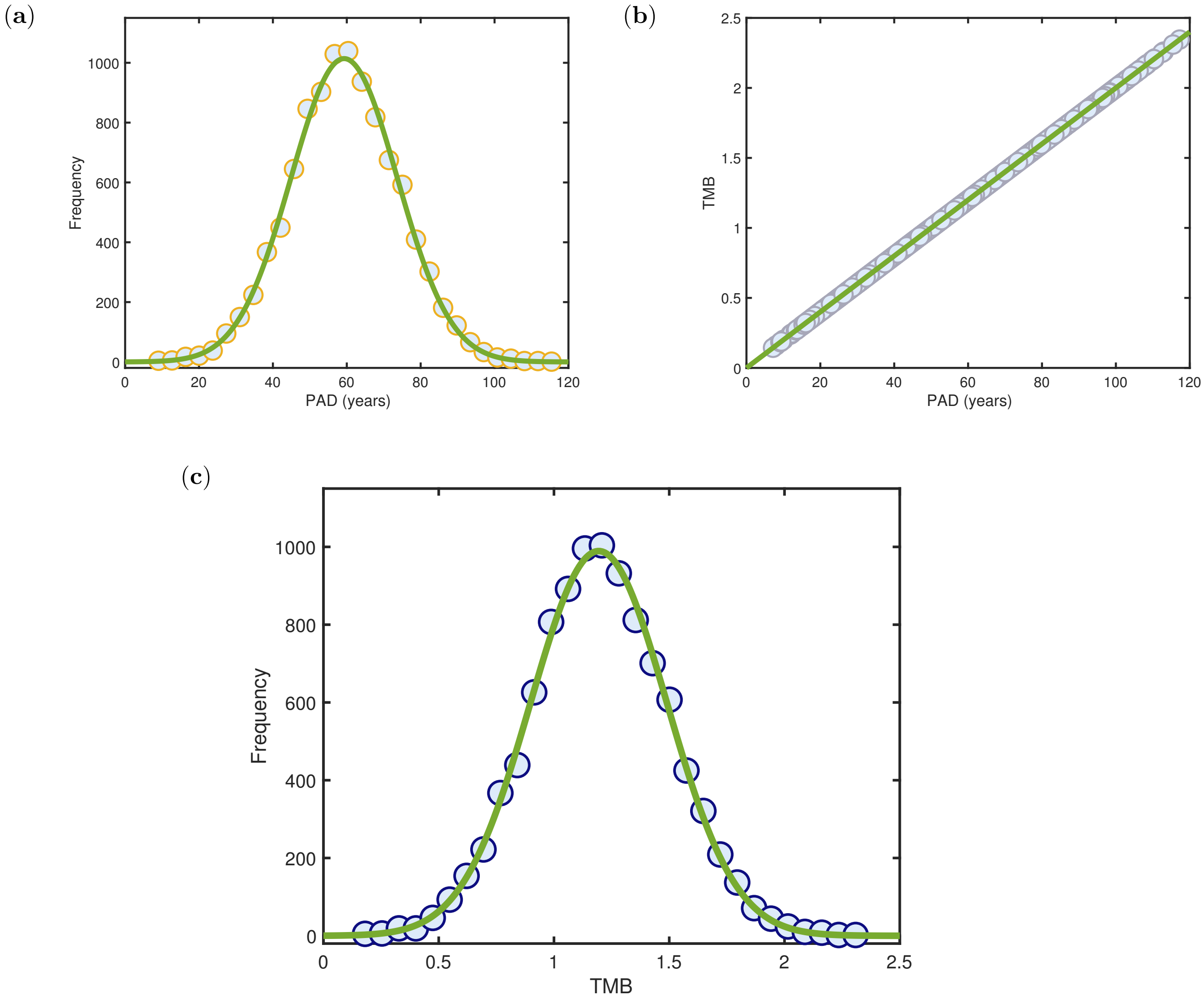

FIG. S8. (a) The distribution of PAD obtained from a 10,000 sample using the Gaussian distribution. The green curve is described by a Gaussian function, $1014 \times e^{-[(\mathrm{PAD}-59.28)/20.36]^2}$. (b) The TMB for each patient is proportional to their age (TMB $=$ 0.02$\times$ PAD, see the green line). (c) The TMB distribution obtained from the 10,000 patient samples above. The green curve is again described by a Gaussian function, $989.3 \times e^{-[(\mathrm{TMB}-1.2)/0.42]^2}$. The circles are simulation data and the green lines show the best fit to the data.

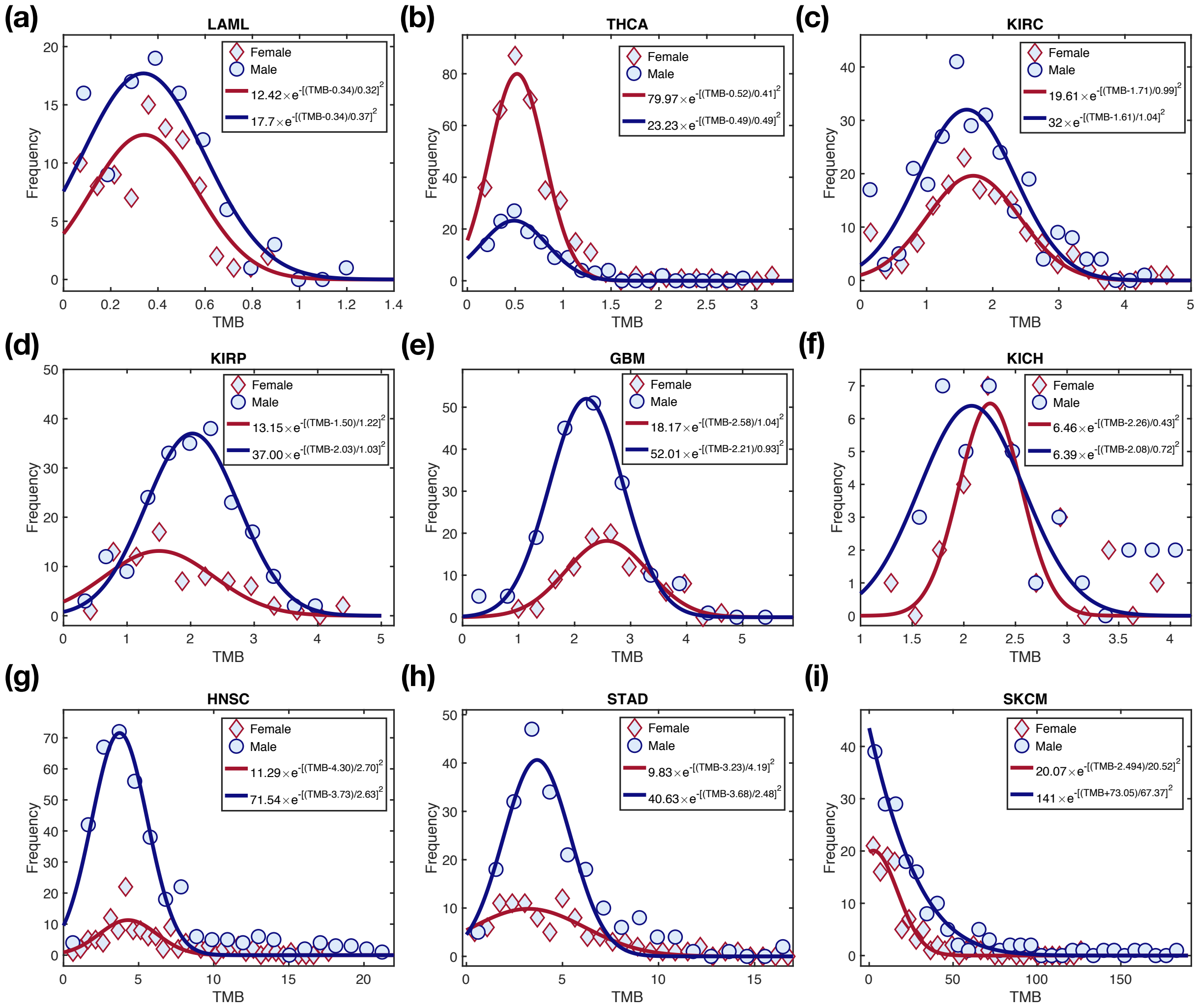

FIG. S9. The TMB distribution for both the sexes for the nine cancers analyzed in Fig. 1 and Fig. 3. The data for females and males are represented by diamonds and circles, respectively. And the corresponding fitting functions (listed in the upper right box of the figure) for the data are shown by the red and blue curves.

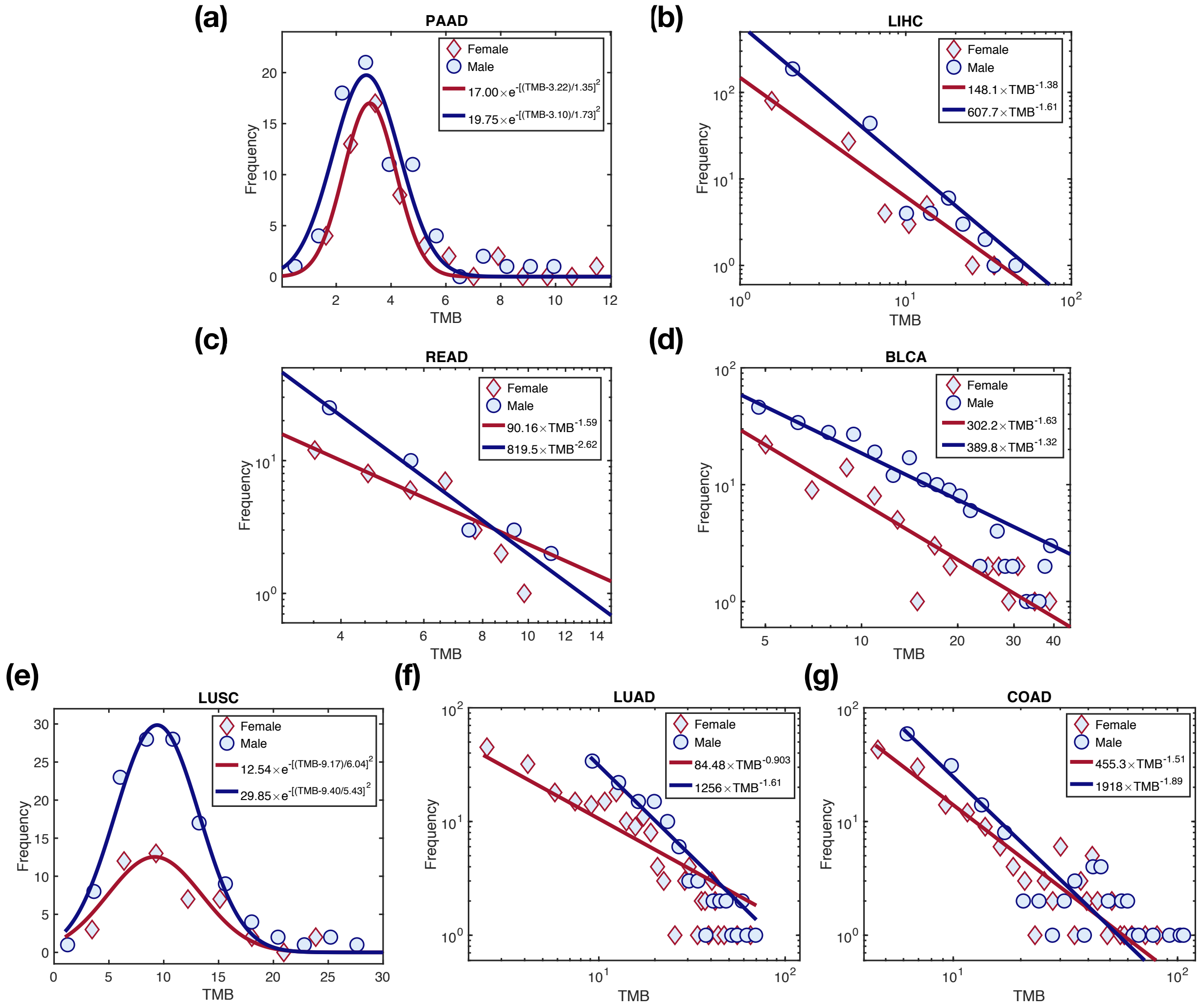

FIG. S10. The TMB distribution for both the sexes over the seven cancers analyzed in Fig. 2. The meaning of the symbols and curves are the same as in Fig. S9.

(**a**)

(**b**)

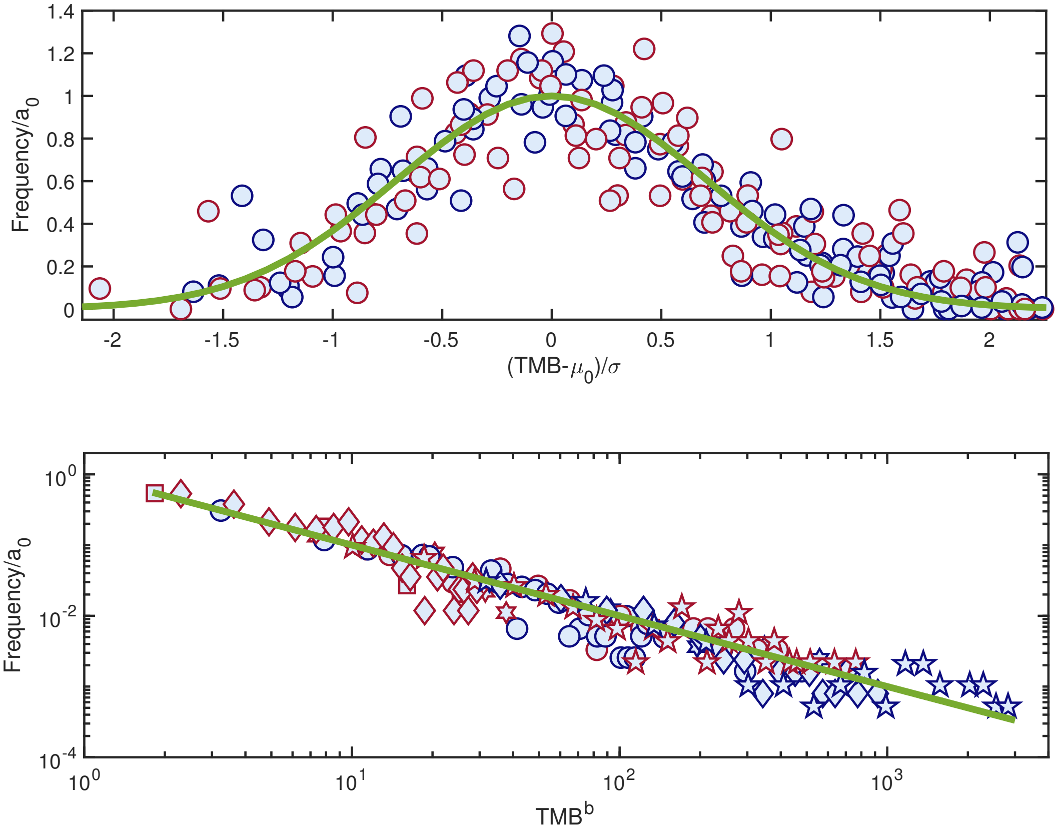

FIG. S11. The rescaled TMB distribution for the 16 cancers. (a) A Gaussian distribution (with the mean value $\mu_0$, variance $\sigma$, and coefficient $a_0$) describes (see Fig. S9) both the sexes for the low TMB cancers. The rescaled TMB distributions for these cancers collapse into the standard normal distribution (see the green line). (b) A long tail distribution, described by a power-law (see Fig. S10 showing $P(\text{TMB}) = a_0 \times \text{TMB}^b$) is found for the cancers (except for PAAD and LUSC) listed in Fig. 2. The green straight line has a slope value of -1. The red (blue) symbols represent data for females (males).

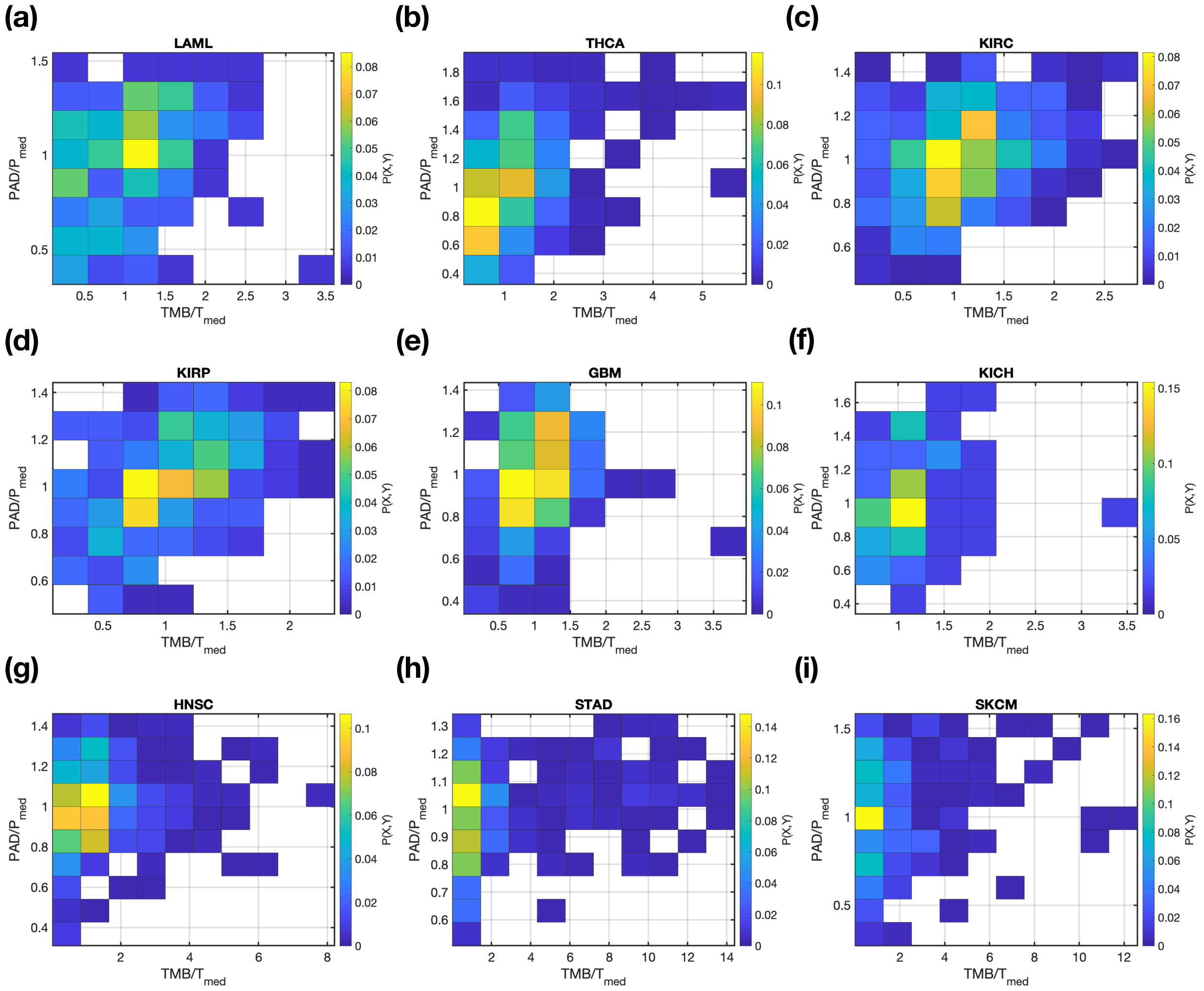

FIG. S12. The joint probability distribution, $P(X, Y)$, for TMB and PAD among the nine cancers. $X$ = TMB/T$_{med}$ with T$_{med}$ the median value of TMB. $Y$ = PAD/P$_{med}$ with P$_{med}$ the median value of PAD. The probability $P(X, Y)$ is normalized by the total number of patients in each cancer and is color coded according to the value indicated by the scale on the right side of each figure.

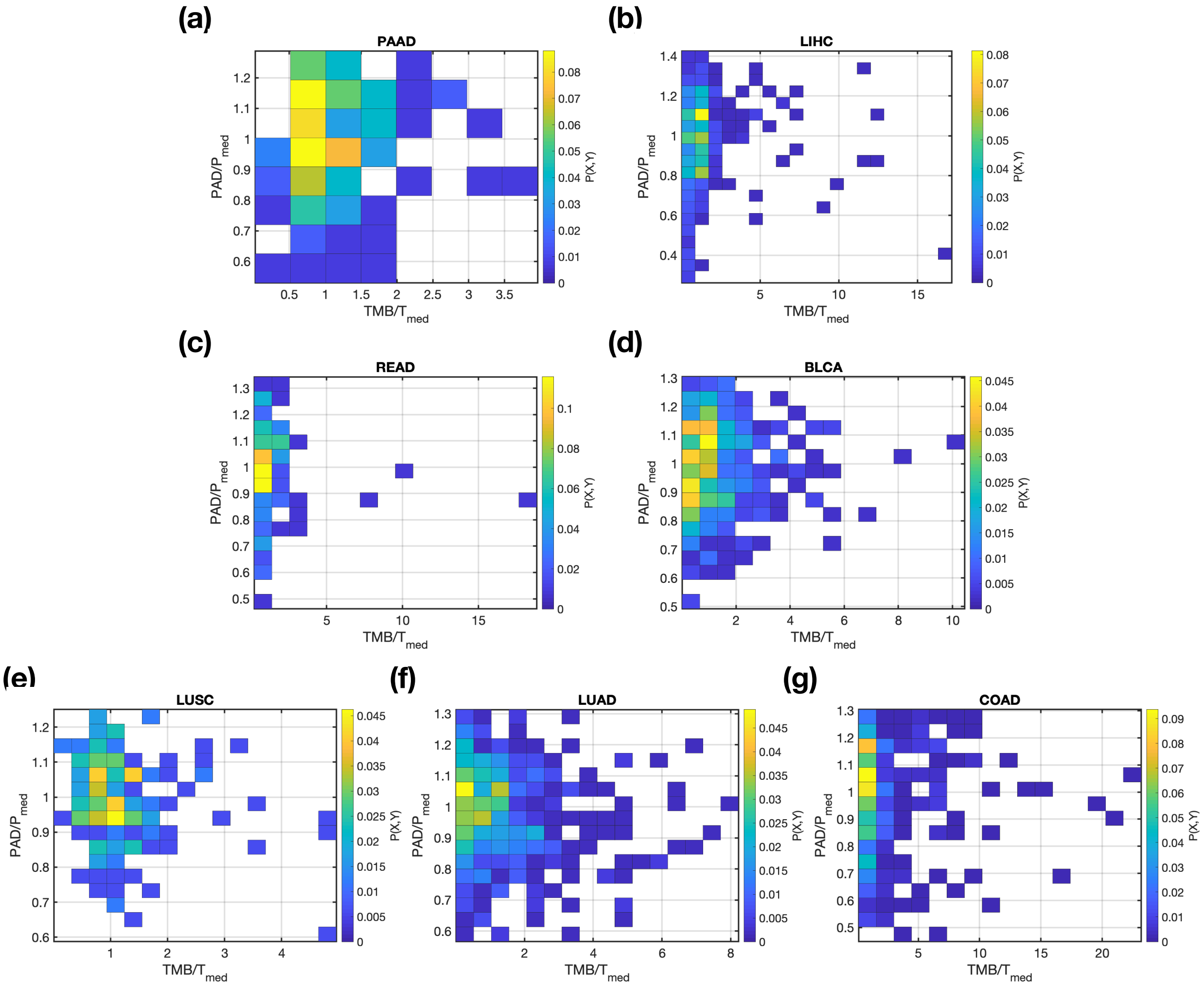

FIG. S13. Same as Fig. S12 except these are for cancers analyzed in Fig. 2.

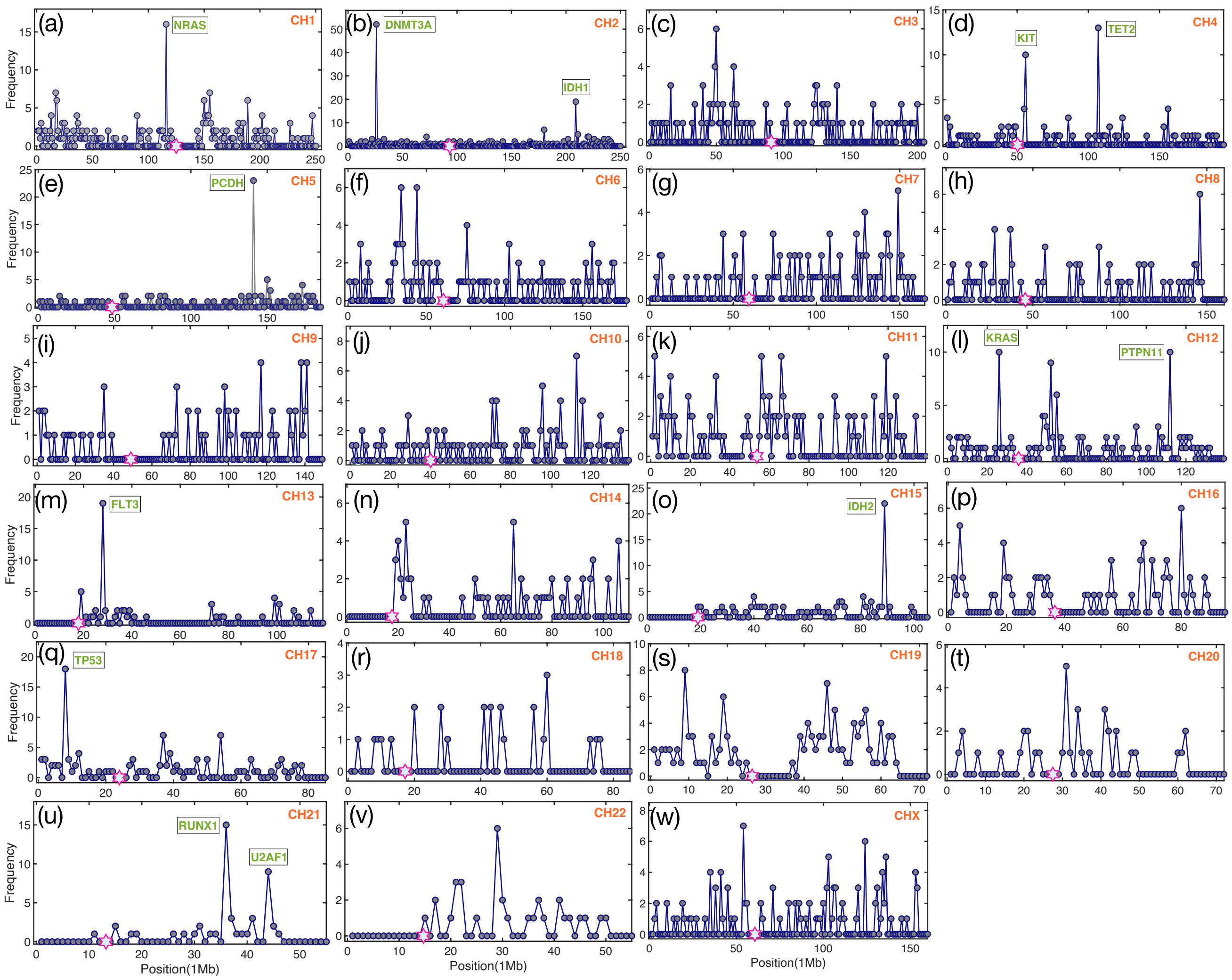

FIG. S14. Tumor mutation frequency (number of mutations in a chromosome region of 1 Mb length) along each chromosome for the LAML. The genes (labelled in green), mutated at a high frequency, are listed in the figure. The hexagram in magenta shows the centromere location on each chromosome. The total number of patients is 197, taken from the Firehose pipeline (http://firebrowse.org).

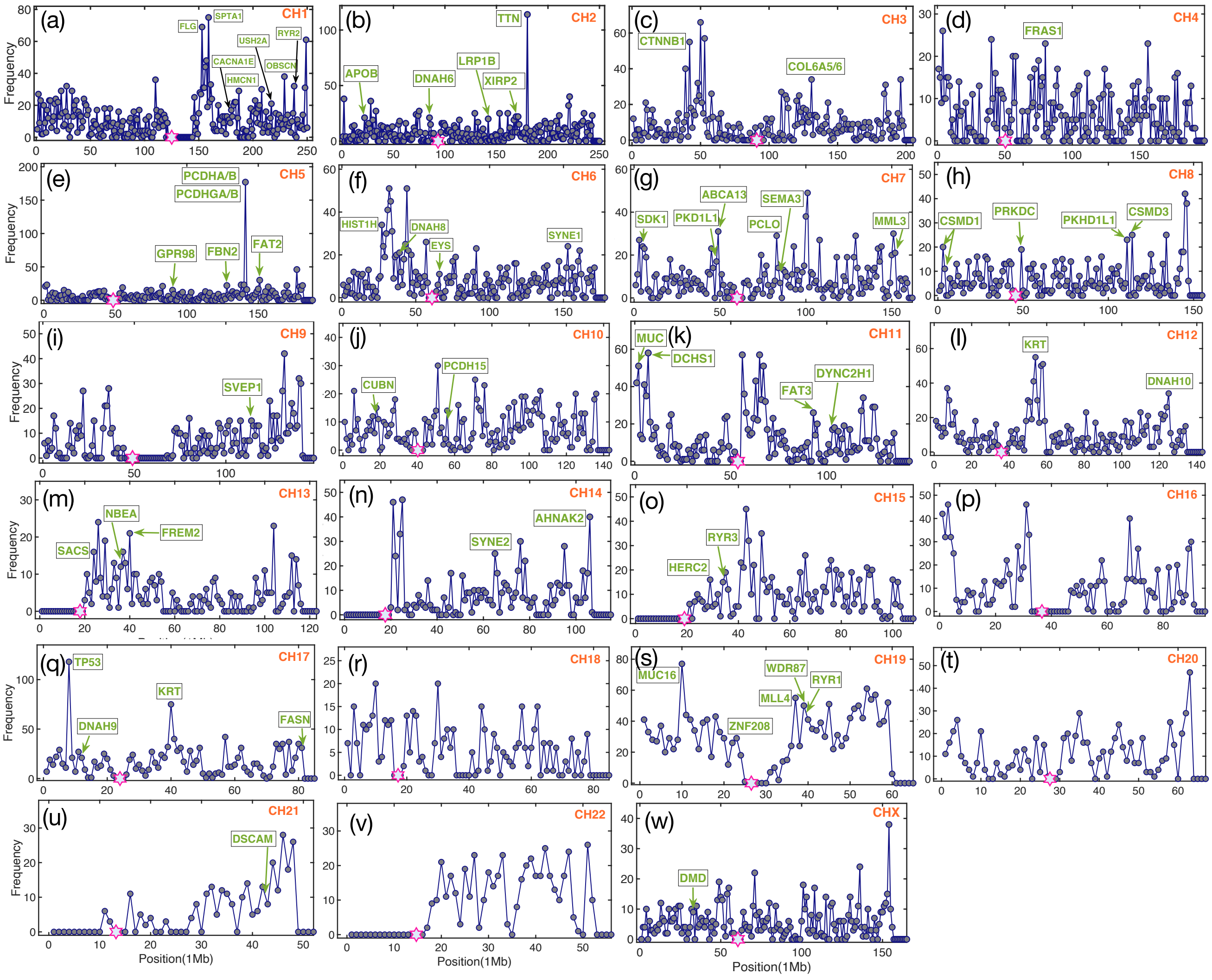

FIG. S15. Same as Fig. S14 except these are for the LIHC cancer. The total number of patients used here is 198, taken from the Firehose pipeline (http://firebrowse.org).

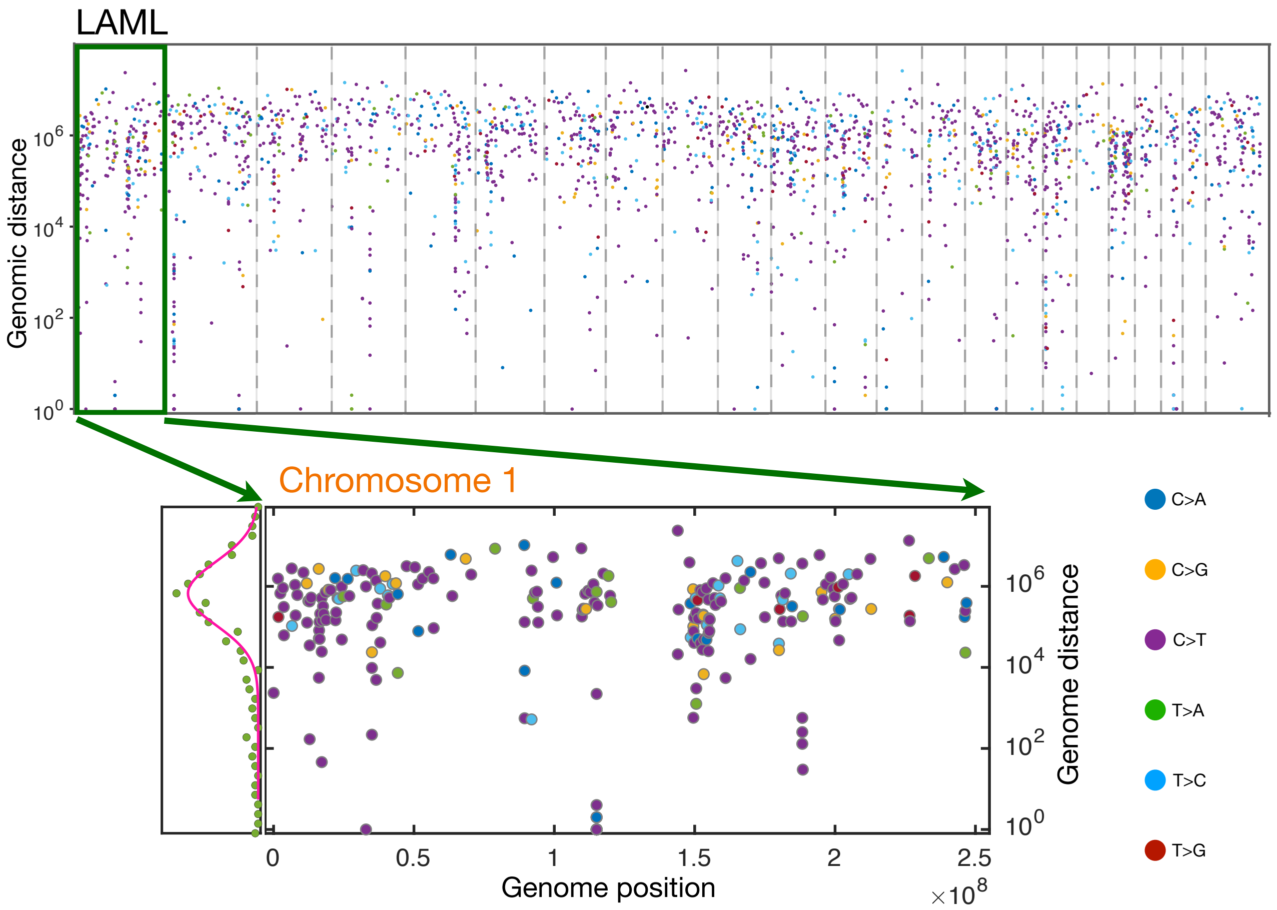

FIG. S16. The upper panel shows the rainfall plot for the low TMB LAML (see also Fig. 5e in the main text). A zoom-in of the green rectangle region (chromosome 1) is shown in the lower panel. The distribution of genomic distance between two successive mutations in chromosome 1 is also shown on the left of the lower panel. The solid line, a Gaussian-function, provides an excellent fit to the data. The unit for genome position/distance is in basepair. The colors of the dots represent six different classes of base substitution, as demonstrated at the right of the figure. No hypermutation region is observed in this figure. The complete absence of SNP mutations in this chromosome close to the centromere region (around 125Mb) is striking.

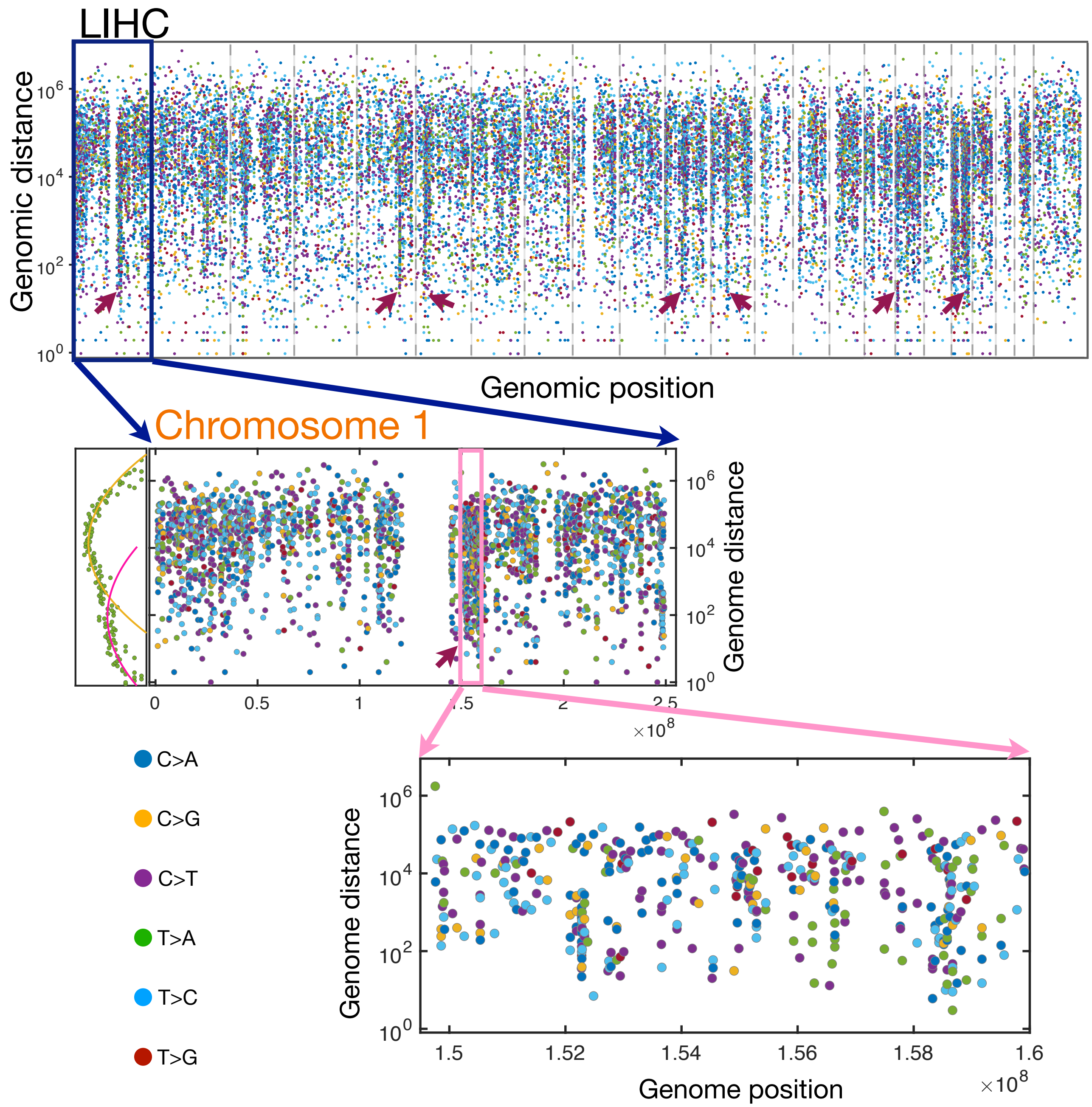

FIG. S17. The upper panel shows the rainfall plot for LIHC (see also Fig. 5f in the main text). Just as in Fig. S16, we illustrate the rainfall plot for chromosome 1 only in the middle panel. The distribution of genomic distance is also shown on the left which can only be described by two Gaussian-functions (see the solid lines) instead of a single one used in Fig. S16. A zoom-in of the hypermutation region (see the pink rectangle) in chromosome 1 is demonstrated in the bottom panel. The red arrowheads indicate hypermutation regions. Comparison of the data in this figure and Fig. S16 illustrates stark differences in the density of mutations.

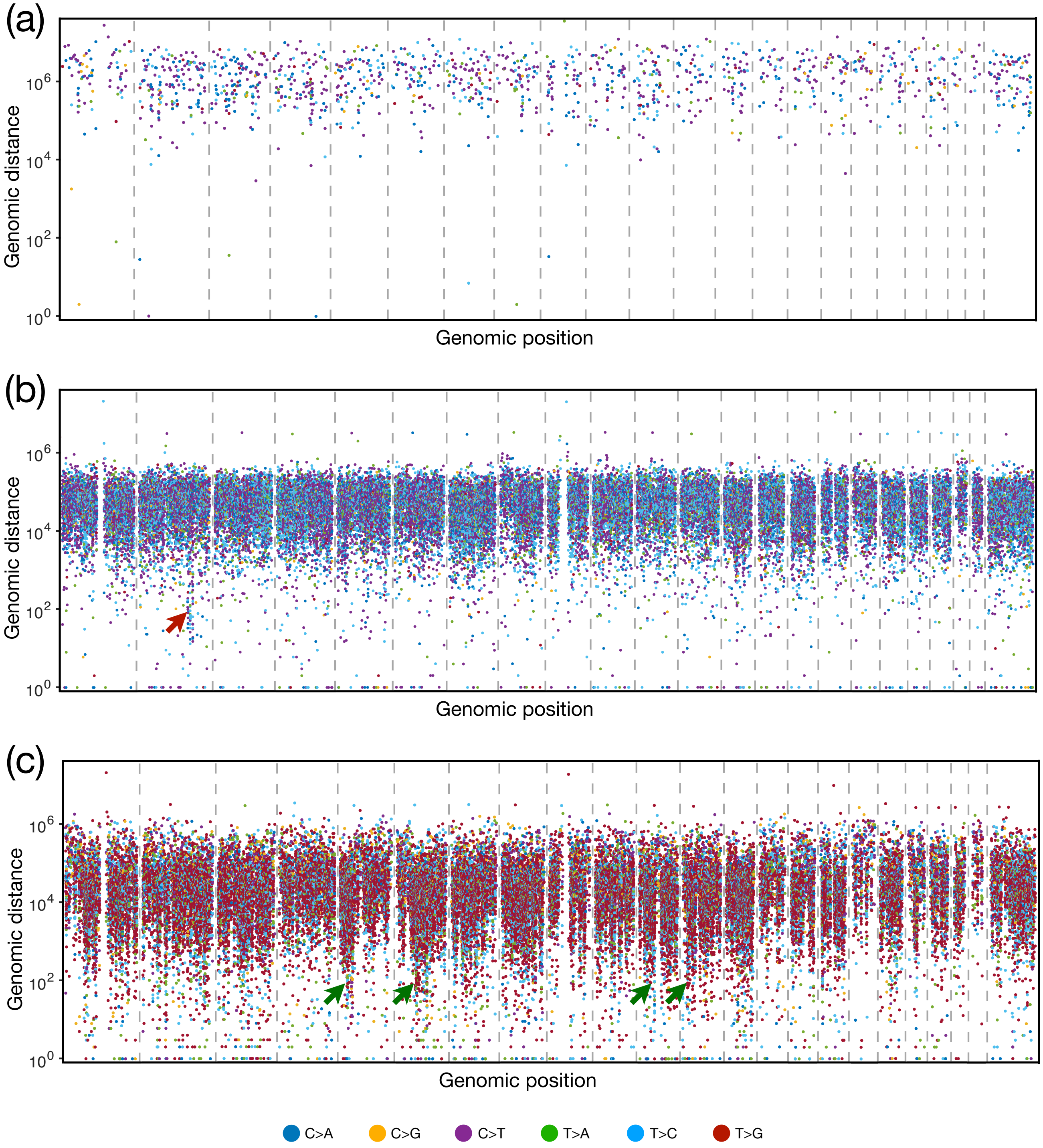

FIG. S18. Kataegis patterns for the whole genome of (a) LAML, (b) LIHC, (c) ESAD cancer patients. The somatic mutations represented by small dots are ordered on the x axis in line with their positions in the human genome. The vertical value for each mutation is given by the genomic distance from the previous mutation. The colors of the dots represent different classes of base substitution, as demonstrated at the bottom of the figure. Several hypermutation regions are indicated by the red/green arrowheads. The data are taken from the reference[7].

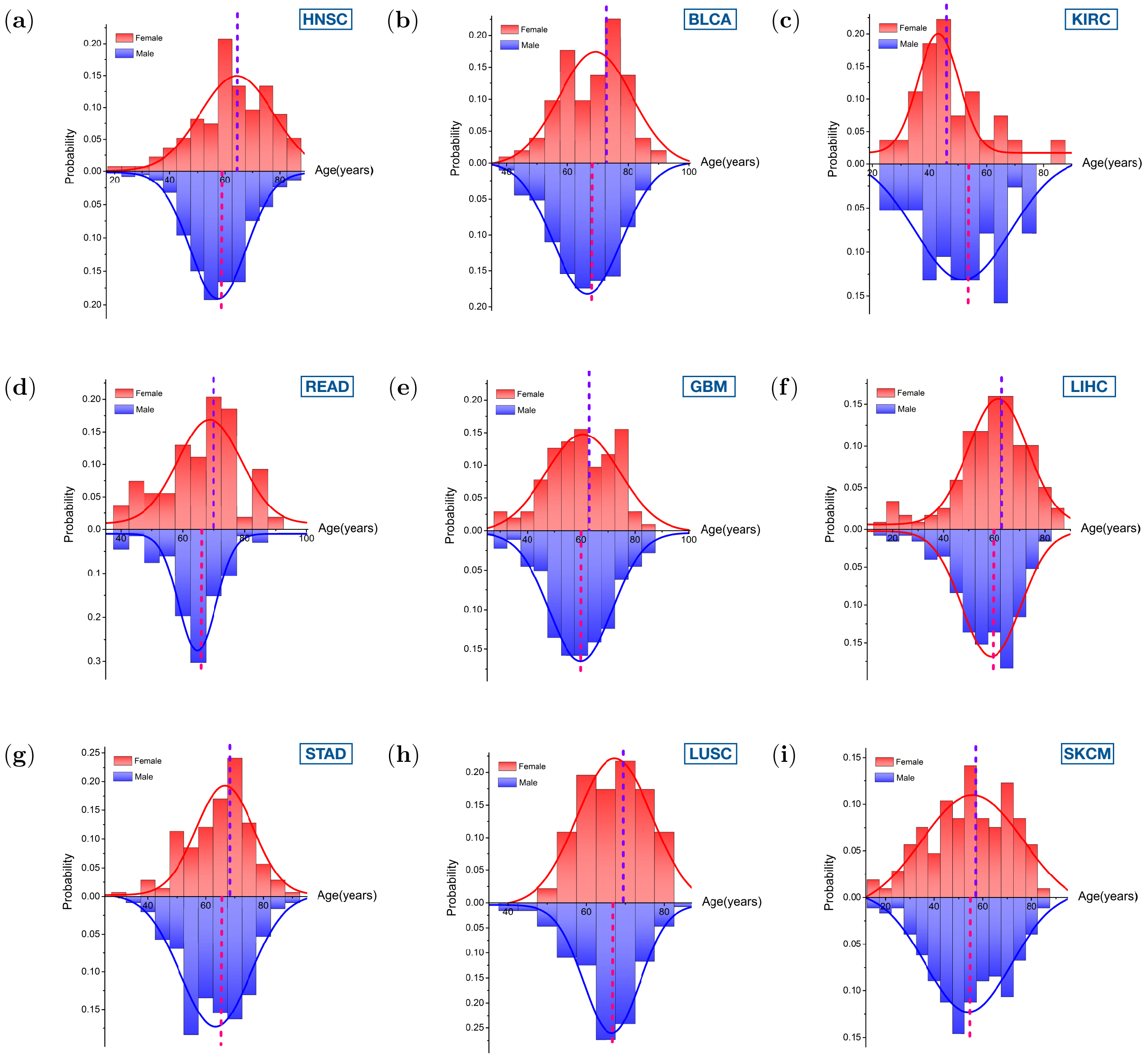

FIG. S19. The patient age distribution (PAD) for both the sexes (red for females and blue for males) across the nine cancer types. The solid lines show the Gaussian-function fit of the data. The dashed lines indicate the median age of cancer patients. The median age for female is larger than that for male in this group of cancers. The median ages for female and male patients at diagnosis are listed in Table VI.

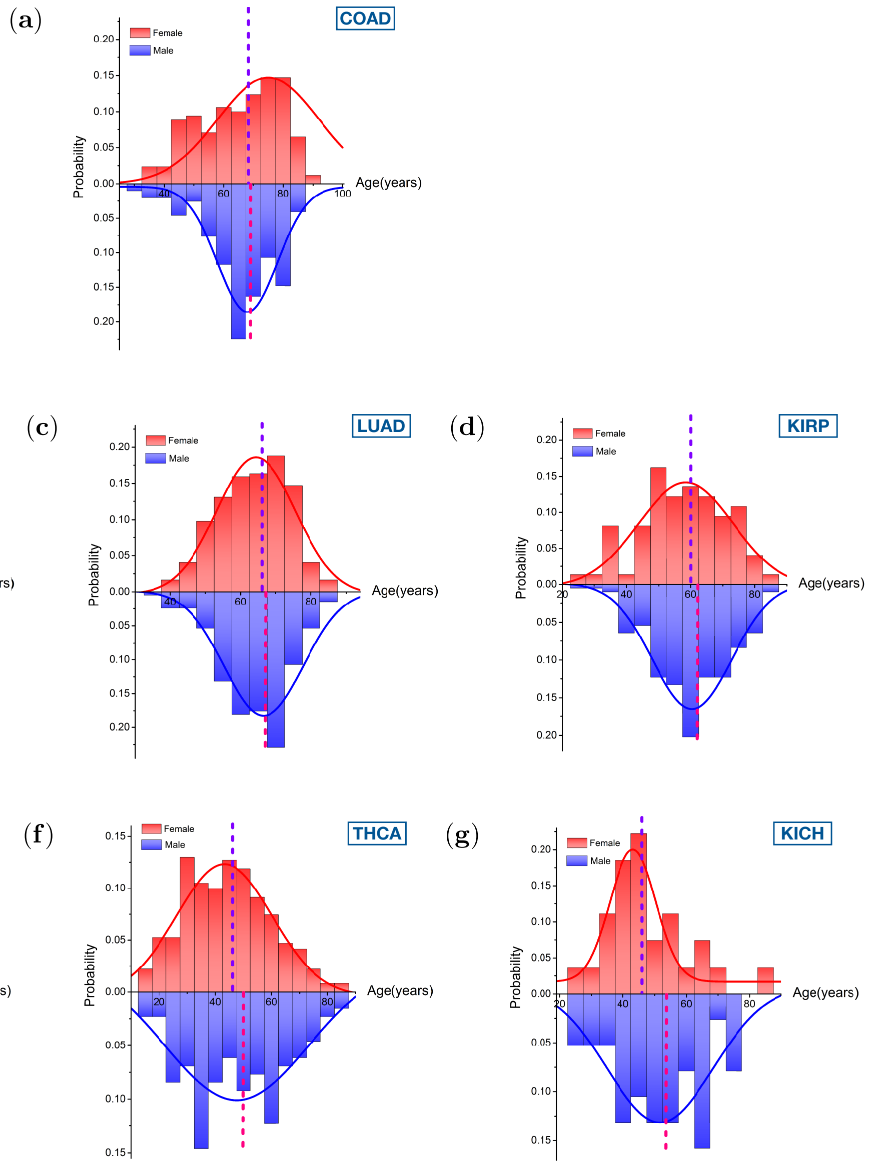

FIG. S20. Same as Fig. S19, except we plot the data for different cancer types. Different from Fig. S19, the median male age is typically larger than for females.

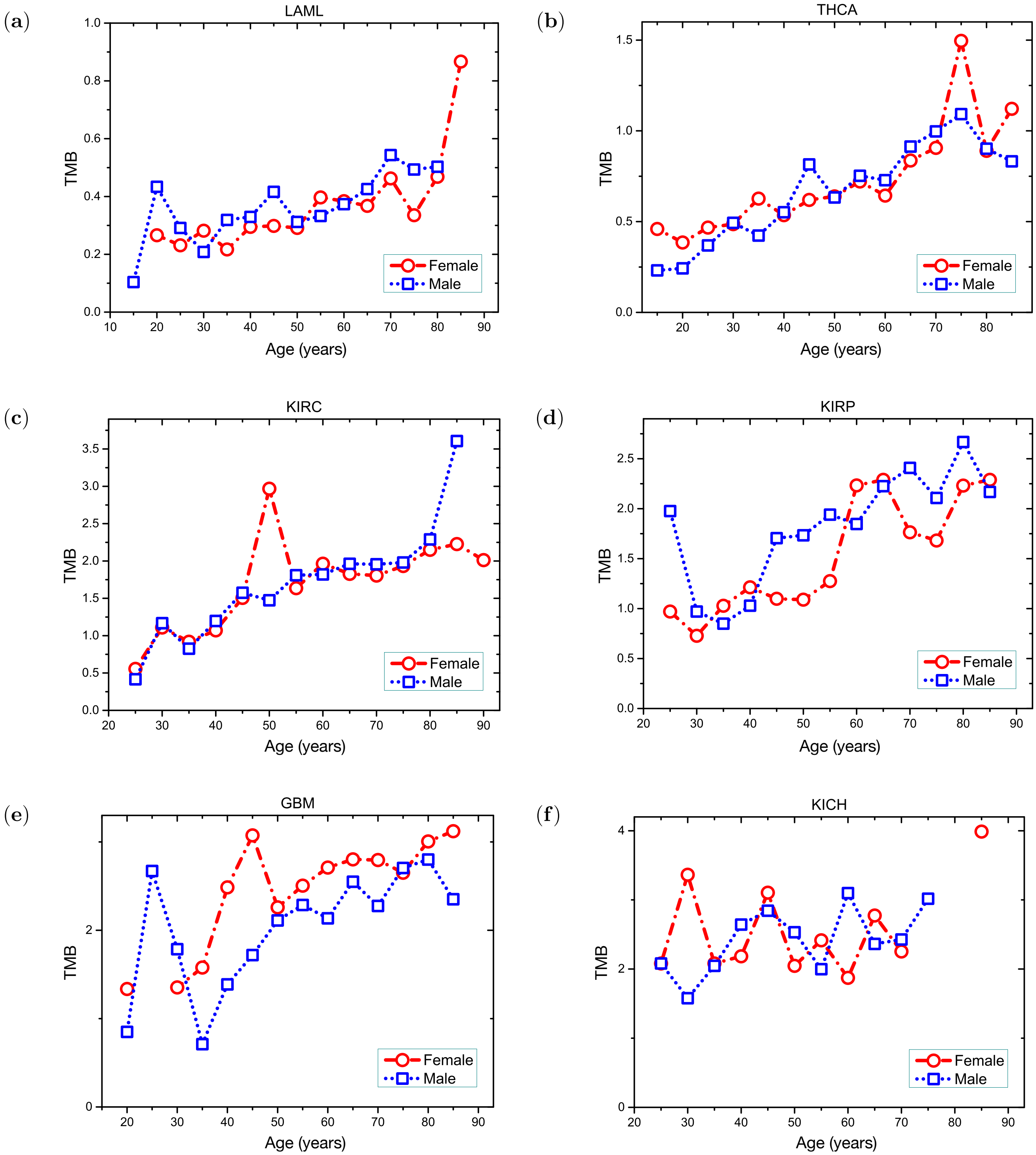

FIG. S21. The tumor mutation burden as a function of age for both the sexes with low overall TMB. The mean value for the mutation burden is used for each 5-year period.

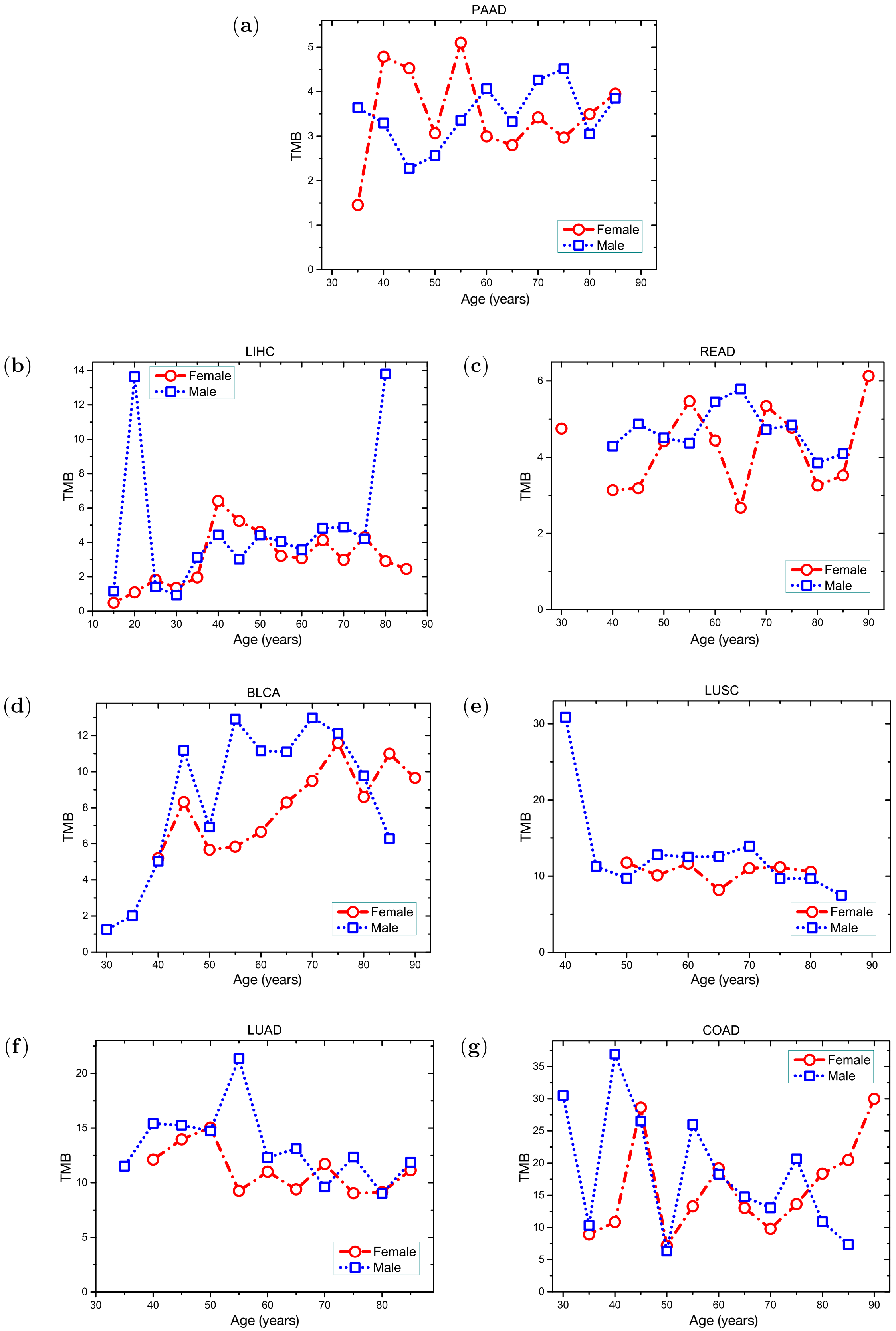

FIG. S22. Same as Fig. S21, except we plot the data for cancer types with high overall TMB.

(a)

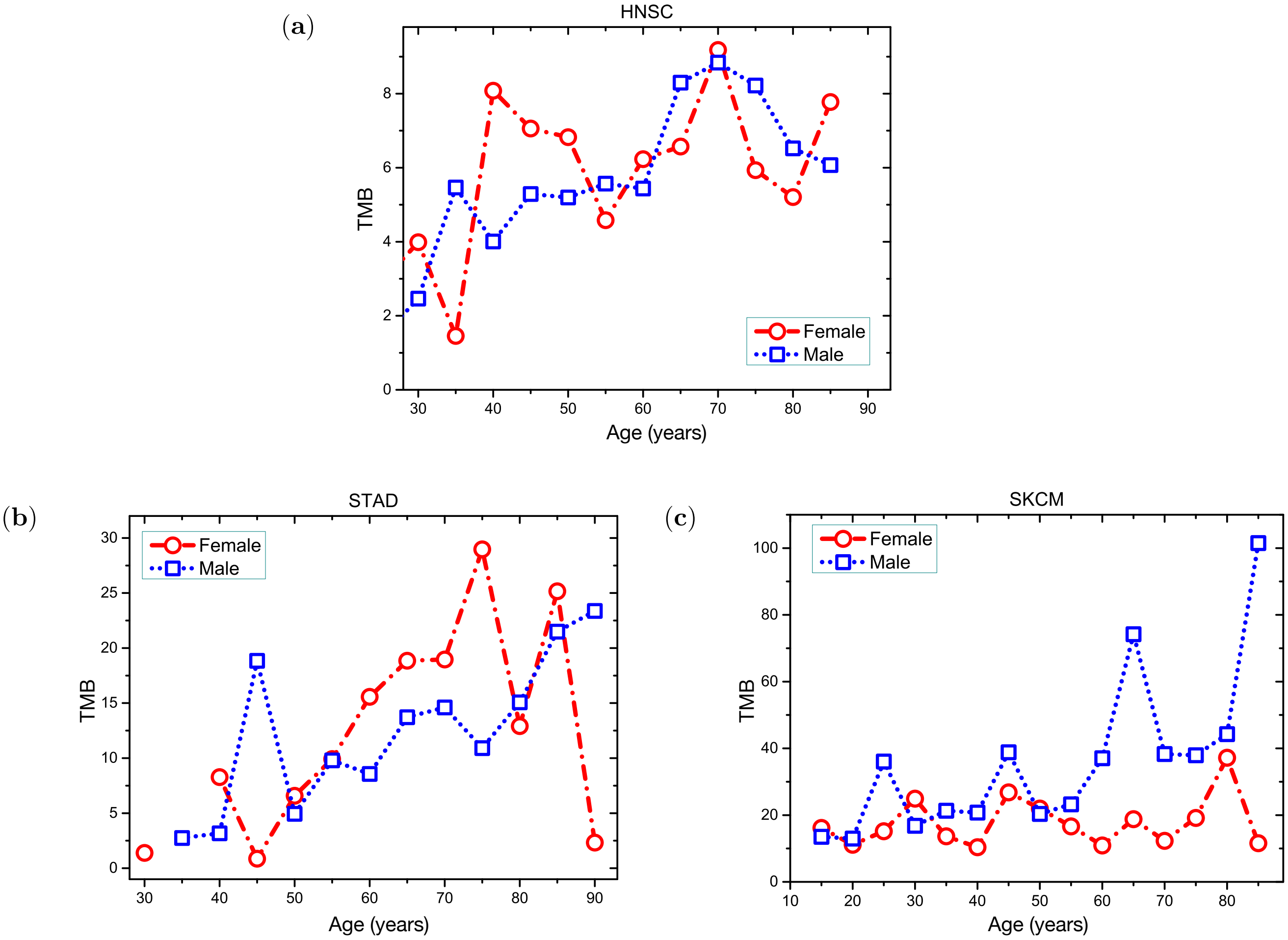

(b) (c)

FIG. S23. The tumor mutation burden as a function of age for both sexes at high overall mutation burden with strong environmental influence.

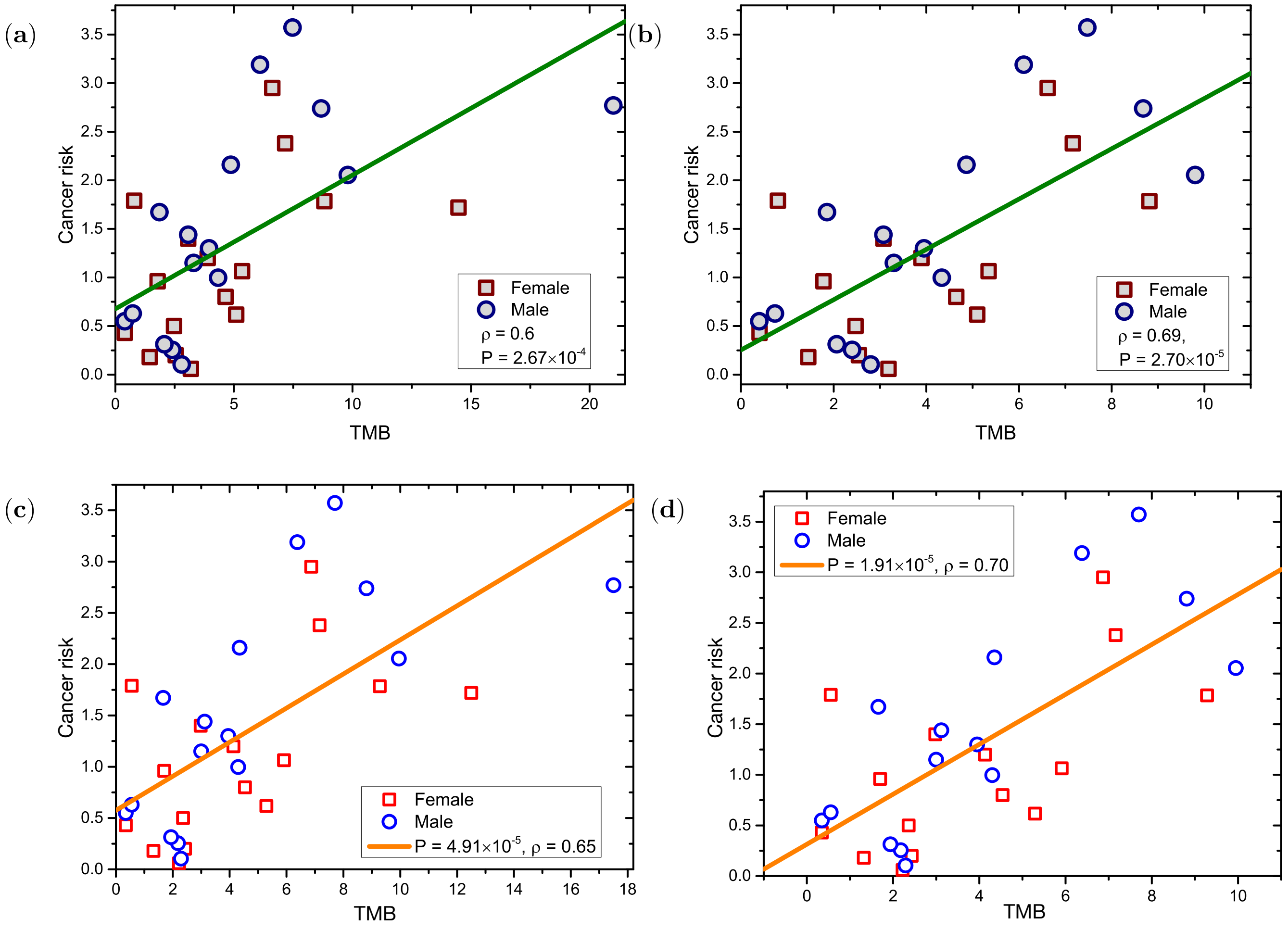

FIG. S24. Cancer risk as a function of the TMB across (a), (c) 16 types or (b), (d) 15 types (excluding SKCM) of cancer for both the sexes. The median value for mutations (sum of non-synonymous and synonymous mutations) per megabase (Mb) is used. The solid line gives the regression linear. The P value from an F test, and the Pearson correlation coefficient $\rho$ for each cancer type are also shown in the figure. The age-adjusted TMB is used in (a), and (b) but not in (c) and (d).

(a)

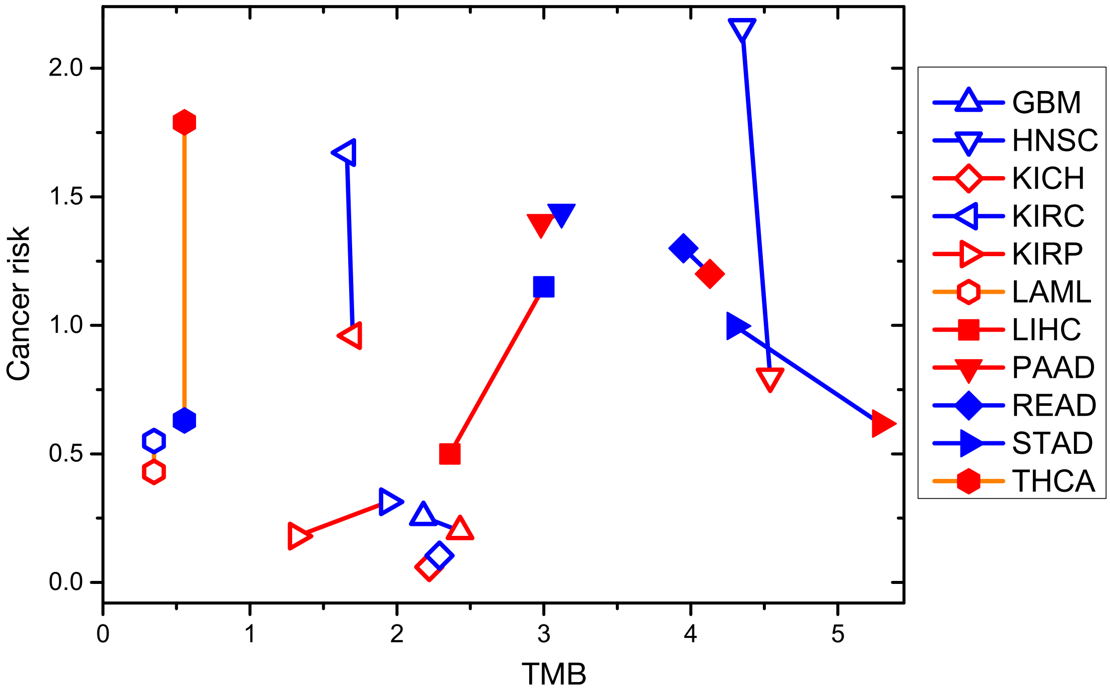

(b)

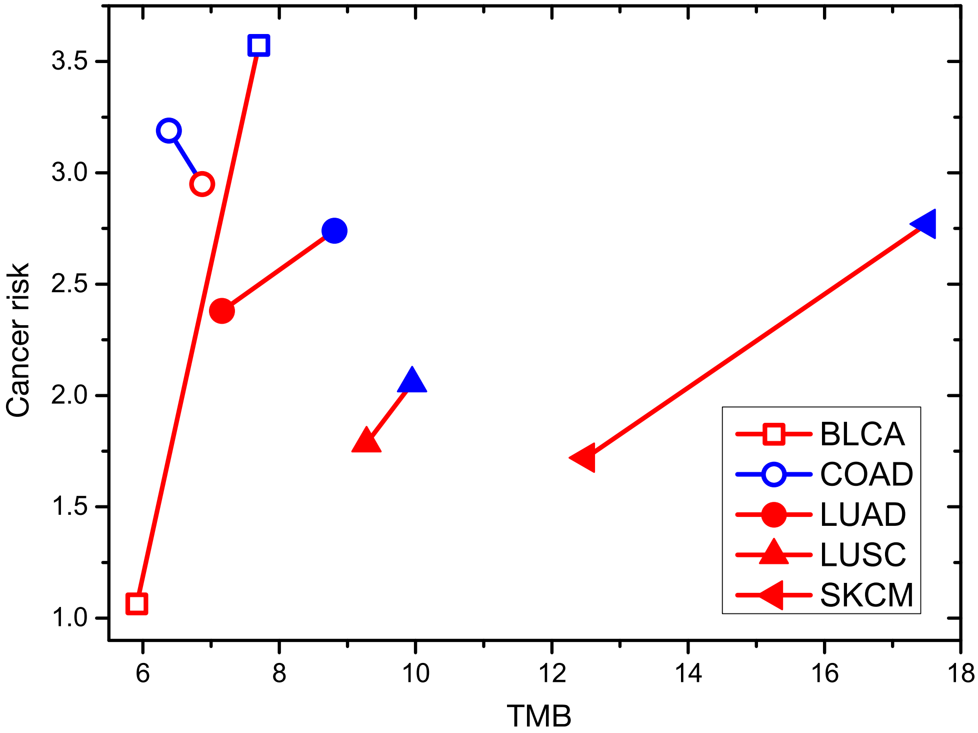

FIG. S25. The relationship between cancer risk and mutation burden for both the sexes across 16 types of cancer. The data are the same as in Fig. S24 while the solid lines connect data for both sexes under the same type of cancer. The data are separated into two groups for visualization purpose. Red symbols show the data for females and the data for males are illustrated in blue.

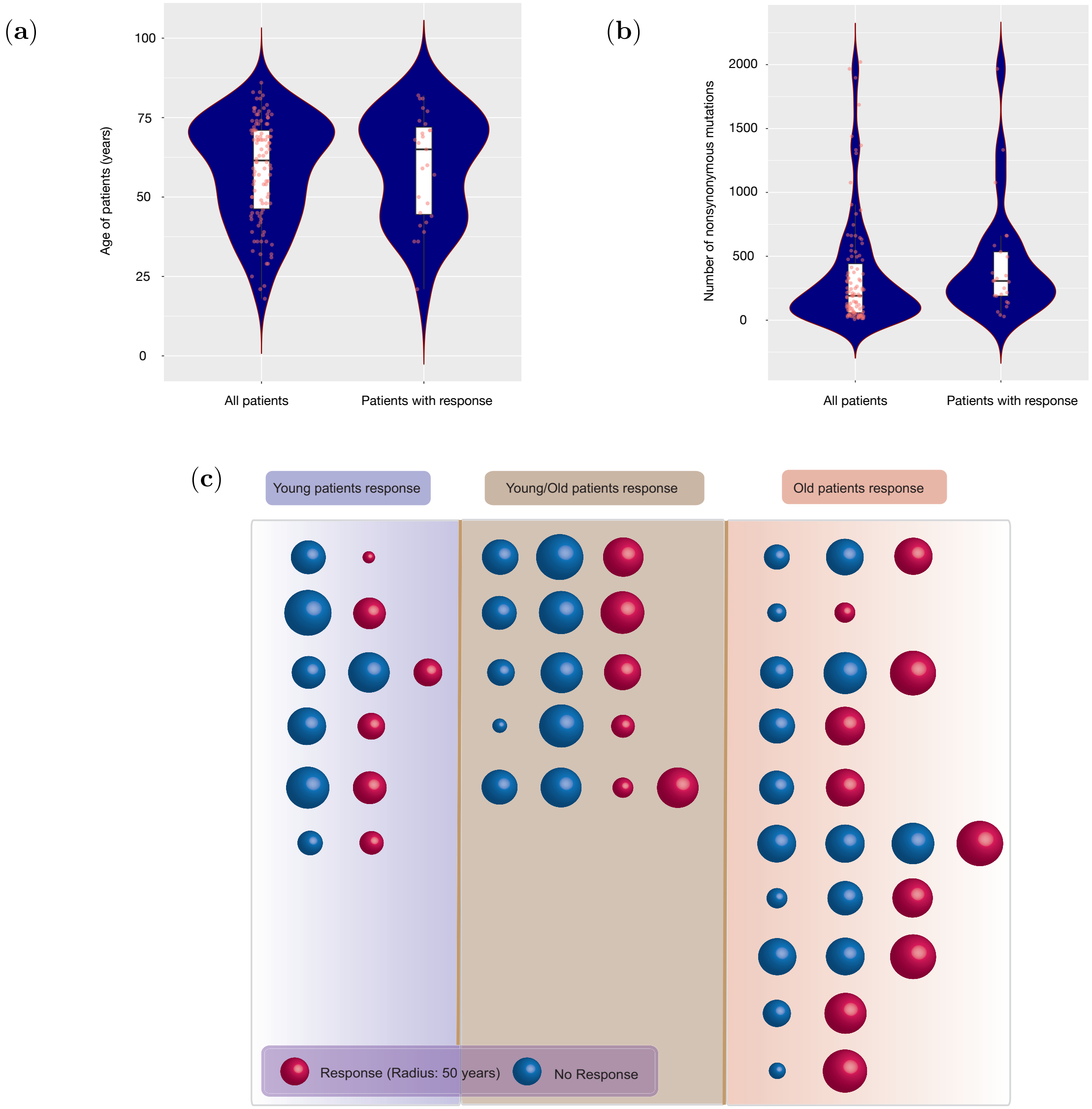

FIG. S26. (a) Violin plots for the age distribution of all patients and the patients showing response to the cytotoxic T-lymphocyte-associated antigen 4 (CTLA-4) blockade[19]. (b) Violin plots for the number of non-synonymous mutations for all patients and the patients responding to CTLA-4 blockade. Five data points are not shown in (b) because they are outliers. A box plot is overlaid on each violin plot, and all the data are shown in red dots in the same figure. (c) Schematic of the dependence of response of melanoma patients of differing ages to immunotherapy. The differences in values of the TMB across the rows is within 5%. Red sphere indicates patients who show response to treatment and patients who do not show response are illustrated by blue spheres. The size of the sphere is scaled with patients age as indicated in the lower left corner. The left (right) column gives all pairs in which young (old) patients respond to the treatment. The middle column shows all mixed cases where either young or old patients respond to the treatment.

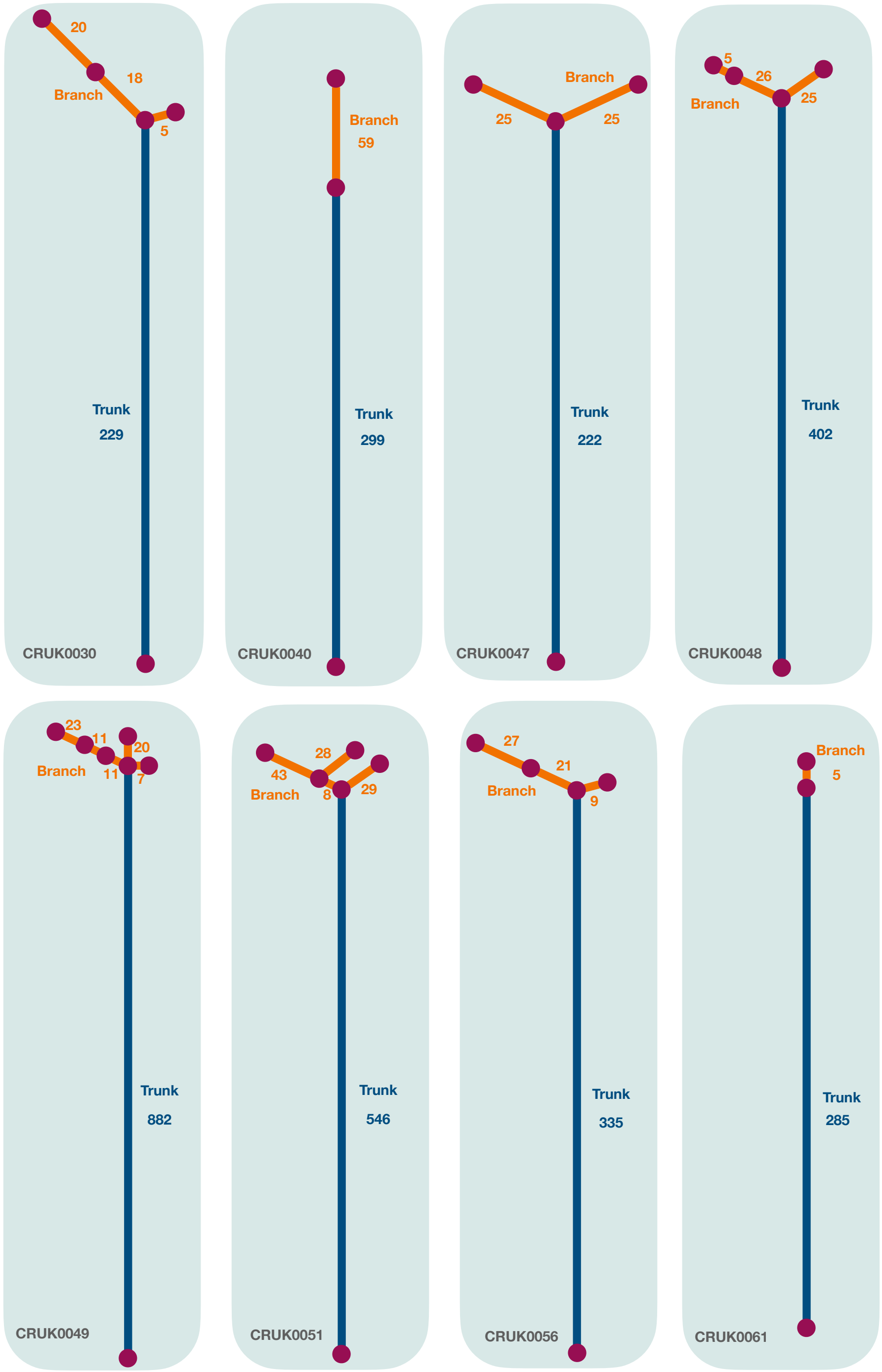

FIG. S27. The phylogenetic trees (revised from Ref.[23]) for cancer patients with high cTMB and low ITH discussed in figure 8(e) in the main text. The length of the trunk (branches) is proportional to the number of clonal (subclonal) mutations found in the patient which is also listed in the figure accordingly. The trunk of the phylogenetic trees is in navy and the branches are in orange color.

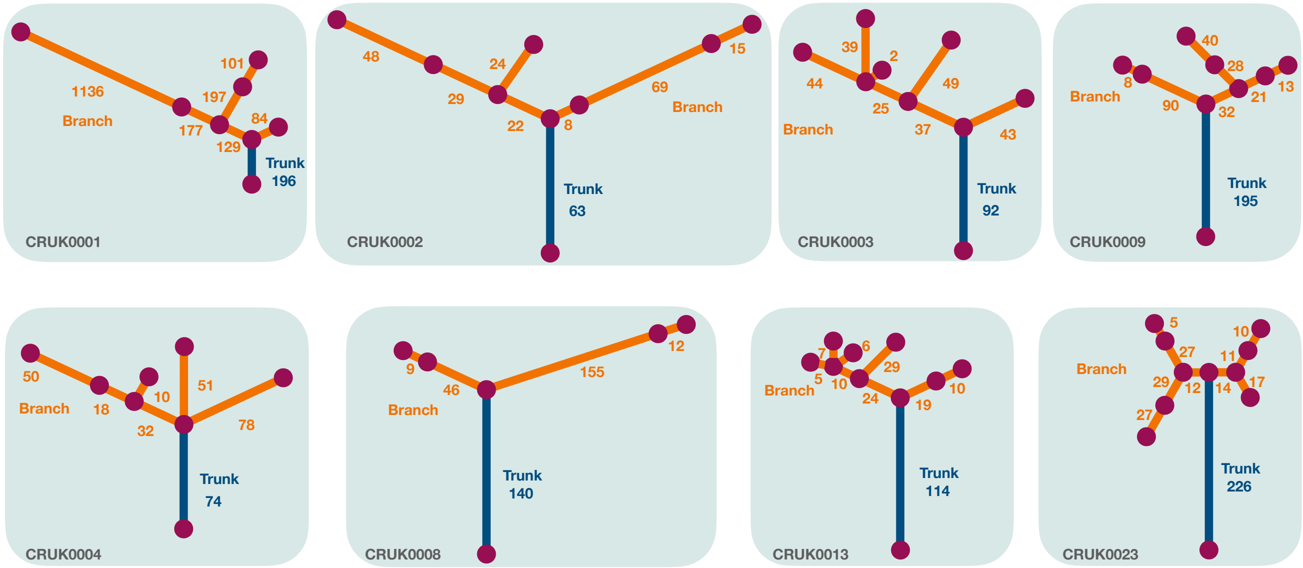

FIG. S28. Same as Figure S27 but for the cancer patients with low cTMB and high ITH discussed in figure 8(e) in the main text.

TABLE I. The number of patients for the 16 types of cancer taken from the TCGA database[6]. Cancer name abbreviations: BLCA – Bladder Urothelial Carcinoma; COAD – Colon Adenocarcinoma; GBM – Glioblastoma multiforme; HNSC – Head and Neck Squamous Cell Carcinoma; KICH – Kidney Chromophobe; KIRC – Kidney Renal Clear Cell Carcinoma; KIRP – Kidney Renal Papillary Cell Carcinoma; LAML – Acute Myeloid Leukemia; LIHC – Liver Hepatocellular Carcinoma; LUAD – Lung Adenocarcinoma; LUSC – Lung Squamous Cell Carcinoma; PAAD – Pancreatic Adenocarcinoma; READ – Rectum Adenocarcinoma; SKCM – Cutaneous Skin Melanoma; STAD – Stomach Adenocarcinoma; THCA – Thyroid Carcinoma.

| **Cancer types** | **Number of patients** |
|---|---|
| BLCA | 394 |
| COAD | 366 |
| GBM | 280 |
| HNSC | 509 |
| KICH | 65 |
| KIRC | 431 |
| KIRP | 277 |
| LAML | 188 |
| LIHC | 369 |
| LUAD | 450 |
| LUSC | 174 |
| PAAD | 125 |
| READ | 120 |
| SKCM | 284 |
| STAD | 386 |
| THCA | 492 |

TABLE II. The latency period and fraction ($F_{ini}$) of accumulated mutations before the initiation of tumors in Figs. 1 and 3 in the main text.

| **Cancer types** | **Latency period (Years)** | **Fraction of mutations $F_{ini}$ (%)** |
|---|---|---|
| LAML | 5-25[36,37] | 71-94 |
| THCA | 5-20[38] | 71-93 |
| KIRC | 10-30[39,40] | 71-90 |
| KIRP | 10-30[39,40] | 60-87 |
| GBM | 10-20[37,41] | 84-92 |
| KICH | 10-30[39,40] | 80-93 |
| HNSC | 12-35[37,42] | 49-82 |
| STAD | 15-22[37,43] | 54-69 |
| SKCM | 5-20[44,45] | 67-92 |

TABLE III. The upper bound for the fraction ($F_{ini}$) of mutations that accumulate before the initiation of tumors in Fig. 2 in the main text.

| **Cancer types** | **Fraction of mutations $F_{ini}$ (%)** |
|---|---|
| PAAD | 34 |
| LIHC | 25 |
| READ | 22 |
| BLCA | 9 |
| LUSC | 10 |
| LUAD | 10 |
| COAD | 10 |

TABLE IV. Mutation burden (median value) for both females and males across the 16 types of cancer from TCGA database[6].

| **Cancer types** | **Mutations/Mbp (female)** | **Mutations/Mbp (male)** |
|---|---|---|
| BLCA | 5.91 | 7.7 |
| COAD | 6.87 | 6.38 |
| GBM | 2.43 | 2.18 |
| HNSC | 4.54 | 4.35 |
| KICH | 2.22 | 2.29 |
| KIRC | 1.70 | 1.66 |
| KIRP | 1.32 | 1.94 |
| LAML | 0.347 | 0.347 |
| LIHC | 2.36 | 3.0 |
| LUAD | 7.16 | 8.81 |
| LUSC | 9.28 | 9.95 |
| PAAD | 2.98 | 3.12 |
| READ | 4.13 | 3.95 |
| SKCM | 12.5 | 17.5 |
| STAD | 5.29 | 4.3 |
| THCA | 0.555 | 0.555 |

TABLE V. Cancer risks (%) for both female and male populations across the 16 types of cancer (see the analysis above in the Supplementary Information).

| **Cancer types** | **Risk (female)** | **Risk (male)** |
|---|---|---|
| BLCA | 1.06 | 3.57 |
| COAD | 2.95 | 3.19 |
| GBM | 0.20 | 0.255 |
| HNSC | 0.80 | 2.16 |
| KICH | 0.06 | 0.105 |
| KIRC | 0.96 | 1.67 |
| KIRP | 0.18 | 0.314 |
| LAML | 0.43 | 0.55 |
| LIHC | 0.50 | 1.15 |
| LUAD | 2.38 | 2.74 |
| LUSC | 1.79 | 2.06 |
| PAAD | 1.40 | 1.44 |
| READ | 1.20 | 1.30 |
| SKCM | 1.72 | 2.77 |
| STAD | 0.618 | 0.998 |
| THCA | 1.79 | 0.63 |

TABLE VI. The median age (years) for female and male patients at diagnosis across the 16 types of cancer[6].

| **Cancer types** | **Median age (female)** | **Median age (male)** |
|---|---|---|
| BLCA | 73 | 68 |
| COAD | 68.5 | 69 |
| GBM | 63 | 60 |
| HNSC | 64.5 | 59 |
| KICH | 46 | 54 |
| KIRC | 63 | 59 |
| KIRP | 60 | 62 |
| LAML | 57 | 58 |
| LIHC | 63 | 60 |
| LUAD | 66 | 67 |
| LUSC | 69.5 | 67 |
| PAAD | 64 | 67 |
| READ | 70 | 66 |
| SKCM | 57 | 55 |
| STAD | 68.5 | 65.5 |
| THCA | 46 | 50 |

TABLE VII. Correlation between TMB of SNVs and PAD for 24 cancer types from WGS data[7]. Two different methods (The Pearson and Spearman correlations) are used to calculate the correlations. The correlation coefficient, P-value and the number of patients for each cancer type are listed in the table. Five of the 24 cancer types from different countries are also calculated separately and listed at the end of the table.

| Cancer Name | Pearson correlation coefficient | P-value | Spearman's rho | P-value | Number of samples |
|---|---|---|---|---|---|
| CNS_PILO | 0.46 | 5.75E-06 | 0.5618 | 1.01E-08 | 89 |
| KICH | 0.5087 | 5.82E-04 | 0.4841 | 0.001 | 42 |
| THCA | 0.5693 | 2.41E-05 | 0.7027 | 2.59E-08 | 48 |
| CNS_MEDU | 0.7573 | 8.13E-26 | 0.7679 | 6.42E-27 | 132 |
| CLL | 0.3294 | 0.0015 | 0.3684 | 3.53E-04 | 90 |
| KIRC | 0.4395 | 0.0065 | 0.4543 | 0.0047 | 37 |
| PRAD | 0.2084 | 0.0289 | 0.2177 | 0.0224 | 110 |
| KIRP | 0.5801 | 5.01E-04 | 0.5956 | 3.22E-04 | 32 |
| BONE-OST | 0.335 | 0.0346 | 0.1918 | 0.2359 | 40 |
| GBM | 0.4672 | 0.0031 | 0.4511 | 0.0045 | 38 |
| BRCA | 0.3521 | 0.0097 | 0.3177 | 0.0204 | 53 |
| OVCA | -0.0205 | 0.8673 | 0.0198 | 0.8719 | 69 |
| PAAD | 0.0498 | 0.5605 | 0.0014 | 0.9867 | 139 |
| LIHC | -0.0783 | 0.5772 | -0.1041 | 0.4582 | 53 |
| COAD | -0.0792 | 0.6888 | -0.1077 | 0.5855 | 28 |
| UCEC | 0.0592 | 0.7133 | 0.4297 | 0.005 | 41 |
| BLCA | 0.1666 | 0.4474 | -0.0064 | 0.9768 | 23 |
| STAD | 0.1149 | 0.5244 | 0.0766 | 0.6718 | 33 |
| HSCC | 0.2522 | 0.0608 | 0.3054 | 0.0221 | 56 |
| BNHL | 0.0023 | 0.9819 | -0.064 | 0.5273 | 100 |
| ESAD | 0.0307 | 0.7653 | 0.0138 | 0.8931 | 97 |
| LUSC | -0.1594 | 0.2844 | -0.2214 | 0.1347 | 47 |
| LUAD | -0.1958 | 0.2671 | -0.0781 | 0.6608 | 34 |
| MELA | 0.0407 | 0.7382 | 0.0546 | 0.6537 | 70 |
| RECA | 0.1148 | 0.3302 | 0.3792 | 8.64E-04 | 74 |
| LINC | 0.0696 | 0.2673 | 0.0308 | 0.6234 | 256 |
| HNSC | 0.2209 | 0.1546 | 0.2773 | 0.0718 | 43 |
| SKCM | 0.0295 | 0.8625 | 0.0774 | 0.6489 | 37 |
| OVUS | 0.0828 | 0.6069 | -0.0521 | 0.7465 | 41 |

Cancer name abbreviations: CNS_PILO, idney Chromophobe; THCA, thyroid Carcinoma; CNS_MEDU, kidney renal clear cell carcinoma; PRAD, kidney renal papillary cell carcinoma; BONE-OST, bone osteosarcoma; GBM, glioblastoma multiforme; BRCA, breast invasive ductal carcinoma; OVCA, ovary adenocarcinoma (AU); PAAD, pancreatic adenocarcinoma; LIHC, liver hepatocellular carcinoma (US); COAD, colon adenocarcinoma; UCEC, uterus adenocarcinoma; BLCA, bladder urothelial carcinoma; STAD, stomach adenocarcinoma; HSCC, head-and-neck squamous cell carcinoma; BNHL, lymphoid mature B-cell lymphoma; ESAD, esophagus adenocarcinoma; LUSC, lung squamous cell carcinoma; LUAD, lung adenocarcinoma; MELA, skin melanoma (AU); RECA, kidney renal clear cell carcinoma (EU); LINC, liver hepatocellular carcinoma (JP); HNSC, head and neck squamous cell carcinoma (US); SKCM, cutaneous skin melanoma (US); OVAU, ovary adenocarcinoma (US).
